# Genomics-based quantitative biogeography of marine plankton

**DOI:** 10.1101/2025.10.25.684415

**Authors:** Margaux Crédeville, Roy El Hourany, Swan L. S. Sow, Julie Poulain, Manon Depaty, Eric Pelletier, Zoé Mériguet, Marie-Fanny Racault, Genoscope Sequencing Team, Aude Perdereau, Laurie Bertrand, Frédérick Gavory, Priscillia Gourvil, Céline Orvain, Morgane Ratin, Laurence Garczarek, Tom O Delmont, Adrien Thurotte, Corinne Le Quéré, Juan J Pierella Karlusich, Chris Bowler, Samuel Chaffron, Patrick Wincker, Fabien Lombard, Olivier Jaillon

## Abstract

Marine plankton are key drivers of ocean productivity and global carbon cycling, yet their quantitative biogeography remains poorly characterized. Environmental genomic datasets are inherently compositional, restricting analyses to relative abundances and limiting their integration into ecological and biogeochemical models. Here we combine DNA mass measurements, filtered seawater volumes, and metagenomic relative abundances to generate absolute estimates of cell concentrations and carbon biomass of plankton across the global ocean. Leveraging thousands of samples, this approach provides quantitative estimates for hundreds of eukaryotic and thousands of prokaryotic environmental genomes, unveiling ecological associations not captured by compositional data. Using the *psbO* marker gene, we reconstruct quantitative biogeographies of photosynthetic lineages and, through global-scale modeling, project their distributions to generate quantitative maps of phytoplankton communities across the world’s ocean. By bridging genomic data with biogeochemical metrics, this study provides novel resources for integrating plankton at genomic resolution into next-generation biogeochemical models.

## Introduction

Marine plankton are the foundation of ocean ecosystems and major drivers of global biogeochemical cycles, sustaining primary production and modulating the flux of carbon between the atmosphere and the ocean (*1*, *2*). The biomass of marine plankton is unevenly distributed across the global ocean, reflecting ecological and biogeochemical gradients. Global-scale surveys and representations that combine in situ observations, satellite data, and ecosystem models have shown that phytoplankton biomass peaks in nutrient-rich regions at high latitudes and in upwelling zones, and is lowest in subtropical gyres (*3*, *4*). These variations can span several orders of magnitude between oligotrophic subtropical gyres and productive polar or upwelling regions (*5*, *6*). Within prokaryotic plankton, cyanobacteria of the *Prochlorococcus* genera in particular dominate tropical and subtropical oligotrophic waters, while diatoms and other eukaryotic lineages, including green algae, dominate in polar biomes (*7–9*). At the planetary scale, despite having much lower biomass than terrestrial ecosystems (*10*), plankton collectively sustain a level of primary production comparable to that of land plants (*11*). These large-scale modeled patterns provide an essential framework for interpreting plankton ecology. But quantitative assessments of plankton biomass that integrate genomic resolution and extend across multiple ocean basins are still lacking.

In the last decade, advances in environmental genomics thanks to lowering costs of DNA sequencing have considerably expanded the scope for studying marine plankton communities by offering unparalleled resolution and data magnitude. Omics studies can address various ecological questions regarding the diversity of plankton species, as well as molecular mechanisms underlying their evolutionary adaptation and capacity for acclimation. They also allow for the study of specific genes and functional pathways, providing key insights into the metabolic processes crucial to ocean biogeochemistry (*12*, *13*). However, most omics approaches were not originally developed for quantitative measurements and therefore only provide information about the relative abundance of the genomic entities under study (*14*, *15*). The compositional nature of omics data requires specific methodological considerations (*16*, *17*) because inappropriate statistical approaches or data pre-processing can lead to highly biased results and spurious correlations (*18*, *19*).

Beyond methodological constraints, many ecological questions require quantifying *how much is where,* at a biological scale compatible with functional differentiation. Conventional quantitative approaches, such as imaging or satellite observations, often fail to capture functional differences among closely related lineages, as recently exemplified by surveys about the presence or absence of *nifH*, a gene associated with diazotrophy, in plankton genomes (*20*). Therefore, combining quantitative community descriptions with the high resolution of omics tools is essential for improving our understanding of plankton community biogeography and ecosystem functioning, particularly in the context of climate change. Obtaining quantitative omics data also remains a key challenge for improving biogeochemical modelling approaches (*21*, *22*). Indeed, these models require quantitative information on the abundance and biomass of plankton species or functional traits for parameterisation and validation (*23*, *24*). Other modelling approaches may even integrate genomic resolution directly into their design (*25*, *26*).

Numerous studies have already demonstrated the value of quantitative omic approaches to complement results initially obtained from compositional data of microbial communities (*27–29*). Several categories of pre- and post-sequencing methods have been described to extract quantitative information from omics data (*30*, *31*). Harrison et al. (*31*) reviewed various straightforward “spike-in” approaches that involve the addition of an internal DNA standard prior to sequencing. This standard is used as a reference to estimate the mass or numbers of genomic units, such as transcripts or DNA molecules, in the initial sample material (*32*, *33*). Most recently, Bei et al. (*34*) employed such a workflow targeting the photosynthetic *psbO* marker gene (encoding for the manganese-stabilising polypeptide of the photosystem II oxygen evolving complex) to quantify genome-equivalent concentrations of both eukaryotic and prokaryotic phytoplankton lineages in unfractionated seawater. Notably, for cyanobacteria, this analysis was extended further to provide absolute cell concentration estimates. Documented as a single-copy gene in most prokaryotic and eukaryotic phytoplankton lineages (*9*), *psbO* has consequently been proposed as a potential marker for phytoplankton cell quantification (*34*). For several phytoplankton taxa, Bei et al. (*34*) reported metagenomic-based estimates of cell or genome abundance that were remarkably consistent with flow cytometry and microscopy counts. Despite such highly promising results, spike-in approaches must be planned in advance and integrated very early in the sequencing process. These approaches also require prior estimation of the average total DNA mass in samples to determine the appropriate amount of DNA standard. Large-scale plankton surveys must, by design, cover contrasted oceanographic systems, including both oligotrophic and nutrient-rich waters. In addition, these campaigns also collect plankton across several organism size fractions to capture the full extent of the plankton trophic network, up to meso- and macroplankton. Such field constraints lead to substantial variability in biomass among samples and therefore complicate the standardised implementation of spike-in methods.

Other methods have been developed to quantify omics data by converting relative abundances inferred from sequencing into absolute abundances, reflecting the actual quantities of genomic units or organisms present in the original sample. These post-sequencing approaches rely on calibration against a known external reference whose absolute quantity serves as a standard to scale relative abundance data into absolute values, expressed as number of cells or genomes per mass or volume of sample. For bacterial communities, total cell counts obtained by flow cytometry or qPCR have been most commonly used as absolute reference (*27*, *35*). Pierella Karlusich et al. (*36*) also introduced a DNA-based quantification method to estimate *nifH* gene copy concentrations in seawater by combining the mass of extracted DNA, the relative abundances of *nifH* reference sequences in *Tara* Oceans metagenomes, and gene length.

Here, expanding on this DNA-based quantification rationale, we leverage the mass of DNA extracted from plankton filter samples as an external absolute reference to convert metagenomic relative abundance data into absolute abundances. More specifically, DNA quantities are expressed as DNA concentrations in seawater after retrieving the volumes of seawater filtered during sampling from previous campaigns.

Using samples from the *Tara* Oceans, *Tara* Oceans Polar Circle and *Tara* Pacific expeditions, we integrate *psbO*-based phytoplankton characterization (*9*) with genome-resolved quantification of eukaryotic and prokaryotic lineages (*26*, *37*, *38*), generating a large-scale quantitative resource that provides an integrated view of plankton community structure across geographic and taxonomic dimensions.

Specifically, our genome-based estimates of cell concentration in seawater show the dominance of diazotrophic *Trichodesmium* over non-nitrogen-fixing lineages across *Tara* stations. These estimates also reveal specific ecological associations through co-occurrence networks. At the marker-gene level, this approach enables quantitative analysis of the biogeographical structure of the phytoplankton community, both across *Tara* sampling sites and in global-scale predictive models of quantitative biogeography using machine learning techniques. These quantitative resources also make it possible to revisit the influence of environmental parameters and biotic interaction indicators on plankton communities by considering their quantitative variations across sites, providing a clearer view of the ecological mechanisms that govern their distribution and composition. It also leads to conversion of cell abundances into carbon biomass for major phytoplankton groups, providing a link between genomic, biogeochemical data and their potential use in modelling.

## Results

### Overview of the Quantification Framework

This study builds on samples collected during three *Tara* expeditions—*Tara* Oceans (*39*, *40*), *Tara* Oceans Polar Circle (*39*, *40*), and *Tara* Pacific (*41–43*)— which together surveyed plankton communities across all major oceanic regions. In this work, we focused mainly on the cellular fractions of plankton, from pico-to mesoplankton, encompassing prokaryotes, unicellular eukaryotes, and metazoans. The methodology for retrieving DNA concentrations in seawater and quantifying DNA-based absolute abundance of plankton genomes or cells is fully detailed in the Methods section. Briefly, masses of DNA extracted from plankton filter samples were used to calculate DNA concentrations in seawater after retrieving the exact volumes of seawater filtered during sampling. This quantification approach was made possible by the largely standardized sampling protocols implemented across all three expeditions, including consistent seawater collection and filtration procedures for each size fraction, precise documentation of filtered volumes, as well as uniform processing of genomic samples (*40, 43–45*).

In total, DNA concentrations were estimated for over 1,000 surface samples of plankton encompassing ten size fractions ranging from 0.2 to 2,000 µm (Fig. S1). Smaller size fractions generally exhibited higher DNA concentrations in seawater compared to larger fractions, with median concentrations ranging from 5.5 x 10^5^ ng.m^-3^ for the 0.2-3 µm size fraction to 48 ng.m^-3^ for the 300-2,000 µm size fraction (Fig. S2). Additionally, larger size fractions (20-180 µm, 180-2,000 µm, and >300 µm) showed a greater magnitude of variation in DNA concentration, with fluctuations spanning up to five orders of magnitude, whereas smaller size fractions varied by only one or two orders of magnitude.

Based on the assumption that metagenomic relative abundances of genomes or genes are equivalent to the proportions of their DNA mass in samples (*46–48*), we used the DNA concentration corresponding to a given sample as a quantitative “anchor” (*30*) to convert metagenomic relative abundances into absolute DNA concentrations of genomes or reference sequences in that sample (Fig. S1). When possible, we derived absolute cell abundances in seawater associated with a given gene or genome, using the DNA concentrations and the nucleotidic length of the target. This final step was conducted only if information on ploidy was available or could be reasonably approximated (e.g., 1 for Prochlorococcus and Synechococcus, 2 for diatoms). Thus, the whole quantification procedure for a target *i* (marker-gene or genome) in sample *j* can be summarised as follows:

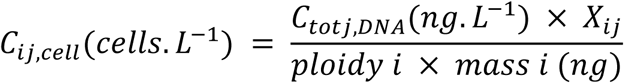

where Xij is the metagenomic relative abundance of target *i* in metagenome *j*.

We first applied this method to estimate DNA concentrations and cell concentrations for more than 5,000 reconstructed and reference genomes of eukaryotes and prokaryotes across all sampled size fractions in *Tara* Oceans and *Tara* Oceans Polar Circle stations (*26*, *37*, *38*). Finally, we used the *psbO* reference sequence database from Pierella Karlusich et al. (*9*) to quantify DNA concentrations and associated gene copy concentrations in seawater for more than 17,000 *psbO* reference sequences across all size fractions from *Tara* Oceans, *Tara Oceans* Polar Circle and *Tara Pacific* stations. We have summarized the resultant data as seawater cell concentrations for 15 photosynthetic lineages of eukaryotes and prokaryotes across all size classes sampled during the three expeditions.

### Validation of DNA concentration as a Quantitative Metric

We verified that the sampling process did not introduce random or non-quantitative biases that would prevent us from using DNA concentration estimates as reliable proxies for plankton biomass or abundance. We expect to underestimate total DNA concentrations and taxon-specific cell abundances, as the overall sampling workflow, including water collection, filtration, and DNA extraction and purification procedure, is inevitably less than 100%-efficient in recovering and quantifying DNA. Our goal here is to ensure that variations in DNA concentrations remain representative of actual *in situ* variation, enabling quantitative comparisons between samples rather than only compositional ones.

Chlorophyll-*a* is widely used as a proxy for phytoplankton and total planktonic biomass (*49*). Across the three expeditions, *in-situ* chlorophyll-*a* concentrations correlate significantly with total DNA concentrations (*R_pearson_* = 0.36–0.48, 6.8 × 10^-8^ < *p* < 0.0041) (Fig. 1). This correlation indicates that variations in estimated DNA concentrations summed over all sampled size fractions reflect changes in plankton biomass. In addition, for each size fraction we compared DNA concentration estimates with independent biovolume measurements from imaging, microscopy, and flow cytometry (*50–53*). Except for the 0.2-3 µm prokaryote-enriched fraction, DNA concentrations correlate strongly with organism biovolumes (0.4 < *R_Pearson_* < 0.81, 2.2e-16 < *p_adj_* < 9e-4) (Fig. 1, Fig. S3). For the 0.2–3 µm fraction, we hypothesized that the weak correlations observed between total DNA concentration estimates and organism biovolumes (Fig. S3) may result from the presence of eukaryotic DNA. Eukaryotic picoalgal genomes, expected within this fraction, are several orders of magnitude larger than prokaryotic genomes and could distort the DNA-content-to-cell-volume relationship. This hypothesis will be examined when estimating cell concentrations of photosynthetic lineages across different size fractions.

**Fig. 1.**
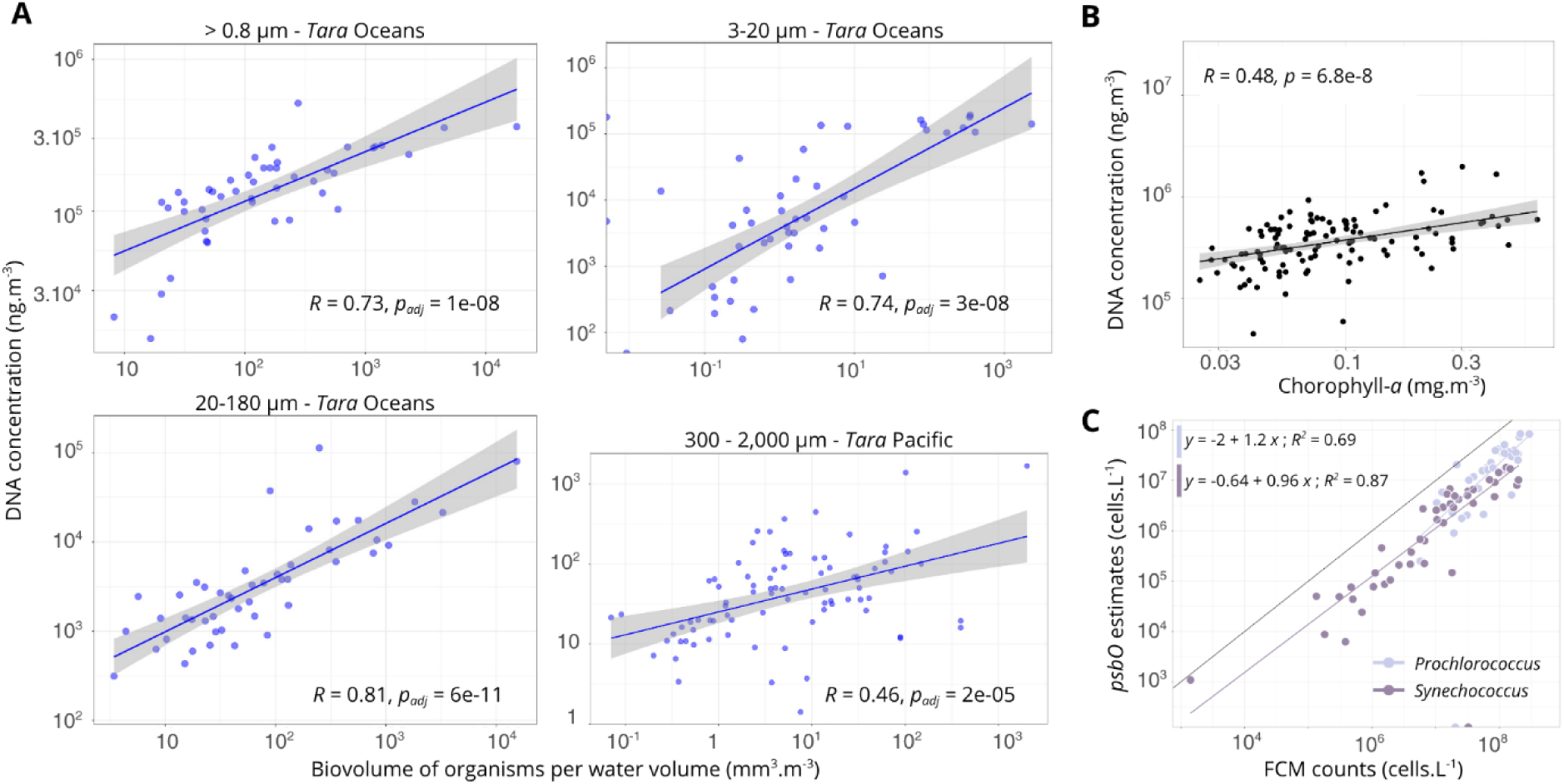
DNA-based concentration estimates correlate with independent quantitative measurements of chlorophyll-a, plankton biovolume and cell abundance. (A) Estimates of DNA concentrations in seawater from Tara Oceans, Tara Oceans Polar Circle and Tara Pacific samples compared to organism biovolumes per water volume, measured for the corresponding size fractions and stations. For Tara Oceans and Tara Oceans Polar Circle, only samples from stations 66 to 210 are shown. **(B)** Estimates of total DNA concentrations in seawater (summed across the 0.2 to 2,000 µm size range) at Tara Pacific stations compared to HPLC-derived chlorophyll-a concentrations. **(A)** and **(B)**: Pearson correlation coefficients are indicated. **(C)** Cell abundances of the cyanobacteria Prochlorococcus and Synechococcus estimated from DNA-based quantification of the psbO marker gene versus flow cytometry cell counts across Tara Oceans and Tara Oceans Polar Circle stations. The y-axis shows cyanobacterial cell concentrations estimated from psbO quantification in metagenomes from the 0.2-3 μm size fraction. The x-axis shows corresponding flow cytometry cell counts. Synechococcus: RPearson = 0.93, p-val < 2.2e-16; Prochlorococcus: RPearson = 0.83, p-val = 9.9e-11.

Specifically, we performed additional controls targeting *Synechococcus* and *Prochlorococcus*, which are considered the most abundant cyanobacteria in this size range. To do so, we compared their estimated cell numbers obtained from flow cytometry data with those derived from our method applied to the *psbO* marker gene (described below). Cell concentration estimates for both cyanobacteria genera in the 0.2-3 µm fraction demonstrate robust log-log linear relationships with flow cytometry cell counts across *Tara Oceans* and *Tara* Oceans *Polar Circle* stations (Fig. 1). Similar analyses conducted on *Tara Pacific* data show strong correlations for *Synechococcus* (*R_Pearson_* = 0.84, *p-val* < 6e-12), while *Prochlorococcus* exhibited a weaker association (*R_Pearson_ =* 0.25*, p-val =* 0.18) (Fig. S4). This observation is consistent with reports that *Prochlorococcus* auto-fluorescence is more difficult to detect by flow cytometry in high-light surface waters compared to *Synechococcus* (*54*). This aligns with *Tara* Pacific samples being collected from the very surface (0-1m), whereas *Tara* Oceans samples were obtained from around 5m depth.

For both *Synechococcus* and *Prochlorococcus*, the slopes of the linear regression lines were not significantly different from 1 (Fig. 1) (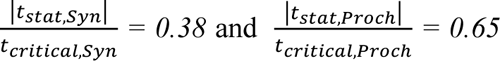 for a significance level of 0.05), indicating that the relative error of DNA-based estimates remains approximately constant across different orders of magnitude of cell concentration values. However, this relative error appears to introduce a systematic underestimation of actual concentrations, by a factor of five for *Prochlorococcus* and by one order of magnitude for *Synechococcus*. The underlying causes, consequences and limitations of this bias are further examined in the Discussion section. Although DNA and cell concentrations presented in this study are likely underestimated, the consistent relative error validates the use of our DNA-based estimates as reliable quantitative proxies for taxon-specific abundance. Also, the much stronger correlations obtained when targeting well-characterized taxa rather than the whole community were expected, likely reflecting a greater number of uncontrolled factors in the latter case. Nevertheless, this confirms that the approach performs well when focusing on known taxa, without excluding its potential use for studying abundance variations in total communities.

### Converting Relative to Absolute Genome Abundance Using DNA Concentration

Based on the mapping of *Tara* Oceans and *Tara* Oceans Polar Circle metagenomes to two genome collections (Methods), we estimated DNA concentrations in seawater for 713 eukaryotic metagenome-assembled genomes (MAGs) (*38*) and 8,196 prokaryotic MAGs and reference genomes (*26*, *37*) across sampled size fractions.

Echoing previous studies on quantitative metagenomic profiling of microbial communities (*27*, *47*), we carried out comparative analysis between relative and quantitative metagenomic approaches to highlight how quantifying absolute abundances of genomes or lineages affects the representation of plankton community structure (Fig. 2).

**Fig. 2.**
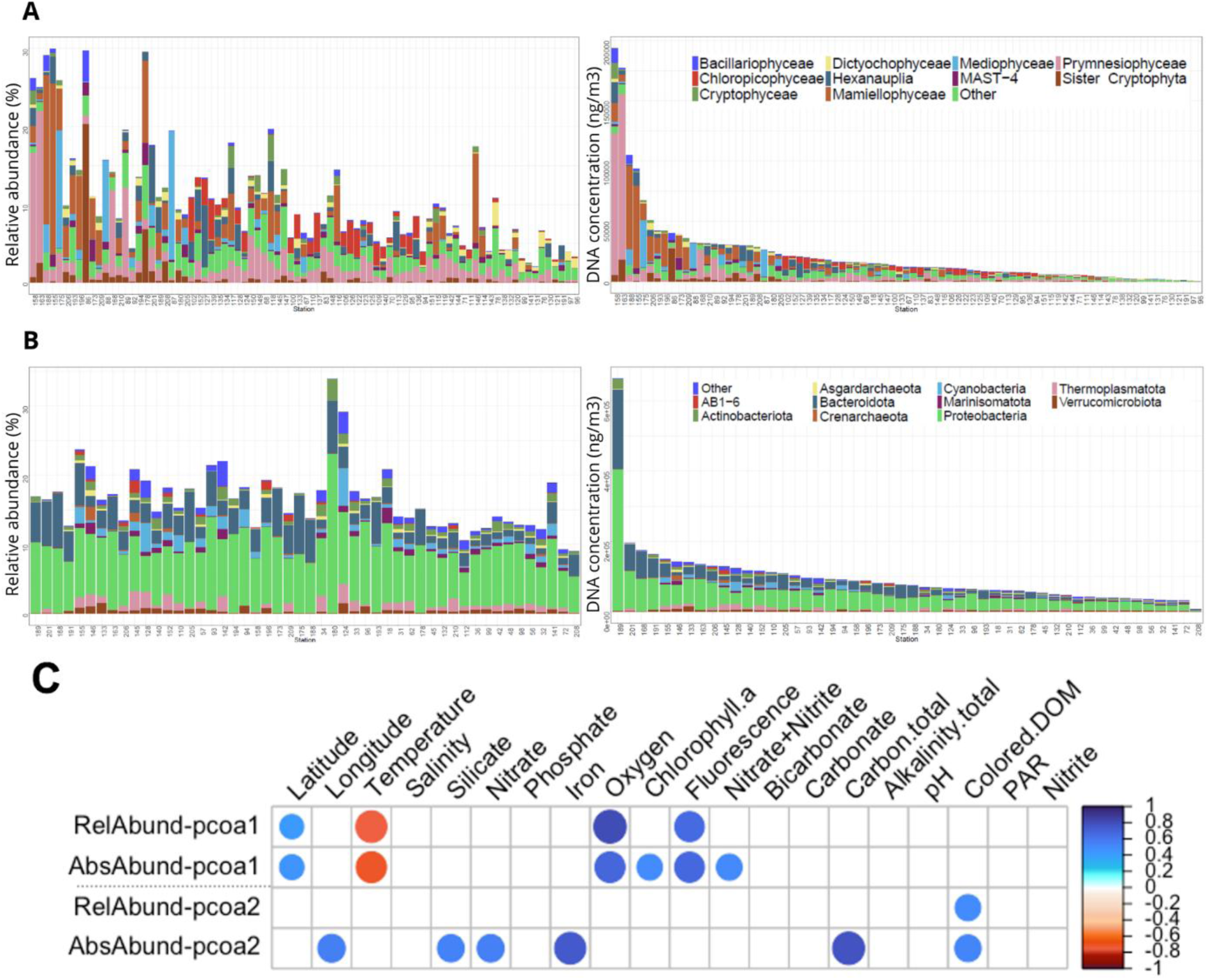
Quantitative metagenomic profiles differ from relative abundances and reveal additional environmental drivers of community structure. **(A-B)** Relative and quantitative metagenomic profiles of samples from the 0.8-2,000 µm **(A)** and 0.2-3 µm **(B)** size fractions across the Tara Oceans and Tara Oceans Polar Circle stations. (Left) Relative metagenomic abundances of the 10 most abundant eukaryotic MAG classes **(A)** and prokaryotic MAG phyla **(B)** across all metagenomes in the respective size fractions. Remaining MAGs are grouped under the ‘Other’ category. (Right) DNA concentrations (ng.m-3) in seawater for the same 10 classes **(A)** and phyla **(B)** associated with samples from the 0.8-2,000 µm **(A)** and 0.2-3 µm **(B)** size fractions. Stations are indicated on the vertical axis, in the same orders for both ‘relative’ and ‘quantitative’ histograms (sorted by decreasing total DNA concentration). Relative abundances do not sum to 100% because MAGs account for only a fraction of the total microbial diversity captured by metagenomic sequencing. **(C)** Correlation of Principal Coordinate Analysis (PCoA, Fig. S5) axes with various environmental parameters. Comparisons of correlations were computed between non-polar community ordination plots generated based on relative and absolute abundance matrices. White boxes indicate non-significant correlations (p > 0.01). See Fig. S5 for correlation plot of the polar community.

As expected, large variations in total DNA concentration across stations (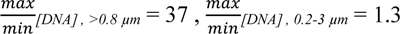) changes the ranking of stations based on absolute concentrations of a given genome or lineage. For example, this effect is evident in the distributions of relative and absolute abundances of the green picoalgae genera *Bathycoccus* and *Micromonas* (Fig. 2), as well as the bacterial phylum *Proteobacteria* (Fig. 2), for which stations exhibiting the highest relative abundances (e.g., stations 180 and 146) are in fact among those with the lowest DNA concentrations for these taxa.

The highest DNA concentrations for both prokaryotes and eukaryotes, across both size fractions, were observed at high-latitude stations (Fig. 2). At this stage, we did not compare total DNA or cell concentrations between lineages or stations, as the two collections of MAGs incompletely represent plankton communities, evidenced by low percentages of mapped metagenomic reads (*26*, *37*, *38*). Prokaryotic MAGs accounted for at most 34% of the total reads sequenced in the 0.2-3 µm fraction (minimum 7.5%; mean 17%; Cyanobacteria 5%). Eukaryotic MAGs recruited only up to 40% of the total number of reads in the eukaryote-enriched samples (minimum 0; mean 11%). Additionally, the panel and level of diversity represented in each collection vary across taxa, potentially biasing representation of lineages.

We then investigated how absolute genome quantification could influence the evaluation of their associations with abiotic factors. Spearman correlation between key environmental parameters and the first two Principal Coordinate Analysis (PCoA) axes derived from polar and non-polar relative and absolute abundance dissimilarity matrices (Fig. 2, Fig. S5) showed largely coherent correlation patterns between absolute and relative community profiles and environmental parameters, yet with a few important exceptions. Within the non-polar community, five environmental parameters (latitude, temperature, oxygen, fluorescence, and dissolved organic matter) correlated significantly with both relative and quantitative datasets (*55–57*). In contrast, seven parameters (longitude, chlorophyll-*a*, NO₂+NO₃, silicate, nitrate, iron, and total carbon) showed significant correlations only when using quantitative abundance estimates. These environmental parameters reflect nutrient concentrations, light availability and organic matter quality, all key drivers of microbial community structures. Notably, quantitative estimates revealed the expected relationship between total carbon and microbial community biomass, consistent with the coupling between carbon availability and total community mass, which becomes evident here only when using absolute rather than relative data. Such patterns suggest that absolute abundance data, by incorporating a quantitative dimension of community organization, may better capture ecologically relevant environmental parameters and provide a more comprehensive understanding of community–environment interactions, even when communities are only partially resolved, as with MAGs.

We examined how MAG co-occurrence networks, used as proxies for ecological interactions, differ when reconstructed from relative versus absolute abundance matrices. Using the same set of MAGs in both cases (after removing low abundance MAGs to limit spurious correlations), the absolute abundance network contained substantially more edges (616,780 versus 438,967 in the relative abundance network), with a vast majority being positive (99.9% *vs* 64.9% for absolute and relative abundance networks, respectively; Fig. S6). Average node connectivity (mean degree) was also higher in the absolute abundance network (4.32-7.13 compared to 2.59-4.43 in relative abundance network; Table S1). Similar trends have also been consistently observed in previous studies of various ecosystems comparing co-occurrence networks derived from quantitative and relative data (*27*, *58*, *59*), highlighting the potential for reducing negative correlation biases and spurious associations (*15*).

We further analyzed the topology of interactions within the 616,210 and 284,704 positive edges from the absolute and relative abundance networks (Fig. 3). The ratio of inter-kingdom associations remains fairly similar in both networks. However, the absolute abundance network detected a slightly higher percentage of Bacteria-Eukaryote associations, even though it had a lower percentage of inter-kingdom associations. As a result, many key predicted interactions (e.g., Rhodobacteraceae↔*Bathycoccus*; Rhodobacteraceae↔*Phaeocystis*; *Prochlorococcus*↔*Phaeocystis*; Rhodobacteraceae↔Marine Hexanauplia A; Flavobacteriaceae↔ Marine Hexanauplia A) are much more represented in the absolute abundance network, with many of the predicted Bacteria-Eukaryote interactions detected only in this network.

**Fig. 3.**
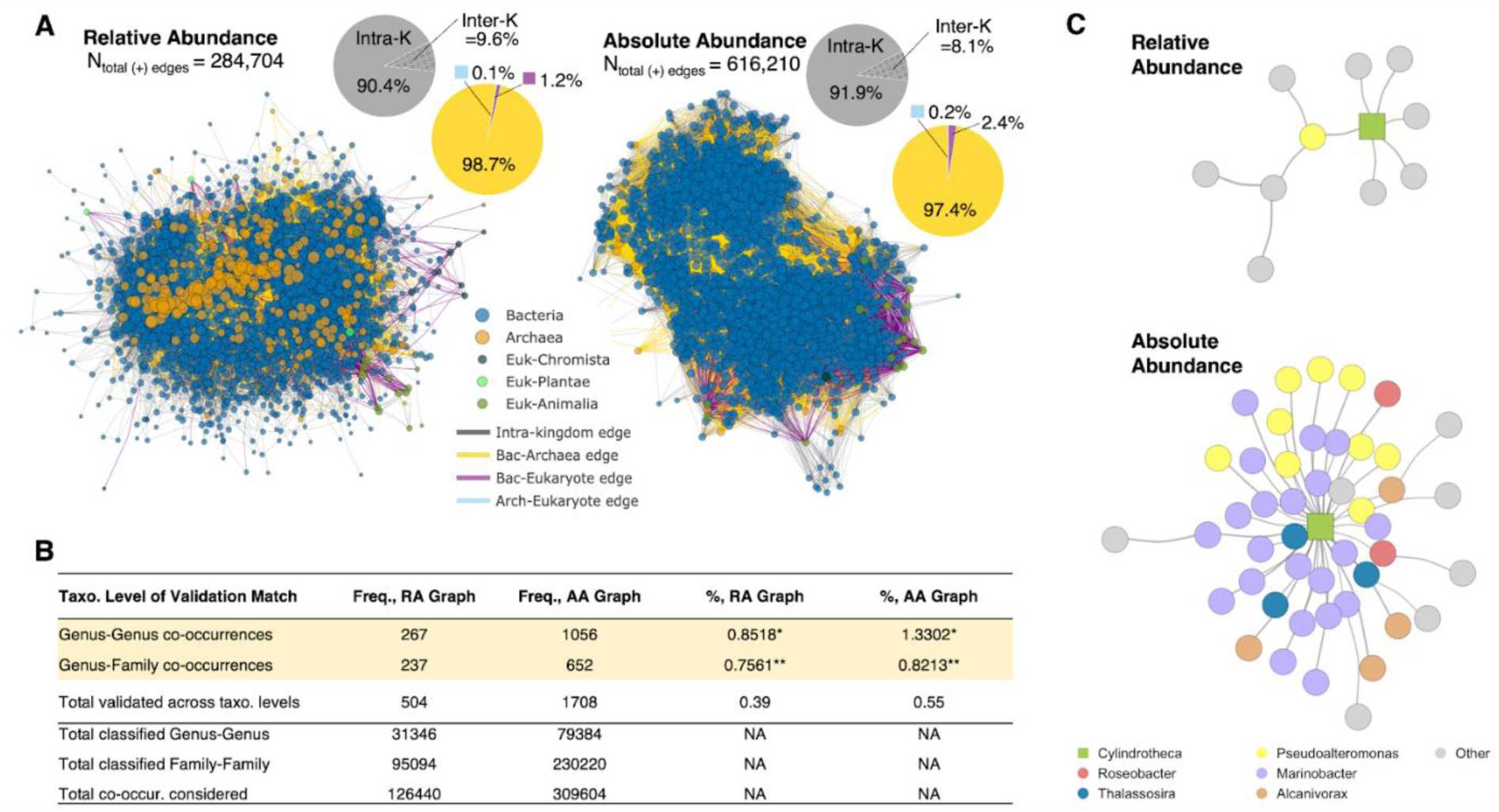
Quantitative genome abundances improve ecological interaction networks inference. **(A)** Co-occurrence networks reconstructed using bacterial, archaeal and eukaryote MAG relative (left) and absolute (right) abundance matrices. Pie chart colors indicate inter-kingdom (Inter-K) edge categories **(B)** Summary of frequency and percentages of recovered associations that are validated against known biotic interactions from the GLOBI database, up to family level. **(C)** Subnetworks generated from relative (top) and absolute (bottom) abundance matrices, highlighting inferred eukaryote-prokaryote co-occurrences. “Other” refers to additional nodes connected to the eukaryote-prokaryote co-occurrence subset. * - fraction of total co-occurrences where both MAGs were classifiable at the genus level; ** - fraction of total co-occurrences where both MAGs were classifiable at the family level.

We aimed to confirm these results with curated interaction data from the Global Biotic Interactions (GLOBI) database (*60*). Considering only co-occurrences between MAGs classifiable to genus level, the absolute abundance network recovered a moderately higher proportion of known interactions from literature (Fig. 3). But more importantly, the absolute abundance network revealed higher percentages (2.3% versus 0.19% in the relative abundance network) of validated Bacteria-Eukaryote interactions with a higher diversity of taxa. For example, the relative abundance network recovers only one validated interaction (between the diatom *Cylindrotheca* with bacteria *Pseudoalteromonas*), whereas the absolute abundance network recover 39 validated interactions between *Cylindrotheca* with 5 different bacterial genera (*Pseudoalteromonas*, *Roseobacter*, *Marinobacter, Thalassosira* and *Alcanivorax*) (Fig. 3). Taken together, these results suggest that the absolute abundance network provides a much broader coverage of validated interactions and is the more robust framework for predicting novel biotic interactions with higher confidence.

### Quantitative Estimates of photosynthetic lineages using the *psbO* marker gene

Using the *psbO* reference database compiled by Pierella Karlusich et al. (*9*), we estimated absolute cell concentrations of over 10,000 eukaryotic and prokaryotic *psbO* sequences detected across more than 1,000 surface samples from *Tara* Oceans, *Tara* Oceans Polar Circle and *Tara* Pacific. Based on prior taxonomic annotations, these sequences are classified into 15 photosynthetic lineages.

Eukaryotic lineages were predominantly found in small protist-enriched size fractions (0.8-5 µm, 0.8-2,000 µm and 3-20 µm), and most appeared in roughly similar abundances in the bacteria-enriched fraction (0.2-3 µm), supporting our previous hypothesis that a high proportion of eukaryotic DNA in this size-fraction affected validation based on total DNA mass (Fig. S3). Median concentration within these size classes were generally about one order of magnitude higher than those observed in the >20 µm fractions (Fig. S7). In contrast, *Prochlorococcus*, *Synechococcus* and the group of other cyanobacteria were found to be significantly more abundant in the 0.2-3 µm size fraction, with median concentrations differing by at least one order of magnitude from the other small size fractions (Fig. S7). Consistent cell concentrations of the cyanobacterial genus *Trichodesmium* were estimated across multiple size fractions from 0.2 to >20 µm, likely reflecting the presence of their different biological forms, including free-living cells, filaments, and colonies of varying sizes (*36*).

We found that *Prochlorococcus*, *Synechococcus* and *Chlorophyta,* while being the most abundant groups in the smallest size fraction (0.2-3 µm), also exhibited significant cell concentrations in the largest size classes (Fig. S7). Specifically, *Prochlorococcus* and *Synechococcus* reached concentrations above 1 × 10^2^ cells.L^-1^ in the >180 µm and >300 µm size fractions, and exceeded 1 × 10^3^ cells.L^-1^ and 1 × 10^4^ cells.L^-1^, respectively, in the other >20 µm fractions (Fig. S7). These observations are congruent with previous confocal microscopy and metagenomics studies conducted on the same sampled water masses, which reported *Synechococcus* and *Prochlorococcus* cells in large-size-fraction samples, revealing symbiotic or aggregate-forming cyanobacterial taxa (*9*, *61*). Here, the actual quantification of *psbO* gene sequences confirms that the relatively high proportions of cyanobacteria observed in large size-fraction samples are not compositional artifacts arising from the collapse in abundance or biomass of other organisms normally dominant in these size fractions.

One of the objectives of our method is to enable the summation of abundances across plankton size fractions for a given sampling site. To estimate cell concentrations for comprehensive phytoplankton communities spanning the 0.2-2,000 µm size range, we applied summation criteria to ensure consistency across stations and lineages (see Materials and Methods). After summation across size fractions, the cyanobacterial genera *Prochlorococcus* and *Synechococcus* are the most abundant cells in phytoplankton communities from 0.2 to 2,000 µm (Fig. 4), as expected. When present, they were observed at median abundances of 2 × 10^7^ cells.L^-1^ and 1 × 10^6^ cells.L^-1^, respectively, and with maximum concentrations above 1 × 10^8^ cells.L^-1^ and 3 × 10^7^ cells.L^-1^, respectively. These results likely carry the same underestimation biases as those seen with flow cytometry controls. Still, the estimated cell concentrations are within the reported ranges for both cyanobacterial genera in pelagic ecosystems (*62*, *63*).

**Fig. 4.**
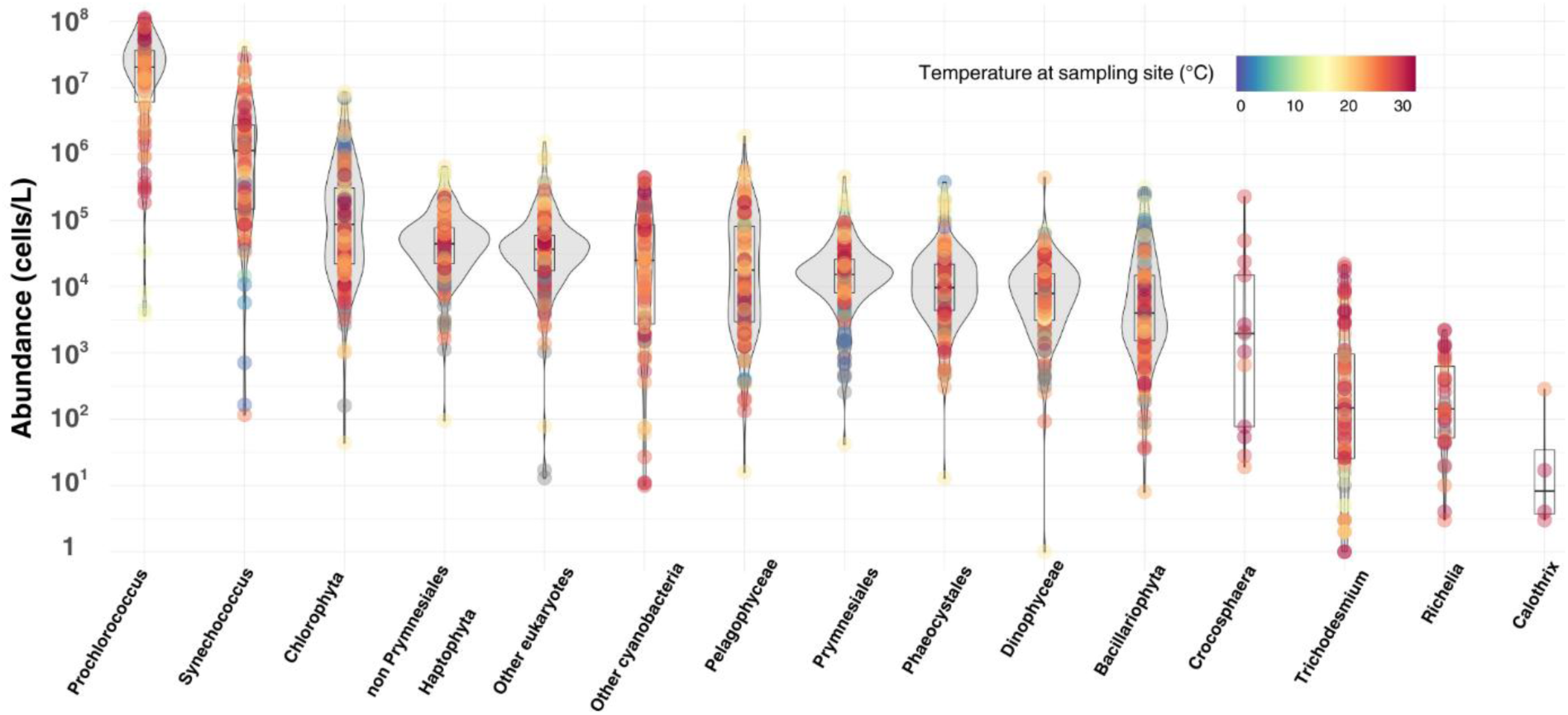
Estimates of total cell abundance comparing eukaryotic and prokaryotic phytoplankton lineages ranging in size from 0.2 to 2,000 μm across polar, temperate, and tropical sampling sites. Each colored dot represents a sampling station, with color indicating sea surface temperature at the time of sampling. Boxplots show the median and interquartile range of cell abundance values per lineage (excluding zero values). Lineages are defined by the psbO marker gene and are ordered by decreasing median abundance.

The phylum *Chlorophyta* exhibited the highest cell concentrations among eukaryotic lineages with values of detected abundances ranging from 30 cells.L^-1^ to above 1 × 10^7^ cells.L^-1^, with a median abundance of 9 × 10^4^ cells.L^-1^. Diatoms had the lowest average and median cell concentrations among eukaryotes, with a median cell concentration of 4 × 10^3^ cells.L^-1^ and a maximum abundance of 3 × 10^5^ cells.L^-1^. For diatoms, these estimates are lower than the median value reported in the MAREDAT global survey and the maximum abundance estimated by the quantitative metagenomic assessment of the *psbO* gene by Bei et al. (*34*, *64*), approximately 10⁵ cells.L⁻¹ and > 10^6^ cells.L⁻¹ at the surface, respectively. Our estimates are still consistent with other observations of diatom abundance in pelagic environments (*64*, *65*). For diatoms, assessing the concordance between our estimates and published cell abundances is challenging, as reported concentration ranges are often broad, with most studies encompassing pelagic and coastal waters, as well as blooming areas (*64*).

All the other eukaryotic lineages displayed median cell concentrations within less than one order of magnitude, ranging from approximately 8 × 10^3^ cells.L^-1^ for *Dinophyceae* to 4 × 10^4^ cells.L^-1^ for non-*Prymnesiales* haptophytes. Notably, the “other eukaryotes” group, mainly composed of Ochrophytes *psbO* sequences that could not be further taxonomically classified, constituted a large part of the phytoplankton community, representing the third most abundant group of photosynthetic eukaryotes. These estimates of eukaryotic phytoplankton cell concentrations generally fall within the broad range of cell abundance values reported in the literature for open-ocean environments, but they are consistently lower than average cell abundances typically measured using imaging techniques, microscopy, or flow cytometry (*34*, *66*, *67*).

Size-fractionated sampling is a methodological approach designed to address the decrease in plankton organism concentration with increasing body size (*40*). The volume of seawater that must be filtered to collect sufficient numbers of organisms larger than 20 µm exceeds, by several orders of magnitude, that required for bacteria. Consequently, unfiltered seawater samples are inherently biased toward the smallest components of the community. On the other hand, compositional data derived from traditional size-fractionated filtrations prevent integrating the abundance of a given taxon across multiple size fractions. Here, our approach allows metagenomic-derived cell concentrations to be summed across size fractions by accounting for differences in filtered seawater volume and other dilution factors. In this specific analysis, some samples from large size fractions may be missing for a given *psbO* lineage, causing a slight heterogeneity in cumulative cell concentration estimates. However, we verified that missing larger size fractions did not significantly affect total cell concentrations from 0.2 to 2,000 µm. Median, and average cell abundances remained consistent with cell concentration estimates from 0.2 to 20 µm (Fig. S8).

### Quantified Biogeography of Phytoplankton

To present a synoptic view of phytoplankton absolute abundances across major taxonomic groups, we developed a global ensemble modeling framework allying the derived cell abundance dataset with global environmental variables (Fig. S1). This integration of empirical quantification with predictive modeling offers a unified, quantitative perspective on phytoplankton biogeography, with a potential to support their incorporation into biogeochemical and carbon flux frameworks (*68–70*). To this end, we trained regression models for each group on log-transformed cell counts based on *psbO*-based estimates of total cell concentrations pooled across the size ranges from 0.2 to 2,000 µm (Fig. 5, Fig. S10). The ensemble models relied on a suite of key environmental predictors, sea surface temperature (SST), sea surface salinity (SSS), chlorophyll-*a*, iron, nitrate, phosphate, and silicate, chosen for their established influence on phytoplankton ecology and biogeography. Model predictions were made over monthly global environmental grids, with phytoplankton maps generated at 1° × 1° resolution. To quantify prediction confidence and extrapolation risk, we computed Mahalanobis distances between environmental conditions at each grid point and the training dataset, identifying regions outside the empirical environmental envelope. In parallel, we conducted sensitivity analyses by varying subsets of the data, quantifying how training data composition influences global predictions. These diagnostics allowed us to distinguish robust patterns from regions of higher uncertainty and guide the ecological interpretation of predicted biogeographies (see Discussion section).

**Fig. 5.**
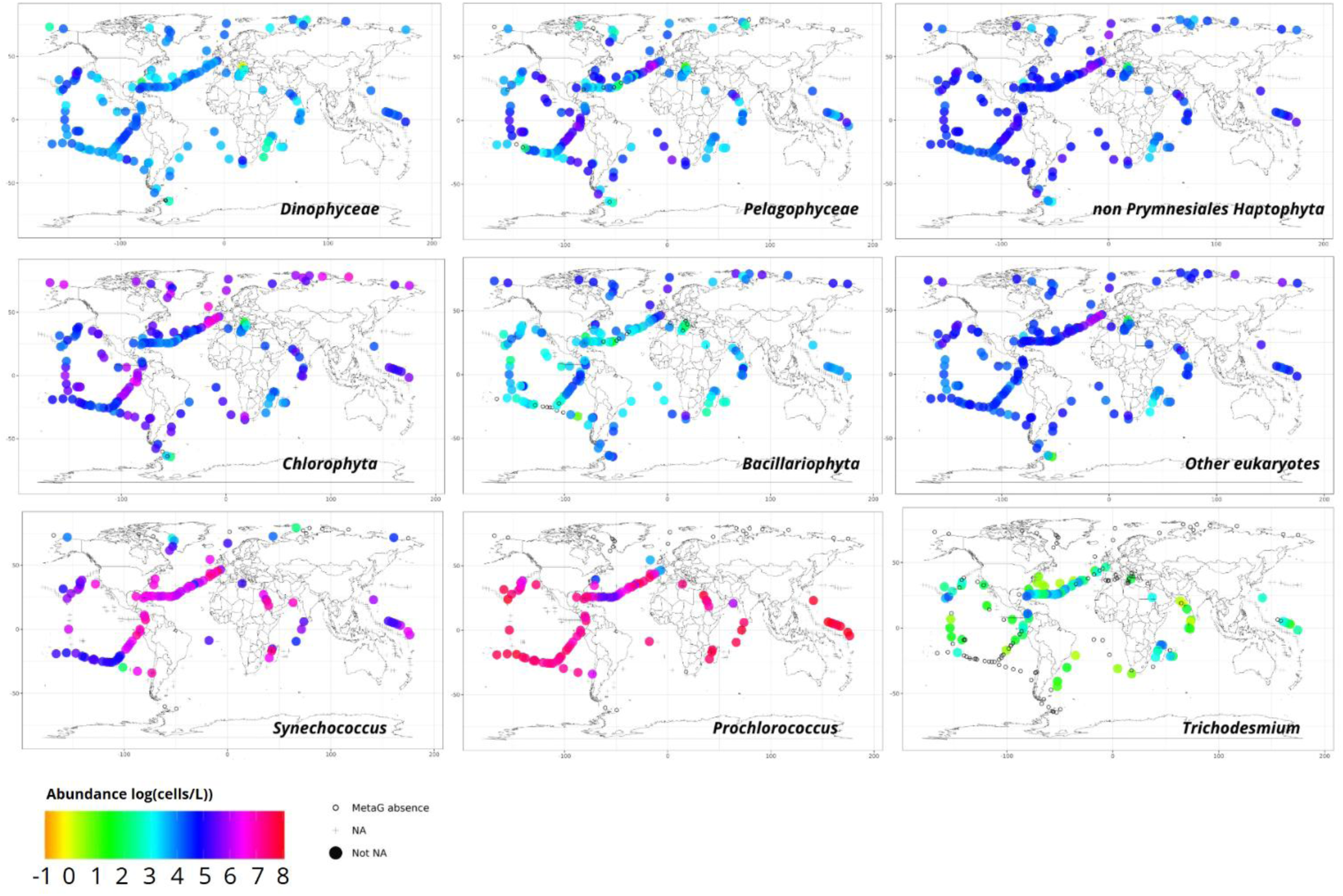
Global distribution of estimated cell concentrations across major phytoplankton lineages. Cell abundance values (log10(cells.L-1)) represent cumulative concentrations from 0.2 to 2,000 µm across surface stations of the Tara Oceans, Tara Oceans Polar Circle and Tara Pacific expeditions. Phytoplankton lineages are defined by the psbO marker gene. Crosses (+) indicate stations with missing data for the corresponding lineage. Black circles (○) indicate stations where the lineage was not detected in the available metagenomes.

The ensemble model reproduces a pronounced latitudinal increase in total phytoplankton abundances toward the equator, with maxima in equatorial and subtropical gyres and lower abundances toward the poles. Predicted totals span over two orders of magnitude (10⁶-10⁸ cells.L^-1^) (Fig. 6). These large-scale gradients are consistent with the driver analysis from the ensemble model (Fig. S12), which identifies SST as the dominant predictor structuring phytoplankton communities (*71*, *72*). This mirrors results from metagenomic-based community analyses, where absolute quantification revealed stronger and more coherent correlations with environmental gradients such as temperature (Fig. 2), highlighting its central role in microbial distributions.

**Fig. 6.**
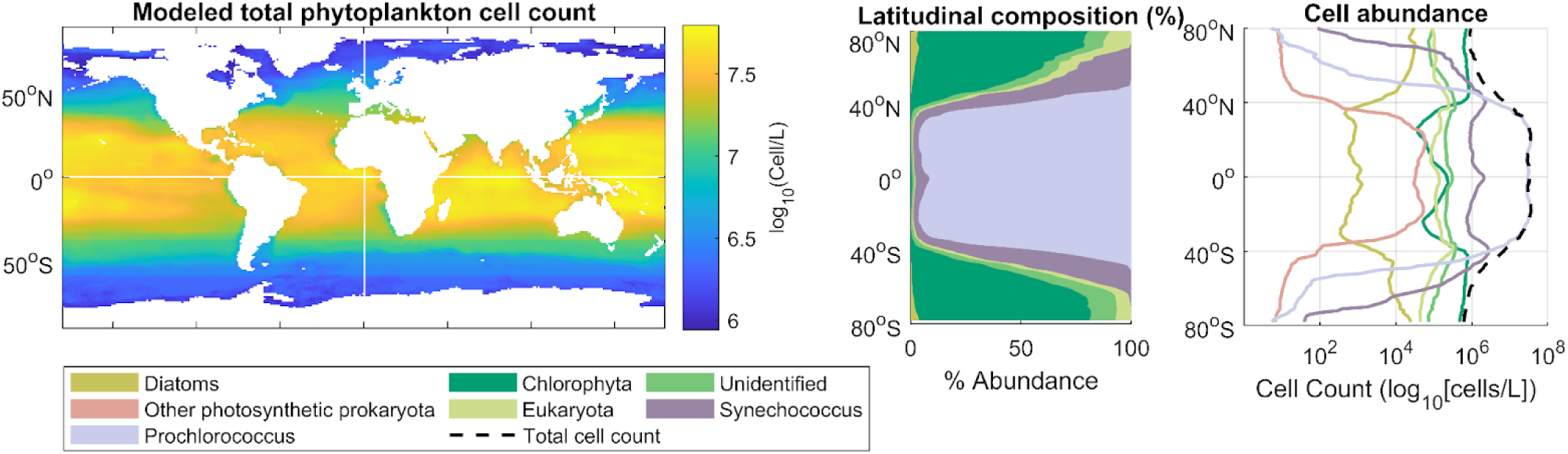
Modeled distribution and latitudinal variation of phytoplankton cell abundance and composition. (Left) Global map of total phytoplankton cell concentration (log₁₀ cells L⁻¹). (Center) Latitudinal composition (% abundance) of major phytoplankton groups, showing relative dominance across latitude bands. (Right) Latitudinal variation in absolute cell concentration (log₁₀ cells L⁻¹) for each group, with the dashed line indicating total modeled cell concentration. Colors represent functional groups: Diatoms, Other photosynthetic prokaryotes, Prochlorococcus, Chlorophyta, Other Eukaryota, Synechococcus, and Unidentified.

Cyanobacteria appeared predominantly abundant in temperate and warm-water stations (Fig. 5): *Synechococcus* declined sharply above 50° N, while *Prochlorococcus* and *Trichodesmium*, along with almost all other cyanobacterial genera, were absent from stations above 55° N and from cold-water stations in the Southern Ocean and South Atlantic. This latitudinal structuring of cyanobacterial abundances, dominated by *Prochlorococcus* and *Synechococcus*, is consistent with well-established patterns from flow cytometry and quantitative molecular surveys (*62*, *73*). Both datasets agree on the ecological transition from cyanobacterial dominance in low latitude oligotrophic waters to more diverse, eukaryote-rich assemblages at higher latitudes. *Tara* observations, similarly to other large quantitative surveys (via cytometry data compilation (*63*)), show *Prochlorococcus* to be uniform in abundance across low-latitude stations, while *Synechococcus* and *Trichodesmium* display more heterogeneous distributions, with *Synechococcus* declining near hydrographic boundaries such as the Humboldt Current and *Trichodesmium* showing patchy hotspots in the North Pacific, Caribbean, and Mozambique Channel, interspersed with stations below the detection threshold (1 cell.L⁻¹; e.g., Cape Agulhas, Marquesas Islands).

The model captures the broad-scale uniformity of *Prochlorococcus*, the decline of *Synechococcus* above 55°, and the emergence of chlorophytes in temperate and polar waters (*63*, *74*, *75*). Because *Trichodesmium* and *Richelia* blooms with symbiotic diazotrophs are sparse and transient, the limited and uneven training data lead to high predictive uncertainty. While *Tara* observations demonstrate sporadic patterns for these diazotrophs, related to local bloom dynamics, the model identifies global patterns consistently more abundant in tropical oligotrophic regions, as observed in prior large-scale surveys (*76*, *77*). For instance, looking at the ensemble model predictions and drivers importance (Fig. S12), diazotrophs display elevated sensitivity to phosphate and, to a lesser extent, iron, both essential for nitrogen fixation and frequently limiting in subtropical and oligotrophic systems (*78–80*). Nitrate, by contrast, appears less important, consistent with the ecological niche of diazotrophs in nitrate-depleted environments where they avoid competition with nitrate-assimilating phytoplankton.

Eukaryotic groups show equally strong concordance between observations and predictions. Data reveal that haptophyte lineages (excluding Phaeocystales), dinoflagellates, and other eukaryotes display relatively homogeneous abundances across temperate and tropical basins, with declines near the Agulhas Current and in the Southern Ocean. In contrast, pelagophytes, diatoms, and green algae exhibit more heterogeneous cell distributions within each ocean basin. Diatoms peak in high-latitude, nutrient-rich regions; pelagophytes are more abundant in tropical and subtropical waters; and chlorophytes are consistently the most abundant photosynthetic eukaryotes across all basins, including the polar biome. The model reproduces these patterns: Phaeocystales are concentrated in high-latitude and upwelling regions; diatoms and chlorophytes dominate nutrient-rich temperate and polar waters; and haptophytocalcifiers and pelagophytes occur broadly in temperate bands (*81*, *82*). Observations also identify localized declines in pelagophyte and diatom abundances around 100° W in the eastern tropical Pacific, aligned with the transition from the Humboldt Current to the oligotrophic South Pacific gyre.

At the community level, the ensemble model demonstrates additive coherence: summing group-level abundances closely matches total phytoplankton predictions (Fig. S11, Fig. S12). Dominant groups (*Prochlorococcus*, *Synechococcus*, chlorophytes) drive both the strongest observational signals and the highest model skill, while bloom-forming or patchy taxa contribute less to global totals and therefore introduce limited instability in reconstructions. This stabilizing effect explains why, despite variability in taxon-specific accuracy, the ensemble model captures global total abundances with high fidelity.

Model-predicted cell abundances display an inverse gradient with global patterns of chlorophyll-*a*, reflecting the contrasting dominance of cyanobacteria in oligotrophic gyres (high abundance, low Chl-*a*) versus large eukaryotes in high-latitude, nutrient-rich regions (lower abundance, high Chl-*a*). Second in the importance rank (Fig. S13), chlorophyll-*a* likely acts as an integrative proxy for ecosystem productivity and trophic status. However, it is important to interpret the role of chlorophyll-*a* in our model carefully: as a biologically derived product, it can be both a predictor and a response variable, and its statistical importance here likely reflects shared covariance with phytoplankton cell density rather than a direct mechanistic influence. The scarcity and patchiness of observations for several groups, especially diazotrophs, limit the ability to disentangle true environmental drivers from covariate artifacts. Nonetheless, the congruence between the learned dependencies and biological expectations lends credibility to the model’s capacity to extract ecologically meaningful structure, even in data-limited regimes.

Model uncertainty was closely linked to the geography of the training data rather than to algorithmic instability. Under-sampled environments such as the Southern Ocean, upwelling systems, and some marginal seas exhibited higher extrapolation risk, confirmed by Mahalanobis distance analyses (Fig. S10, Fig. S11). In contrast, tropical and temperate oceans, well covered by training data, showed high predictive confidence.

To complement abundance-based estimates, we converted phytoplankton cell concentrations estimated at each station into carbon biomass. This step is essential because carbon biomass represents the currency of marine biogeochemistry (*1*): it links community structure to primary production, trophic transfer, and global carbon cycling. Abundance alone does not capture differences in cell size or carbon content between taxa, whereas biomass provides a more ecologically meaningful metric for comparing contributions across groups. Carbon mass is also a commonly used unit in biogeochemical models that represent multiple phytoplankton groups such as DARWIN (*83*), PISCES (*24*), and PlankTOM (*84*). Therefore, expressing genomic data in units of carbon mass represents a step toward integrating genomic resolution into the next generation of biogeochemical models. For each taxonomic group, carbon biomass was estimated from cell abundances using established group-specific conversion factors (Table 2), applied consistently across size classes.

**Table 1.** Methods for calculations of effective filtered volumes depending on seawater collection and filtration strategies for *Tara* Oceans, Polar Circle and *Tara* Pacific expeditions.

| Size fraction (µm) | Event device | Volume of water collected | Resuspension | Final filtration on filter sample | “Effective” volume of seawater sample |
| --- | --- | --- | --- | --- | --- |
| 0.2-3 | HVP - PUMP | Vc = 100 L (carboys) | / | Vf | V = Vf |
| 3-20 | HVP - PUMP | Vc = 100 L (carboys) | / | Vf | V = Vf |
| 0.8-5 | HVP - PUMP + GPSS | Vc = 100 L (carboys) | / | Vf | V = Vf |
| 0.8-2,000 | HVP - PUMP | Vc = 100 L (carboys) | / | Vf | V = Vf |
| 5-20 | [NET-SINGLE-5]<br>Single net<br>Opening Area = 0.196 m<sup>2</sup> | Vc = (flowmeter end - flowmeter start)*0.3*0.196 | Vr | Vf | $V = (V_f * V_c) / V_r$ |
| 5-20 | HVP - PUMP + GPSS | Vc = pump end - pump start | Vr | Vf | $V = (V_f * V_c) / V_r$ |
| 20-180 | [NET-DOUBLE-20]<br>2-net instrument<br>Opening Area = 0.192 m<sup>2</sup><br>N = nb of cod end used for resuspension | Vc = (flowmeter end - flowmeter start)*0.3*0.192*N | Vr | Vf | $V = (V_f * V_c) / V_r$ |
| 180-2,000 | [NET-BONGO-180]<br>2-net instrument<br>Opening Area = 0.258 m<sup>2</sup><br>N = nb of cod end used for resuspension | Vc = (flowmeter end - flowmeter start)*0.3*0.192*N | Vr | Vf | $V = (V_f * V_c) / V_r$ |
| 20 - 2,000 | INLINE PUMP + DECKNET-20 | Vc = pump end - pump start | Vr | Vf | $V = (V_f * V_c) / V_r$ |
| 300-2,000 | [HSN-NET-300]<br>Single net<br>Opening Area = 0.116 m<sup>2</sup> | Vc = (flowmeter end - flowmeter start)*0.3*(0.34*0.34) | Vr | Vf | $V = (V_f * V_c) / V_r$ |

**Table 2.** Methodology for conversion of phytoplankton lineages cell abundances into carbon biomass.

| Lineage | Average Carbon content (pgC.cell<sup>-1</sup>) | Average cell size (µm) | Average cell volume (µm<sup>3</sup>) | Conversion equation |
| --- | --- | --- | --- | --- |
| *Trichodesmium* | 300 (76, 120) |  |  |  |
| *Richelia* | 10 (76) |  |  |  |
| *Calothrix* | 10 (76) |  |  |  |
| *Crocospaera* | | | 7 (120) | $C_{pgC.cell-1} = 0.216 * Vol_{\mu m^3}^{0.939}$<br>(110) |
| *Prochlorococcus* | 0.06 (4) |  |  |  |
| *Synechococcus* | 0.15 (4) |  |  |  |
| Other cyanobacteria | | | 1.5 (120–123) | $C_{pgC.cell-1} = 0.216 * Vol_{\mu m^3}^{0.939}$<br>(110) |
| Chlorophytes | 1.3 (4) |  |  |  |
| Pelagophytes | 1.3 (4) |  |  |  |
| Other eukaryotes | 1.3 (4) |  |  |  |
| Phaeocystis | 16 (124) |  |  |  |
| Prymnesiales | | 4 | 30 (82, 125) | $C_{pgC.cell-1} = 0.228 * Vol_{\mu m^3}^{0.899}$<br>(110) |
| Haptophytes non-Prymnesiales | | 4 | 30 (82, 125) | $C_{pgC.cell-1} = 0.228 * Vol_{\mu m^3}^{0.899}$<br>(110) |
| Bacillariophyta | | size fraction lower bound | | $C_{pgC.cell-1} = 0.288 * Vol_{\mu m^3}^{0.811}$<br>(110) |
| Dinophyceae | | size fraction lower bound | | $C_{pgC.cell-1} = 0.216 * Vol_{\mu m^3}^{0.939}$<br>(110) |

As expected, accounting for phytoplankton carbon biomass substantially alters both latitudinal and compositional profiles of the total phytoplankton community. Phytoplankton carbon biomass peaks at high latitudes, particularly at polar stations in the Arctic Ocean and subpolar stations in the North Atlantic Ocean (Fig. 7), where the mean biomass is about five times the average concentration estimated in tropical and subtropical regions. A slight increase in carbon concentrations is observed in the equatorial band between -15° S and 15° N, particularly for Chlorophytes, Pelagophytes, and Haptophytes. Overall, the estimates of total phytoplankton carbon biomass and their latitudinal distribution are consistent with both observed (*4*) and modeled (*85*) values for the global ocean, which typically span up to approximately three orders of magnitude.

**Fig. 7.**
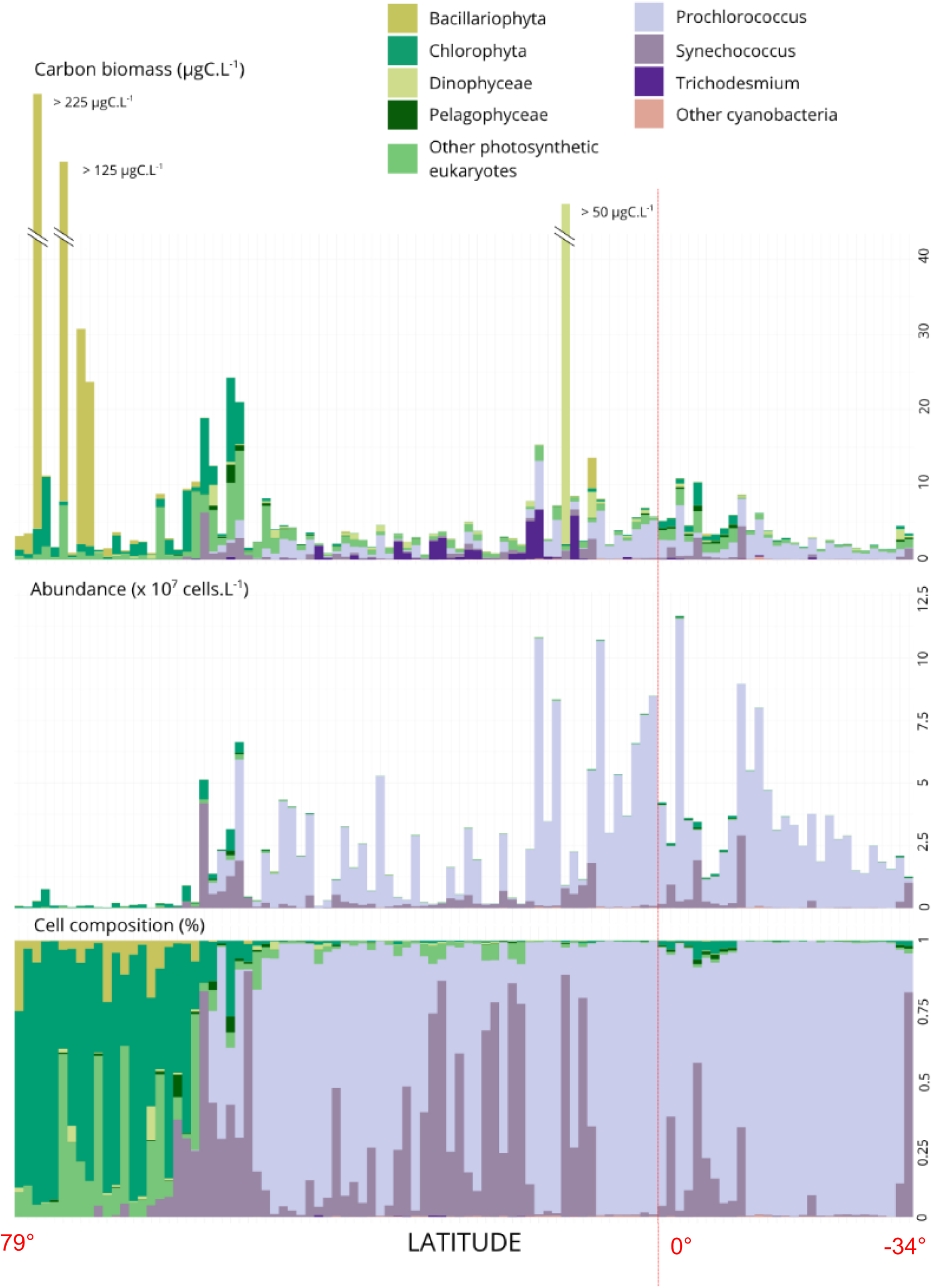
Latitudinal variation in phytoplankton community composition derived from metagenomic-based quantification of *psbO*-lineages, estimated cell abundance, and carbon biomass across global ocean regions, revealing differing patterns among the three metagenomic-based estimates. For each station, community composition is indicated in relative cell composition (bottom), cell abundance (cells.L^-1^) (middle), and carbon biomass (µgC.L^-1^) (top). Colors represent major eukaryotic and prokaryotic photosynthetic lineages defined using the psbO marker gene, integrated over the 0.2–2,000 µm plankton size range at surface stations from Tara Oceans, Tara Oceans Polar Circle, and Tara Pacific.

Compared to cell abundance, the predominance of green algae is less pronounced in the polar biome, where eukaryotic phytoplankton carbon biomass is more dominated by diatoms (Fig. 7, Fig. S16). Notably, several polar stations show significantly higher diatom biomass, with carbon concentrations reaching approximately ten times the average phytoplankton biomass in the polar biome (Fig. 7). At these *Tara* sampling sites (stations 183, 188, and 163), diatom blooms were documented through imaging data as well as sampling logsheets, particularly involving medium to large-sized diatoms in size fractions greater than 20 μm. This shift in lineage contribution, compared to cell abundance-based community composition, was partly expected, as our biomass estimates specifically accounted for the larger and more variable cell sizes of diatoms. Still, the predominance of diatom carbon biomass observed at polar stations aligns with previous results on their biogeography, as well as their prominent role in biogeochemical cycles and carbon uptake in high-nutrient concentration regions (*86*).

This latitudinal gradient of phytoplankton biomass only includes stations for which we could convert carbon biomass for all nine phytoplankton groups considered. Consequently, this representation does not include all sampling sites. For example, the maximum diatom carbon concentration that we estimated (>5,000 μgC.L⁻¹) appeared in the nutrient-rich upwelling waters off Chile (station 92) (Fig. S16).

Finally, converting cellular abundances into carbon biomass highlights the prominent role of the genus *Trichodesmium* within the photosynthetic community, particularly at low-and mid-latitude stations (Fig. 7, Fig. S16). At these stations, *Trichodesmium* carbon concentrations are significantly higher than those of *Synechococcus* and *Prochlorococcus*, and also exceed those of eukaryotic phytoplankton (Fig. 7). Again, some stations are not represented in this latitudinal gradient of phytoplankton biomass, notably those with high *Trichodesmium* carbon biomass in the Mozambique Channel (Fig. 6, Fig. S16).

### Distinct Quantitative Distributions of Diazotrophic and Non-Diazotrophic Trichodesmium

Building on the prominent contribution of *Trichodesmium* to phytoplankton carbon biomass observed at low- and mid-latitudes, we used a set of environmental genomes for this genus to refine its quantitative genomic distribution across *Tara* Oceans stations. We estimated seawater concentrations for five near-complete genomes reconstructed from large-size-fraction metagenomes from *Tar*a Oceans (*87*). Unlike most MAGs from the two collections mentioned earlier, these five genomes provide a fairly comprehensive representation of known *Trichodesmium* populations. A comparison with *psbO*-based abundance estimates confirmed that over 90% of *Trichodesmium* cells detected across *Tara* samples using the marker-gene approach were also represented by one of the five MAGs (Fig. S17).

Nitrogen-fixing *Trichodesmium* genomes (*20*) were found at only about half of the stations where *Trichodesmium* were detected. However, their mean and maximum genome concentrations exceeded those of non-diazotrophic *Trichodesmium* by approximately two orders of magnitude (Fig. 8). Highest concentrations of diazotrophic *Trichodesmium* were observed in the North Pacific Ocean, in the Red Sea, and in the South Indian Ocean. This distribution of nitrogen-fixing *Trichodesmium* mirrors the biogeography of *Trichodesmium* described from imaging data collected during *Tara* Oceans expedition (*36*). Direct comparisons between DNA-based and imaging-based abundance values remain limited due to variability in ploidy within lineages, as well as differences in the number of cells per filament and filaments per colony (*36*).

**Fig. 8.**
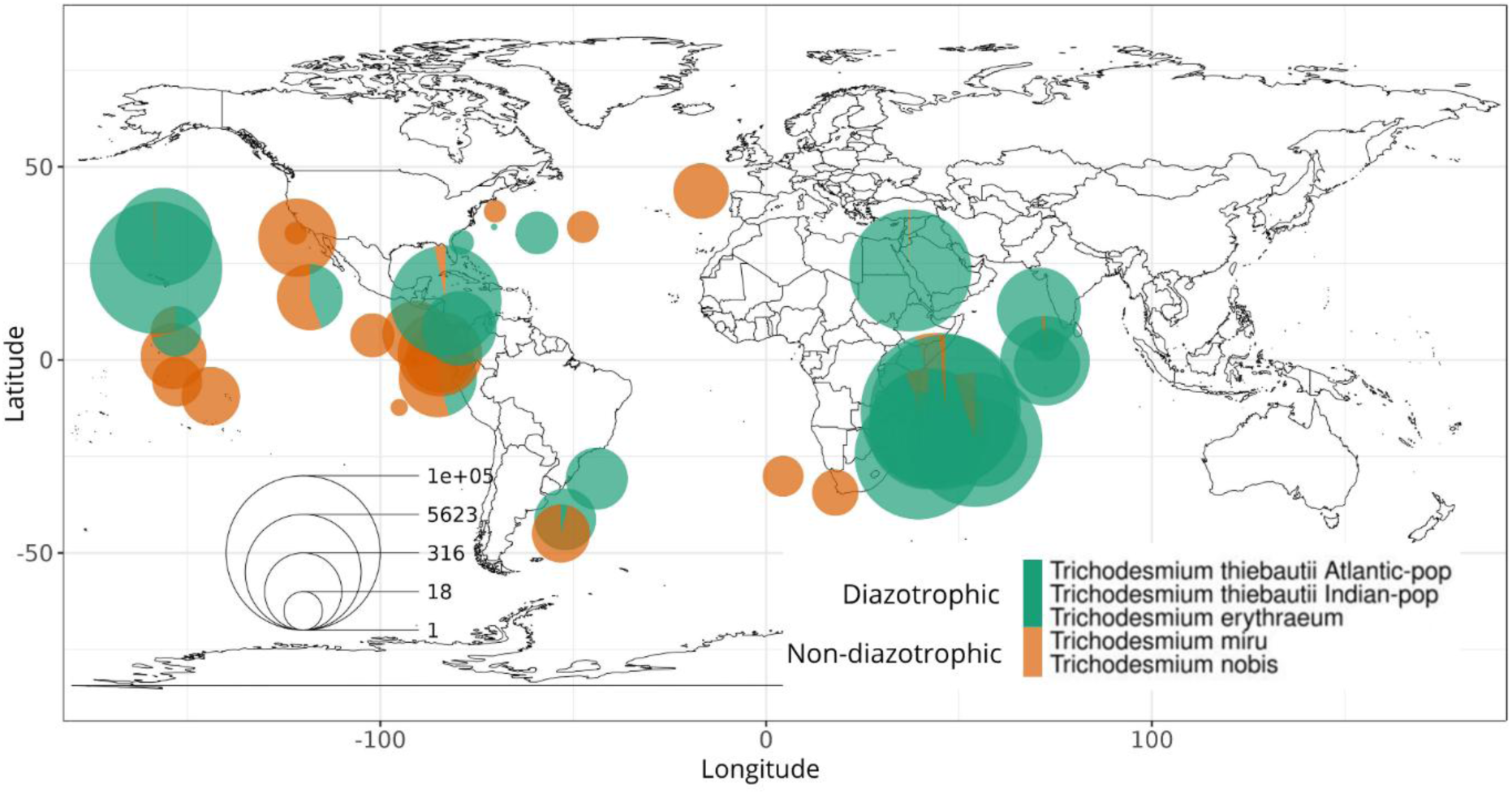
Genome-resolved quantification reveals higher abundance of diazotrophic (*nifH*-bearing genomes) compared to non-diazotrophic *Trichodesmium* across *Tara* sampling sites. The size of each pie chart indicates the total abundance (log10(genome copies.L^-1^)) of all Trichodesmium MAGs. Cumulative abundance of each MAG is calculated across all size fractions from 0.8 to 2,000 µm. Colors indicate diazotrophic (green) and non-diazotrophic (orange) Trichodesmium populations. Diazotrophic Trichodesmium populations correspond to MAGs carrying the nifH-gene.

Only one non-diazotrophic *Trichodesmium* genome (*Candidatus Trichodesmium nobis* (Fig. S18)) was detected at a few stations where nitrogen-fixing *Trichodesmium* were observed. As previously reported (*20*), diazotrophic and non-diazotrophic *Trichodesmium* genomes did not co-occurred in numerous stations across the Pacific and Atlantic Oceans, preventing more global abundance comparisons between nitrogen-fixing versus non-fixing populations using compositional metagenomic data.

Here, absolute quantification enables direct comparison of the two *Trichodesmium* groups across all 106 *Tara* stations analyzed and within the entire 0.8-2000 µm sampled community. At these stations, our estimates reveal that diazotrophic *Trichodesmium* genomes are more abundant than their non-nitrogen-fixing counterparts, providing a more nuanced interpretation than previously suggested by relative abundance data (*20*).

This analysis further demonstrates the complementary value of genome-resolved quantification of marine plankton. Compared to the *psbO*-based approach, genome-level quantification allows for the integration of additional genomic and functional information associated with this biological level—in this case, the nitrogen fixation function associated with the *nifH* gene in certain *Trichodesmium* genomes.

## Discussion

This study provides quantitative assessments of marine plankton in already-sequenced samples, enabling the leverage of valuable environmental datasets from past expeditions. Here, the abundance estimates of both eukaryotic and prokaryotic MAGs and *psbO*-lineages constitute a novel quantitative resource associated with the large-scale metagenomic sampling of marine plankton by *Tara* Oceans, *Tara* Oceans Polar Circle and *Tara* Pacific expeditions.

Based on this resource, we show how quantifying environmental metagenomic data can improve current understanding of the composition, function and biogeography of marine plankton for both species and communities, typically inferred from compositional omics data. We use the *psbO* marker gene to provide a quantitative overview of phytoplankton communities across *Tara* stations, including estimations of cell concentrations and carbon biomass in surface seawater for the main eukaryotic and prokaryotic photosynthetic lineages. We note the critical importance of extending quantification all the way to carbon biomass estimation in order to accurately assess the structure of phytoplankton communities that span a large range of size classes, as evidenced by substantial differences between latitudinal compositional patterns based on cell abundance versus carbon biomass.

This *psbO*-based approach also enables us to map global biogeographies of phytoplankton cell abundance with consistent taxonomic resolution across lineages, and at a finer level than model predictions based on imaging observations or pigment signatures. On a global scale, observations and projections reproduced major known ecological patterns, including the dominance of prokaryotes in oligotrophic subtropical regions and the prevalence of eukaryotic phytoplankton at higher latitudes. At a finer genomic resolution, quantification of eukaryotic and prokaryotic environmental genomes enable us to compare relative and absolute abundance-based co-occurrence networks. Overall, these analyses show significantly more positive associations, including a greater number of Bacteria-Eukaryote interactions, within plankton communities inferred from absolute abundance networks. Importantly, Bacteria-Eukaryote interactions identified from the absolute abundance network are resolved at finer taxonomic levels and with greater confidence compared to those derived from relative abundance data, confirming the value of genome-resolved quantification to uncover novel biotic interactions among planktonic taxa.

The value of genome-resolved quantification is further demonstrated by leveraging functional annotations of *Trichodesmium* genomes to refine the quantitative biogeography of diazotrophic *Trichodesmium* populations. This approach reveals the dominance of nitrogen-fixing *Trichodesmium* across *Tara* Oceans surface stations, providing insights that were previously inaccessible through metagenomic compositional analyses or conventional imaging and microscopy methods.

However, several biases limit the interpretations of the results, as our approach does not account for taxon-or sample-specific biases that may distort community compositions (*88*, *89*). Metagenomic relative abundances used in this analysis may be affected by uneven performances of environmental sampling, DNA extraction, library preparation and shotgun sequencing across targeted planktonic lineages. At first consideration, these issues may, to some extent, hinder reliable comparisons between taxa within the same sample. However, if compositional metagenomic abundances were too biased to reasonably represent true community structure, this would call into question the validity of most studies based on metagenomic profiling of communities, except those using a presence/absence approach (*31*). Nevertheless, numerous metagenomic studies, including those based on *Tara* Oceans samples used in this study, have shown strong correlations with imaging and optical measurements of relative community composition (*9*, *90*, *91*), which likely explains the significant associations observed with environmental variables (*55*, *92*).

In addition to compositional biases, other uncertainties in the absolute quantification of planktonic species arise from underestimating initial DNA concentrations in seawater. Bei et al. (*34*)’s results suggest that efficiency losses occur mainly during DNA purification rather than extraction, as their metagenomic-based estimates of phytoplankton abundances closely matched flow cytometry counts when internal standards were added after DNA extraction but prior to purification.

A relative error in the initial DNA concentration estimates is not critical for large scale comparisons, provided it remains relatively stable across samples and the different orders of magnitude of actual DNA concentrations in seawater. In such cases, applying a constant correction factor could align estimates with “ground-truth” abundances. However, uncorrected DNA concentration estimates would still consistently reflect variations in plankton community biomass or abundance. Here, assessing the consistency of this relative error is challenging, as the relationship between organism biovolume and DNA content needs to be better characterised at the community level. The non-linear relationship observed between community biovolume and DNA content (Fig. 1) does not necessarily mean that DNA quantification efficiency changes depending on *in situ* concentrations. Such patterns are expected at the community level due to multiple power-law relationships between DNA content and cell volume at the species scale (*93*, *94*). Notably, distinct power laws have been reported across various eukaryotic and prokaryotic lineages, including multicellular and unicellular phytoplankton (*95*, *96*), as well as marine bacteria and archaea (*97*). Beyond potentially irregular efficiencies, the complexity of planktonic communities and the presence of multicellular organisms may also contribute to deviations from these lineage-specific scaling laws linking biovolume and DNA mass.

That said, the relative error between DNA-based estimates of cell concentrations and flow cytometry counts remains stable across several orders of magnitude (Fig. 1). This stability reflects substantial consistency in DNA quantification efficiency across analysed samples and emphasizes the importance of adopting standardised protocols. Therefore, despite underestimating absolute concentrations and carbon biomass, the proposed approach yields representative and reliable estimates of variations across sampling sites, thereby supporting its use in eco-biogeographical quantitative assessments at both the lineage and genome scales.

Extending this quantitative framework to global scales demonstrates that absolute abundance estimates derived from metagenomic data can serve as a robust empirical foundation for predictive geographical and quantitative distribution modeling. Using these quantitative estimates, ensemble projections based on key environmental predictors revealed coherent and ecologically meaningful biogeographical patterns of major phytoplankton groups. These projections captured known ecological niches and large-scale gradients in temperature and nutrient supply, underscoring the reliability of abundance-based approaches. Variable-importance analyses identified sea surface temperature, phosphate and iron as the strongest predictors across taxa, consistent with their established roles in controlling marine productivity and phytoplankton distributions. Model uncertainty was primarily driven by the geography of the training data rather than algorithmic instability, with Mahalanobis distance analyses indicating higher extrapolation risk in under-sampled regions such as the Southern Ocean and coastal upwelling systems. While some methodological and sampling biases may influence the quantitative estimates, and hence the resulting projections, the overall consistency of large-scale patterns highlights the strength and scalability of this framework.

Altogether, this study advances a scalable and transferable framework for translating genomic data into quantitative ecological information, providing new means to quantify plankton community structure and function across ocean basins. By linking molecular diversity to biogeochemical and environmental gradients, this approach lays the groundwork for integrating plankton diversity into biogeochemical and Earth system models, ultimately contributing to a more mechanistic understanding of ocean productivity and global carbon cycling.

## Materials and Methods

### Resource Dataset of DNA Concentrations Across Multiple Ocean Basins

#### DNA quantification

We collected masses of extracted DNA from *Tara* Oceans, Polar Circle and *Tara* Pacific genomic samples (*40*, *43*). For *Tara* Oceans, Polar Circle and *Tara* Pacific expeditions, the handling of omics samples and nucleic acids extraction protocols are detailed in Alberti et al. (*44*) and Belser et al. (*45*), respectively. Overall, we retrieved values of DNA quantities extracted from filters for 599 *Tara* Oceans and Polar Circle surface samples - from station 4 to 210, covering size fractions from 0.2 to 2,000 µm. Data on DNA quantities extracted from filters were available for 1,420 *Tara* Pacific Ocean-Atmosphere (OA) plankton samples covering all size fractions. More details are presented in *Supplementary Materials*.

For the *Tara* Pacific expedition, multiple replicates were collected per site and size fraction. In 11 cases, the specific replicate used for sequencing was not identified. The replicate with the highest estimated DNA concentration was then selected.

#### Determination of Filtered Seawater Volumes

Data used to compute filtered water volumes were manually retrieved from various log sheets filled on board. “Event” log sheets (*40*, *43*) consign details on every single deployment of sampling devices used to collect seawater. Full registries of *Tara* Oceans and *Tara* Pacific event log sheets are available on Pangaea at https://doi.pangaea.de/10.1594/PANGAEA.842227 and https://store.pangaea.de/Projects/TARA-PACIFIC/Logsheets/, respectively. After water collection, each step of the protocols leading to the final genomic samples is detailed in “samples” or “WETLAB” log sheets (*40*, *43*). All sample registries for *Tara* Oceans and *Tara* Pacific are available at https://doi.pangaea.de/10.1594/PANGAEA.875580 and https://store.pangaea.de/Projects/TARA-PACIFIC/Logsheets/, respectively.

For each sample, the effective filtered volume was calculated using the formulas presented in Table 1, which account for the specific protocol steps associated with each size fraction. Water collection procedures and filtration protocols for each size fraction of *Tara* Oceans, Polar Circle, and *Tara* Pacific are fully detailed in Pesant et al. (*40*) and Lombard et al. (*43*), respectively.

For a given sample *j* (a specific *Tara* station and size fraction) the corresponding DNA concentration in seawater (C_totj, DNA_ (ng.L^-1^)) was calculated by dividing the total DNA mass extracted from the filter by the effective volume of seawater (V) processed to obtain the sample.

### DNA-based quantification at genomic scale using metagenomic data

#### Computing genomes or genes DNA concentrations in sample

The rationale for this quantification approach lies in the underlying assumption that, given reasonable sequencing depth, the metagenomic relative abundance of a genome or gene directly reflects its proportion in the sample’s total DNA mass, as the following relationship:

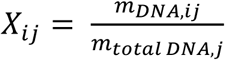

where m_DNA, ij_ (ng) is the mass of DNA corresponding to target *i* in sample *j*

And *X_ij_* is the metagenomic relative abundance of a target *i* in a metagenome *j*

Based on this assumption, the total DNA mass extracted from each sample (m_total DNA_) was used as a quantitative “anchor” (*30*) to convert metagenomic relative abundances into absolute DNA concentrations of genomes or reference sequences within that sample. Incorporating volumes of filtered seawater allows for true *in situ* (or “across-sample”) quantifications, rather than relying only on masses of DNA for post-sequencing, “within-sample” normalization (*14*, *98*) :

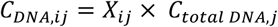

where C_DNA, ij_ is the DNA concentration in seawater of target *i* for station and size fraction defined by sample *j*

This calculation does not account for potential biases related to taxon and community-composition, which may be introduced by water sampling and filtration, DNA extraction, library construction and shotgun sequencing (*88*). These putative biases are detailed further on in the *Supplementary Materials*.

#### Estimates of cell concentrations in seawater

The final step in the quantification workflow consists of computing absolute cell abundances based on the DNA concentrations of specific targets in seawater. Cell densities were estimated only when both genome or gene length and the taxon-specific ploidy were known for a given target. When possible, cell abundances in seawater were calculated as follows:

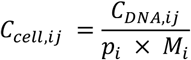

where p_i_ is the ploidy of organisms represented by target *i*

M_i_ (ng) is the mass of target *i* (genome or gene) computed with the formula: 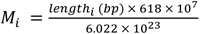

This rationale was first applied by Pierella Karlusich et al. (*36*) to estimate *Trichodesmium*-associated *nifH*-gene concentrations in seawater based on the quantification of extracted DNA. Only the formula used to calculate genome or gene mass differed from this study.

### Metagenomic data from *Tara* expeditions

#### Eukaryotic and prokaryotic genomes collections

Genome-level quantification across *Tara* stations was performed separately for eukaryotic and prokaryotic genomes. For eukaryotes, we used results from the previous mapping of 939 metagenomes, spanning all size fractions of *Tara* Oceans and *Tara* Oceans Polar Circle samples, against a collection of eukaryotic genomes (*38*). For prokaryotes, 180 metagenomes from the bacteria-enriched size fractions (0.2-1.6 µm and 0.2-3 µm) of *Tara* Oceans and *Tara* Oceans Polar Circle were mapped against a recently compiled prokaryotic genomes database (*26*, *37*).

The eukaryotic collection comprises 713 metagenomes reconstructed genomes (MAGs) and single-amplified genomes (SAGs) that were reconstructed and manually curated by Delmont et al. (*38*) from eucaryote-enriched size fractions of *Tara* Oceans samples. The prokaryotic genomic database is composed of the species-level genomes collection from Giordano et al. (*26*) and the 530 Arctic MAGs from Royo-Llonch et al. (*37*). The compilation of the genome collections and the steps of metagenomic mapping are outlined in the *Supplementary Materials*.

To convert DNA concentrations into cell concentrations, specific filtering criteria were applied to select genomes: either a completion rate exceeding 70%, or inclusion among the top 50% of most abundant genomes with a minimum completion threshold of 20%. This filtering step resulted in 318 eukaryotic genomes and 5,152 prokaryotic genomes. DNA concentrations were converted into cell concentrations for all prokaryotic genomes using a ploidy value of 1. Cell concentrations were calculated for the 223 eukaryotic MAGs which were classified into taxa for which ploidy values could be approximately inferred from the literature. The following ploidy values were applied : *Mamiellophyceae*, *Phaeocystis*, and *Ascomycota* : 1 ; *Bacillariophyta* and *Pelagomonadaceae* : 2 ; *Animalia* : 2 ; and *Prymnesiophyceae* : 1.5 (to account for both haploid and diploid stages described in the literature (*99*)).

#### Trichodesmium MAGs

The five *Trichodesmium* MAGs used here were reconstructed in a previous study (*20*). Detailed methodologies for *Trichodesmium* MAG reconstruction and metagenomic mapping are provided in Delmont et al. (*20*, *87*).

#### *psbO* marker gene

We used the *psbO* reference sequence database compiled by Pierella Karlusich et al. (*9*), available through the EMBL-EBI BioStudies repository under accession S-BSST659. This comprehensive database includes over 17,000 *psbO* gene reference sequences from a broad range of marine phytoplankton and terrestrial plant lineages.

We used this reference collection to recruit metagenomic reads from 1,246 metagenomes from the *Tara* Oceans and Polar Circle expeditions, and 227 metagenomes from the *Tara* Pacific expedition. Read mapping was performed using the Burrows-Wheeler Aligner (BWA) tool (*100*), with an identity threshold of 95%. No detection limit was imposed on the number of reads recruited by a given reference sequence. Among the entire database, 9,616 and 4,660 *psbO* reference sequences were detected in *Tara* Oceans–Polar Circle and *Tara* Pacific metagenomes, respectively, for which corresponding DNA concentrations were available. After converting DNA concentrations into *psbO* gene copy numbers per liter of seawater, concentrations below 1 gene copy.L^-1^ were rounded down to zero.

Based on prior taxonomic annotations (*9*), *psbO* sequences were grouped into 15 photosynthetic lineages encompassing both eukaryotic and prokaryotic taxa : *Prochlorococcus* (n = 1,239), *Synechococcus* (n = 629), *Trichodesmium* (n = 14), *Richelia* (n = 4), *Calothrix* (n = 2), *Croccosphera* (n = 6), other *cyanobacteria* (n = 21), *Chlorophyta* (n = 552), *Pelagophyceae* (n = 368), *Dinophyceae* (n = 2,114), *Bacillariophyceae* (n = 1,078), Prymnesiales (n = 588), Phaeocystales (n = 282), non-Prymnesiales Haptophytes (n = 1,438), and other eukaryotes (n = 847). Haptophytes were divided into three categories with unequal taxonomic resolution to isolate a group of non-Prymnesiales haptophytes mainly composed of calcifying organisms, mostly from the *Coccolithales* and *Isochrysidales* orders.

For each lineage, cell abundances were calculated using the following ploidy values : all eukaryotic lineages except Chlorophytes : 2; Chlorophytes : 1; *Trichodesmium*: 5 (*36*) ; *Synechococcus*, *Prochlorococcus*: 1 (*101*) ; for all other cyanobacterial lineages : 2 (*102*).

### Co-occurrence network inference and validation

Based on both the relative abundance matrix and the absolute abundant matrix derived from cell concentrations (with matching stations, size fractions and MAGs), we first removed MAGs with less than ten observations across samples in each matrix independently. This step aims to reduce the incidence of spurious correlations in subsequent network construction. This filtering step resulted in abundance matrices containing 4,118 prokaryote and 111 eukaryote MAGs across 56 metagenomes, which were used as input for co-occurrence network reconstruction. The relative abundance matrix was transformed using the centered log-ratio (CLR) to deal with the compositional constraints of sequencing data. Co-occurrence networks were then reconstructed with *FlashWeave* (*103*) independently for both matrices using the following parameters: n_obs_min=10, max_k = 0, heterogenous=true.

We benchmarked the robustness of the co-occurrence networks reconstructed from the relative and the absolute abundance matrices by comparing the predicted correlations against known biotic interactions registered in the GLOBI database v0.8 (*60*). All possible matches between co-occurring MAG pairs and documented interactions in the database were considered at both genus-genus and genus-family taxonomic levels. Then, for each taxonomic rank pair, we calculated the percentage of predicted correlations that could be validated by determining the proportion of edges supported by known interaction relative to the total number of co-occurring MAG pairs that could be classified at the relevant ranks.

### Environmental Data

Environmental data used in correlation analyses to PCoA ordination axes of the polar and non-polar communities was obtained from https://doi.pangaea.de/10.1594/PAN and from Delmont et al. (*87*) and Chaffron et al. (*104*).

### Imaging, optical and flow cytometry data

Several imaging and optical datasets were combined to be used as quantitative controls of plankton biomass or abundance across *Tara* stations. Detailed protocols to collect water and retrieve imaging samples from *Tara* Oceans, Polar Circle and *Tara* Pacific stations are described in Pesant et al. (*40*) and Lombard et al. (*43*). Depending on the targeted size fraction of organisms, different quantitative imaging and optical devices were used.

For *Tara* Oceans and Polar Circle, the resulting datasets of biovolume of organisms per water volume are described in Lombard et al. (*50*). We used the “Meta-Plk > 20 µm” dataset (*50*) (station 4 to 210), which includes : environmental High Content Fluorescence Microscopy (eHFCM) data from 20 µm net filtrations Colin et al. (*90*) ; ZooScan biovolumes from >200 µm, >300 µm, and >680 µm filtrations (*105*) ; and Underwater Vision Profiler (UVP) (*106*). Biovolumes for organisms <20 µm or between 180–200 µm were estimated by interpolation and extrapolation to match genomic size ranges. Biovolumes were aggregated by size fraction corresponding to *Tara* genomic sampling.

For *Tara* Pacific, imaging datasets are described in Mériguet et al. (*51*), along with the processing of imaging samples and methods for biovolume measurements. Biovolumes were measured using a FlowCam (Fluid Imaging Technologies) (*107*) on samples originating from surface (0–1 m) seawater filtered through a 20 µm Deck-Net and a 200 µm sieve of collected material. Surface seawater was collected using a water-pumping system equipped with a flowmeter to record flow rates and assess filtered volumes (*43*). A High Speed Net (HSN) with a 330 µm mesh collected larger surface plankton (300–2,000 µm). HSN samples were analysed with a ZooScan system to estimate biovolumes (*50*). Unlike *Tara* Oceans and Polar Circle, for which genomic and imaging samples were collected separately, the same water masses were filtered simultaneously through the HSN to collect samples for both genomic and imagery 300-2,000 µm protocols. Deck-Net-FlowCam (*108*) and HSN-ZooScan (*109*) final datasets consist of aggregated biovolumes of organisms per water volume across 212 and 205 *Tara* Pacific OA stations, respectively.

Notably, Mériguet et al. (*51*) demonstrated the greater accuracy of theoretical filtered seawater volumes estimated from the boat’s trajectory, speed, and HSN features, likely due to flowmeter deficiency during high-speed towing. Despite improved accuracy of theoretical estimates, we chose to use the flowmeter-derived volumes to maintain consistency with the methodology previously applied to calculate DNA concentrations for other *Tara* Pacific size fractions and for *Tara* Oceans samples.

Flow cytometry (FC) counts complemented the previous imaging dataset for smaller size fractions. *Tara* Oceans and Polar Circle FC samples were collected by filtering water through a 10 µm mesh and analysed by a FACScalibur flow cytometer (*53*) to measure abundances and biovolumes of heterotrophic bacteria, picocyanobacteria (*Synechococcus, Prochlorococcus)* and some pico- and nano-eukaryotes. We calculated cumulative abundances and biovolumes for the 0.2-3 µm and 0.8-5 µm size fractions at both surface and DCM by aggregating cell counts and biovolumes of each group of prokaryotes and eukaryotes, based on observed average cell sizes. *Tara* Pacific FC samples (*52*) were collected by filtering seawater through a 20 µm mesh and analysed using a FACS Canto II flow cytometer (*43*). Cell counts of heterotrophic bacteria, picocyanobacteria (*Synechococcus, Prochlorococcus)* and pico- and nano-eukaryotes were compared directly to abundance estimates in the 0.2-3 µm genomic fraction, as average cell size measurements were unavailable.

### Summation Criteria and Station Coverage for Community Cell Abundance

To compare relatively homogeneous communities across stations (spanning similar organism size ranges), total cell abundances were estimated only at stations where enough size fractions were sampled to represent the targeted phytoplankton lineages. Selection criteria were defined based on organism size and cell abundance distributions of each lineage across size fractions (Fig. S7).

For the smallest organisms (*Prochlorococcus*, *Synechococcus*, and unclassified cyanobacteria), cumulative concentrations were calculated only at stations where the bacteria-enriched size fraction (0.2–3 µm) was available. For all other lineages, total concentrations were calculated at *Tara* Pacific and Polar Circle stations when both the 0.2-3 µm and 3-20 µm fractions were sampled, and at *Tara* Oceans stations when the 0.8-5 µm fraction was available. When these conditions were met, cell concentrations from all size fractions, from 0.2 or 0.8 µm to 2,000 µm, were summed.

This distinction between cyanobacteria and other lineages is particularly important for the *Tara* Oceans dataset. It prevents summing cell abundances across overlapping (0.2-3 µm and 0.8-5 µm) or discontinuous (0.2-3 µm and 5-20 µm) size fractions, while ensuring that cyanobacterial abundances are not underestimated by excluding the 0.2-3 µm fraction.

### Conversion of Cell Abundances of Photosynthetic Lineages into Carbon Biomass

To convert cell abundances of phytoplankton lineages into carbon biomass, we used two distinct approaches depending on the availability of trait information for each taxonomic group. When available, we used lineage-specific average cellular carbon content values reported in the literature. For groups lacking such average values or known for significant cell size variability, we used conversion equations from the literature. These equations estimate cellular carbon content as a function of cell volume and incorporate lineage-specific parameters reflecting elemental stoichiometry and cellular composition (*110*). The method used for each group, either equation-based or using average values of cellular carbon content, is detailed in Table 2.

When conversion equations were applied, carbon biomass was calculated for each size fraction and then summed across fractions. For non-Prymnesiale Haptophytes, Prymnesiales, the cyanobacteria genus *Crocosphaera*, and the group of unidentified cyanobacteria, a standard cell size was assigned to each lineage according to the literature and applied regardless of the size fraction considered (Table 2). In contrast, for diatoms and dinoflagellates, the lower bound of each size fraction was used as the standard cell diameter to calculate average cell volume, using theoretical spherical cells. This distinction accounts for the greater variability in cell sizes observed within these two phytoplankton lineages. In cases when average cellular carbon content values were used, carbon biomass of the respective lineages was calculated from total cell abundances spanning size fractions from 0.2 (or 0.8) to 2,000 µm.

### Modeling Phytoplankton Abundance From Environmental Predictors

#### Data Preparation and Feature Engineering

We compiled a global dataset combining our estimates for phytoplankton abundances, based on the *psbO* gene (see corresponding section), with matched environmental variables. Each record corresponds to a sample at a given station and size fraction, with associated metadata (temperature, salinity, nutrient concentrations, chlorophyll-*a*, geographic coordinates). Phytoplankton communities were resolved into 16 taxonomic groups across the three target size classes used in *Tara* Pacific and *Tara* Oceans Polar Circle (0.2-2 µm, 2-20 µm, and 20-2,000 µm). To harmonize the dataset with *Tara* Oceans stations, which were originally sampled in fractions of 0.2-3, 0.8-5, 3-20, 20-180, 180-2000, and 20-2000 µm, we mapped these into the *Tara* Pacific classes. Specifically, the 0.2-2 µm class was taken from the 0.2-3 µm fraction or, when missing, from the 0.8-5 µm fraction; the 2-20 µm class was derived from the 3-20 µm fraction, and the 20-2,000 µm class was taken from the 20-2000 µm fraction or reconstructed by summing the 20-180 µm and 180-2000 µm fractions. Missing values for individual phytoplankton groups within any class were not imputed. Abundances were then aggregated across the three size classes to yield total group-level counts per station, while relative size-class contributions were retained as normalized proportions. All abundances were log₁₀-transformed to stabilize variance across orders of magnitude (111).

Environmental data were extracted from the World Ocean Atlas monthly composites (*112*) using the closest station-month match for temperature, salinity, phosphate, nitrate, and silicate. Chlorophyll-*a* (surface) was extracted from the OC-CCI monthly composites (*113*) using the same methodology. Dissolved iron (5 m depth) was added from the PISCES model (*24*). Latitude and longitude were retained for visualization but excluded as predictors. After filtering for complete records, the final dataset averaged 109 valid stations per taxon, reflecting variability in detection coverage across lineages. This limited coverage necessitated careful model design and sensitivity testing.

#### Model Training and Predictive Framework

We trained supervised regression models to predict log-transformed abundances from environmental features using bagging regression trees (114). This ensemble method reduces variance, is robust to overfitting, and captures non-linear relationships in limited, noisy datasets. Separate models were trained for each of 17 responses (16 groups + total phytoplankton), using only samples with complete predictors and targets. Hyperparameters (tree number, minimum leaf size) were tuned by Bayesian optimization (115), which efficiently explores parameter space and outperforms grid/random search under data scarcity (115). Optimal configurations were determined via 5-fold cross-validation, balancing predictive accuracy and data use. Performance was quantified using RMSE and R². Final models were retrained with all valid data and applied to the global prediction grid. Predictor importance was assessed using permutation importance based on out-of-bag (OOB) error estimates (116). This approach quantifies the contribution of each variable to prediction skill and provides ecological insights into environmental drivers of phytoplankton distributions. Variable rankings were summarized as mean OOB error increases across iterations, a method widely used in ecological modelling (116).

#### Global Prediction Grid

To enable global inference, we constructed a 1° × 1° monthly climatological grid excluding land and ice-covered areas. The seven predictors used in training were extracted for each grid cell and month, yielding more than 780,000 environmental vectors. These were reshaped into matrices matching model input, enabling monthly predictions of phytoplankton abundance across the global ocean.

#### Assessment of Environmental Similarity and Model Extrapolation

To evaluate extrapolation risk, we computed the Mahalanobis distance (*117*) between the predictor vector of each grid cell and the distribution of training environments. Distances were mapped onto the 1° × 1° grid for each month and averaged annually. A χ² distribution threshold (p = 0.95, df = 7) was used to flag grid cells lying outside the 95% confidence region of the training domain (*118*). These regions were considered extrapolation-prone, where predictions are less reliable.

#### Evaluation of Prediction Robustness and Sensitivity

Given the relatively limited size of our dataset for machine-learning approach (180 stations total; 109 valid stations per taxon), we tested the sensitivity of predictions to training set composition using a resampling-based robustness analysis (*119*). While final models used the full dataset, robustness tests systematically varied which subsets of *Tara* Pacific stations were included alongside a fixed *Tara* Oceans core. For each increment level, five independent models were trained with different random subsets, repeated for all groups. Each model generated global monthly abundance fields, which were averaged to annual means. Across repetitions, we computed mean and standard deviation of predictions at each grid cell, yielding consensus maps and uncertainty fields. High variance regions correspond to under-sampled or environmentally distinct areas, reflecting epistemic rather than aleatoric uncertainty. The strong spatial overlap between high-uncertainty zones, Mahalanobis exceedance masks, and resampling sensitivity confirmed that prediction confidence is fundamentally constrained by the geography and diversity of training data.

## Acknowledgments

We thank the LAGE (Laboratoire d’Analyses Génomiques des Eucaryotes, CEA) members for stimulating discussions on this project, the commitment of the Research Federation for the Study of Global Ocean Systems Ecology and Evolution (FR2022/TaraGOSEE). We would like to extend our warmest thanks to Alain Perret and Emilie Pataud for their time in conducting cultivation, Damien Eveillard for his insightful discussions on the prospects of this work, and Daniele Iudicone for sharing his vision and support. We also thank Serge Planes and all members of the Tara Pacific consortium for their support.

We thank all members of the Tara Oceans consortium for maintaining a creative environment and for their constructive criticism. Tara Oceans would not exist without the Tara Ocean Foundation and the continuous support of 23 institutes (https://oceans.taraexpeditions.org/).

This article is contribution number XX of *Tara* Oceans and number 51 of *Tara* Pacific

## Supplementary Materials

### Supplementary Text

### Materials and Methods

#### Resource Dataset of DNA Concentrations Across Multiple Ocean Basins

##### DNA quantification

For *Tara* Oceans and Polar Circle samples, DNA and RNA were simultaneously extracted from whole filter samples from size fractions > 0.8 µm as described under method ID Euk_DNA_RNA_ext in Alberti et al. (*44*). For the 0.2-3 µm size fraction, nucleic acid extractions were handled separately through metagenomics-or metatranscriptomics-targeting protocols. DNA was extracted from half filters as described under Acinas_Prok_DNA_ext method ID (*44*), and extracted DNA aliquots were sent to Genoscope. For method consistency, we calculated DNA quantities based on Acinas_Prok_DNA_ext elution volumes for extraction, volumes of aliquots sent to Genoscope.

For *Tara* Pacific, genomic samples from all size fractions designed for plankton sampling (S023, S320, S20 and S300) were processed at Genoscope following the same extraction protocol (*45*).

For all samples, DNA concentrations were estimated using a Qubit 2.0 Fluorometer at Genoscope.

#### DNA-based quantification at the species scale using metagenomic data

##### Eukaryotic and prokaryotic genomes collections

The methodologies employed for the reconstruction of eukaryotic MAGs and SAGs, along with the specifics of metagenomic mapping of *Tara* samples, are detailed in Delmont et al. (*38*).

The prokaryotic genomic database is a non-published resource comprised of 530 Arctic MAGs reconstructed from metagenomes of the bacteria-enriched size fraction of Polar Circle (*37*), combined with a collection of previously reconstructed whole-genome sequences, MAGs, and SAGs from marine prokaryotes curated by Giordano et al. (*26*).

The Arctic MAGs were incorporated into Giordano et al. (*26*) marine genome catalogue prior to the filtration of low-quality and redundant genomes. The de-replication process was conducted using dRep v2.2.3 with default parameters and a 95% ANI threshold. Detailed methods for quality checks and de-replication procedures are described in Giordano et al. (*26*). This process resulted in a total of 8,196 non-redundant species-level prokaryotic genomes with taxonomic assignments from GTDB-Tk v0.3.2. The curated genome collection was utilized to map quality-controlled reads from *Tara* metagenomes using Bowtie 2 v2.3.4.3.ds. Mapped reads were then filtered using Samtools v1.994 and pySAM v0.15.2, applying a MAPQ threshold of ≥ 20 and a 95% ANI criterion. For genomes with a horizontal metagenomic coverage above 30%, relative abundances were calculated based on the estimated number of reads mapping to each genome and the total number of sequenced reads.

For both eukaryotic and prokaryotic datasets, no additional detection thresholds were applied when estimating absolute DNA concentrations from relative genome abundances in the samples. It should be noted that MAGs included in this analysis were reconstructed using different assembling and binning methods and different identity thresholds were used to de-replicate databases of redundant genomes. Metagenomic mappings of *Tara* metagenomes were also conducted following different algorithms and recruitment criteria for the prokaryotic and the eukaryotic collections.

## Supplementary Figures and Tables

**Fig. S1.**
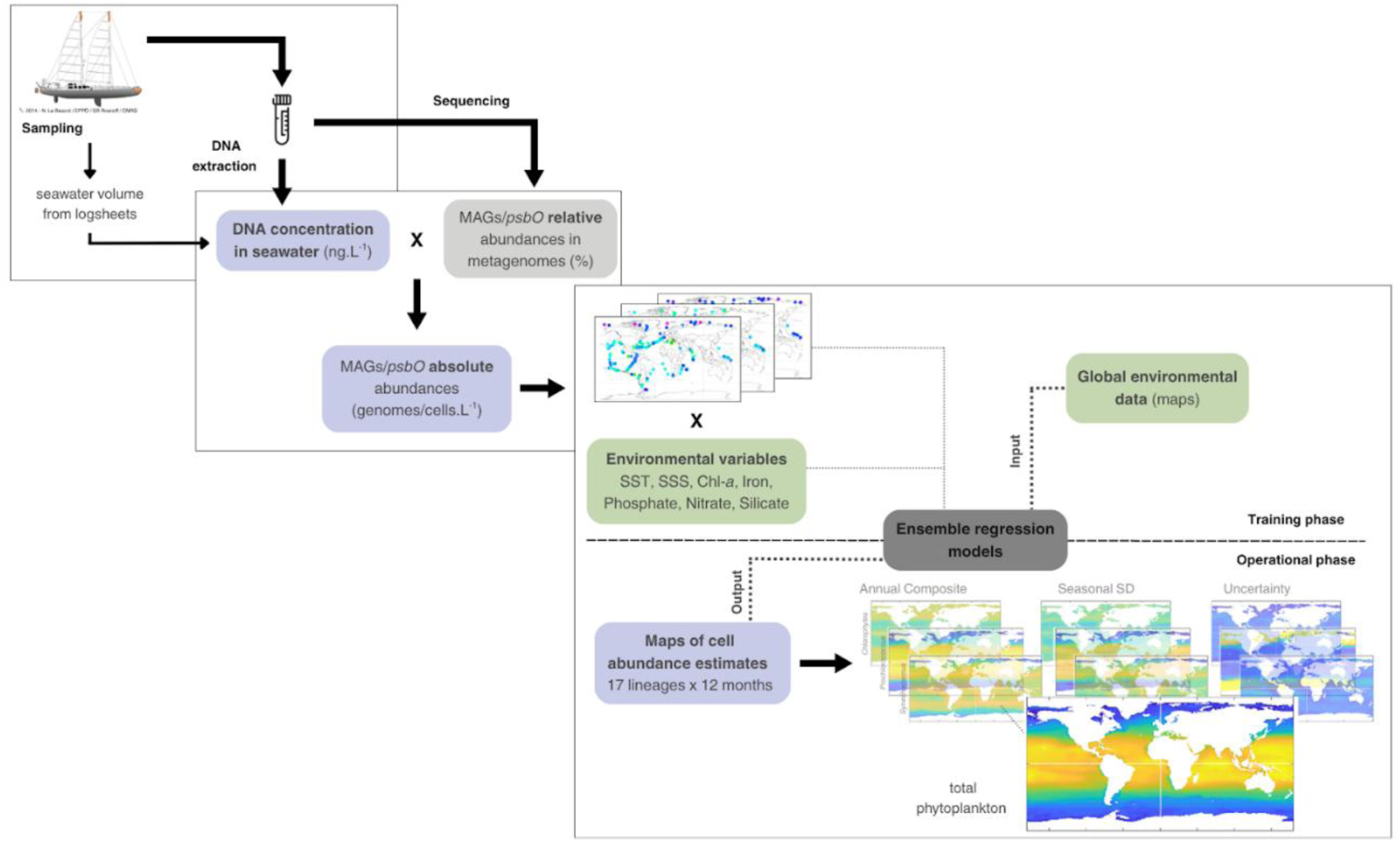
Flowchart describing the genomic-based quantification procedure and the framework for global phytoplankton abundance modeling. For each Tara station and size fraction, the corresponding DNA concentration in seawater (Ctotj, DNA (ng.L^-1^)) was calculated by dividing the total DNA mass extracted from the filter by the effective volume of filtered seawater (V) (Left). Estimates of DNA concentration in seawater (Ctotj, DNA (ng.L^-1^)) were used as absolute references to convert metagenomic relative abundances into absolute DNA concentrations corresponding to a specific genome or gene within samples. For each genome or gene, associated cell abundances were estimated based on their DNA concentrations in seawater and their length (Center). Finally, regression models were trained for each phytoplankton lineage using psbO-based estimates of total cell concentration pooled across the 0.2 to 2,000 µm size range and global environmental variables. These models provided global annual composite maps of phytoplankton cell abundance, as well as an evaluation of the seasonal variability and the robustness of the predictions (Right).

**Fig. S2.**
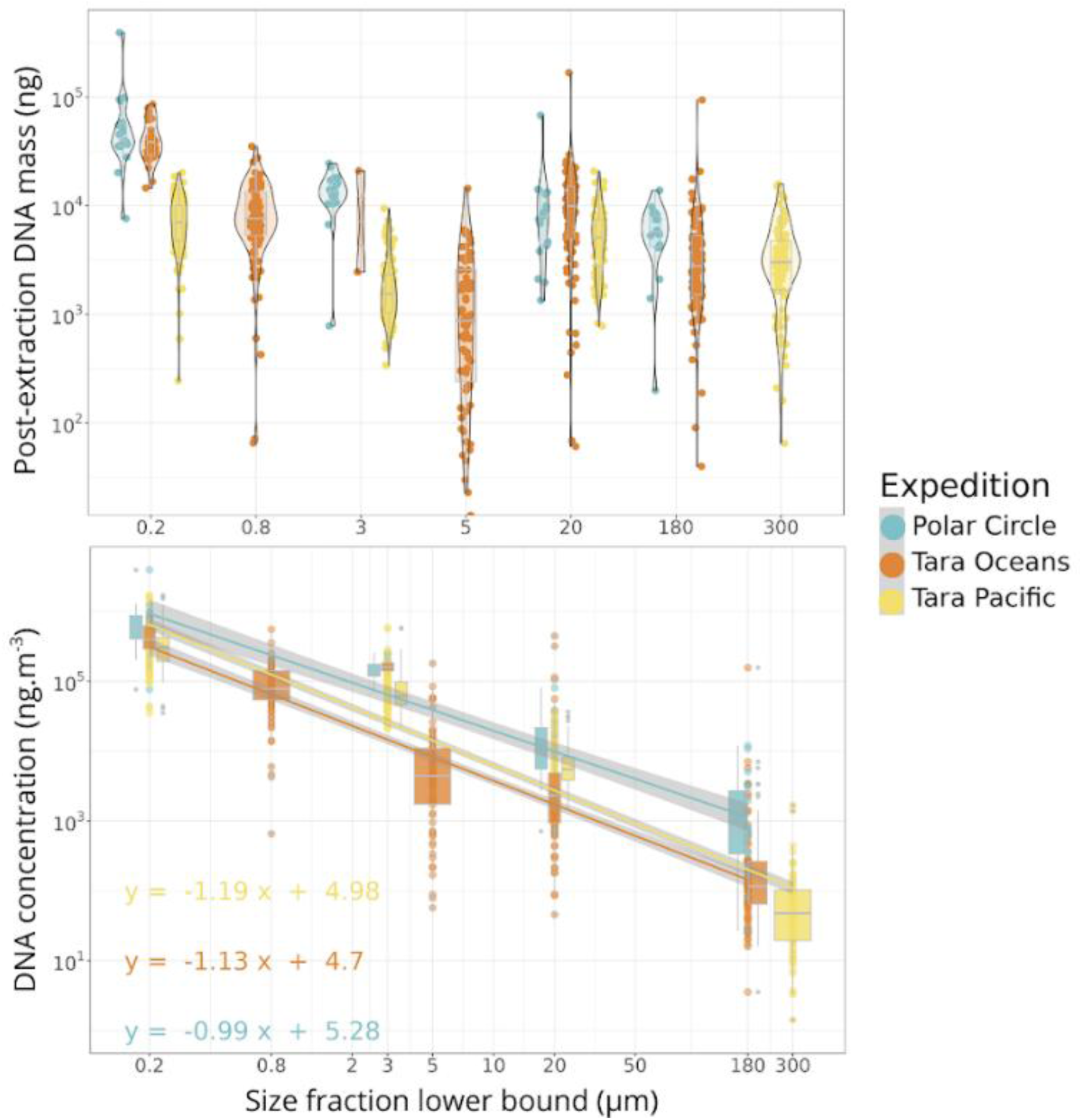
Post-extraction DNA quantification and DNA concentration estimates for *Tara* Oceans, Polar Circle and *Tara* Pacific samples from all size fractions.

**Fig. S3.**
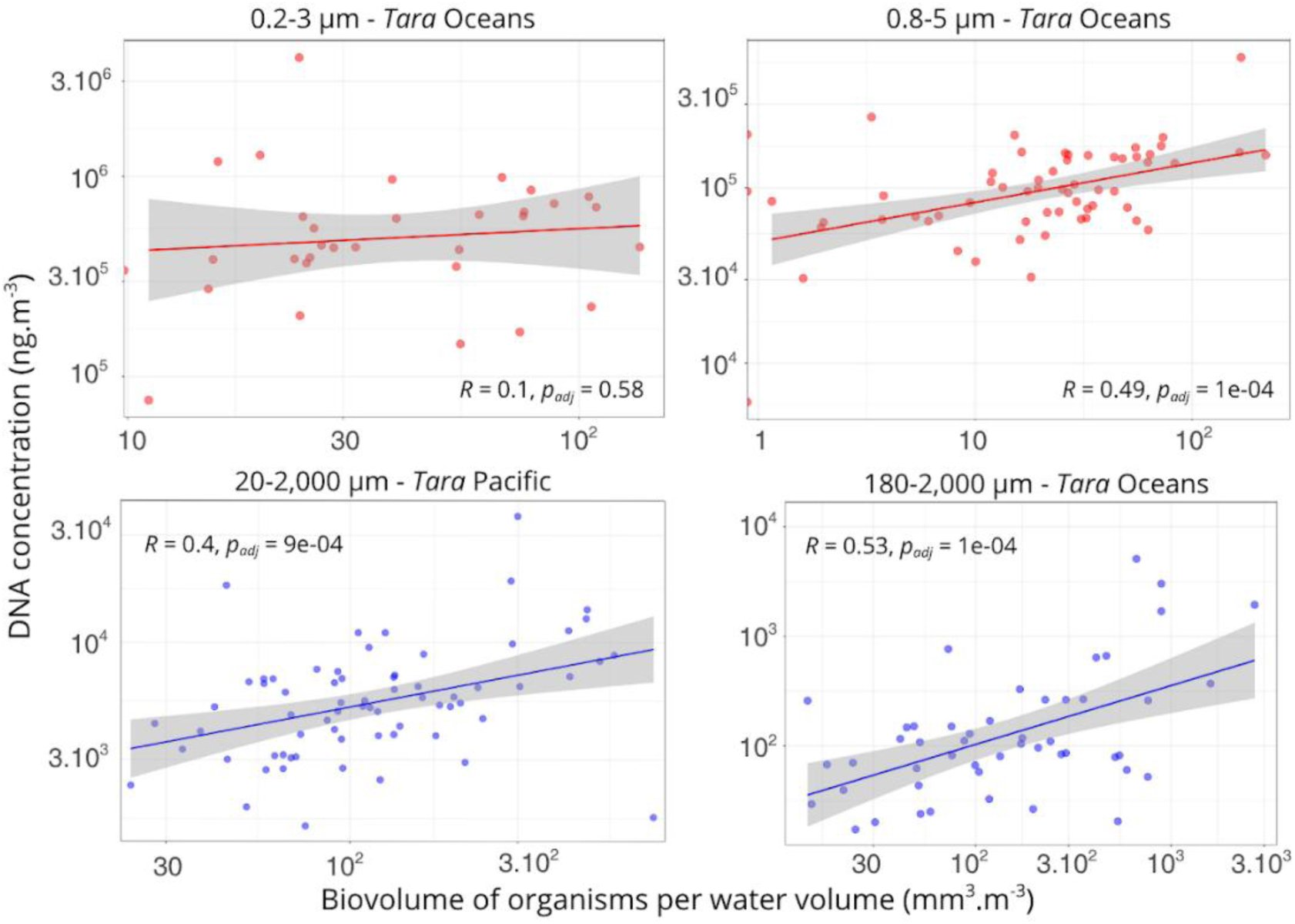
Estimates of DNA concentrations in seawater for *Tara* Oceans, Polar Circle and *Tara* Pacific samples compared to organism biovolumes per unit water volume, measured for the corresponding size fractions and sampling sites. Organism biovolumes were measured by flow cytometry (red) for the 0.2–3 μm and 0.8–5 μm size fractions, and by various imaging techniques (blue) for the other size fractions. Pearson correlation coefficients are indicated.

**Fig. S4.**
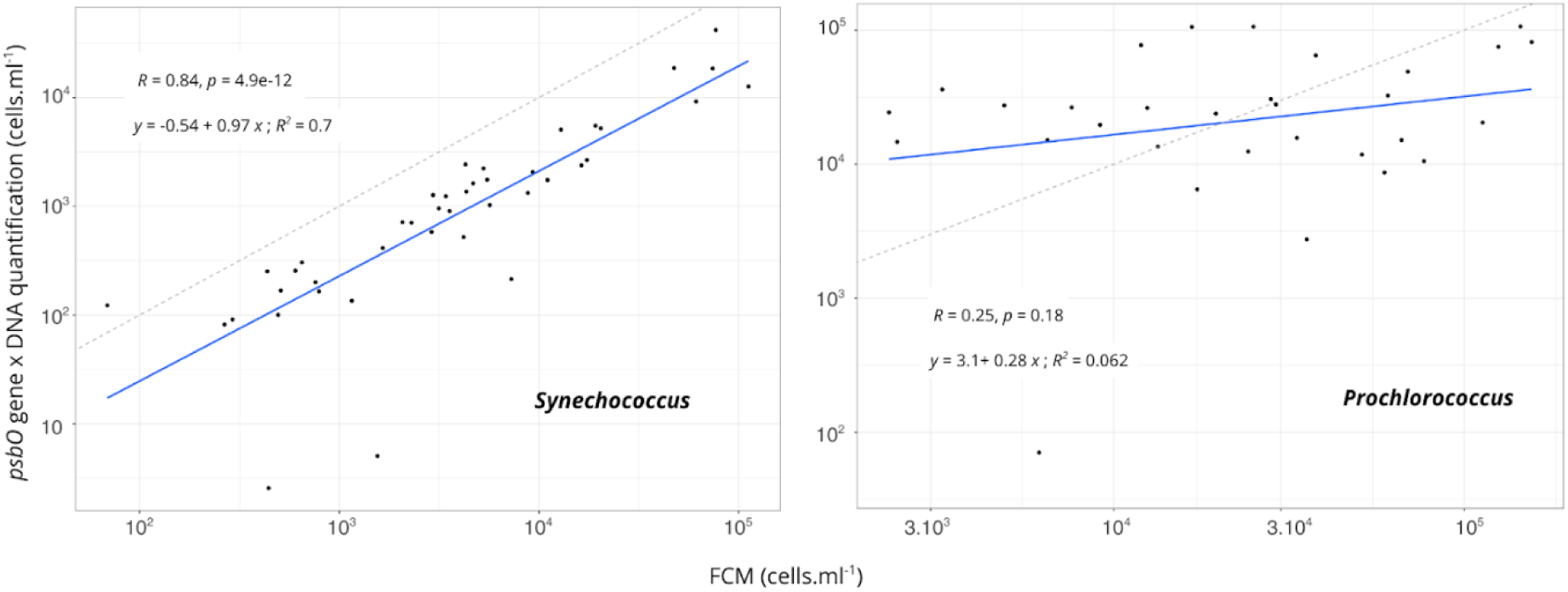
Cell abundances of the cyanobacteria *Prochlorococcus* and *Synechococcus* estimated from DNA-based quantification of the *psbO* marker gene versus flow cytometry cell counts across *Tara* Pacific stations. The y-axis shows cell concentrations estimated from psbO quantification in metagenomes from the 0.2-3 μm size fraction. The x-axis shows flow cytometry cell counts for the corresponding stations. Pearson correlation coefficients are indicated. Only flow cytometry data with quality scores between 3 and 5 are shown.

**Fig. S5.**
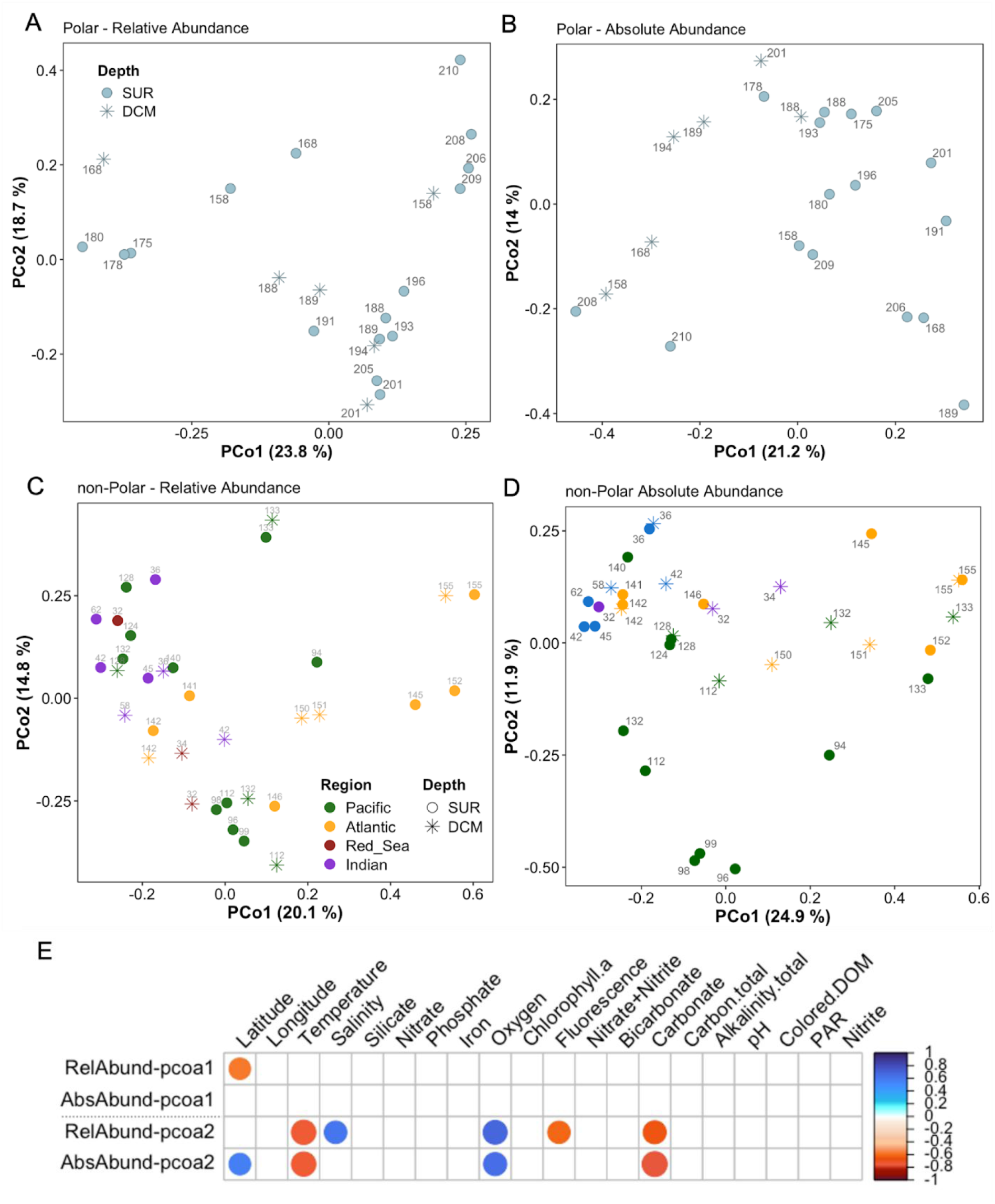
(A-D) Principal Coordinates Analysis (PCoA) ordination of Bray-Curtis similarity matrices for polar and non-polar eukaryote and prokaryote communities based on relative or absolute abundance estimates. **(E)** Correlation of PCoA axes with various environmental parameters for the polar community. White boxes indicate non-significant correlations (p > 0.01).

**Fig. S6.**
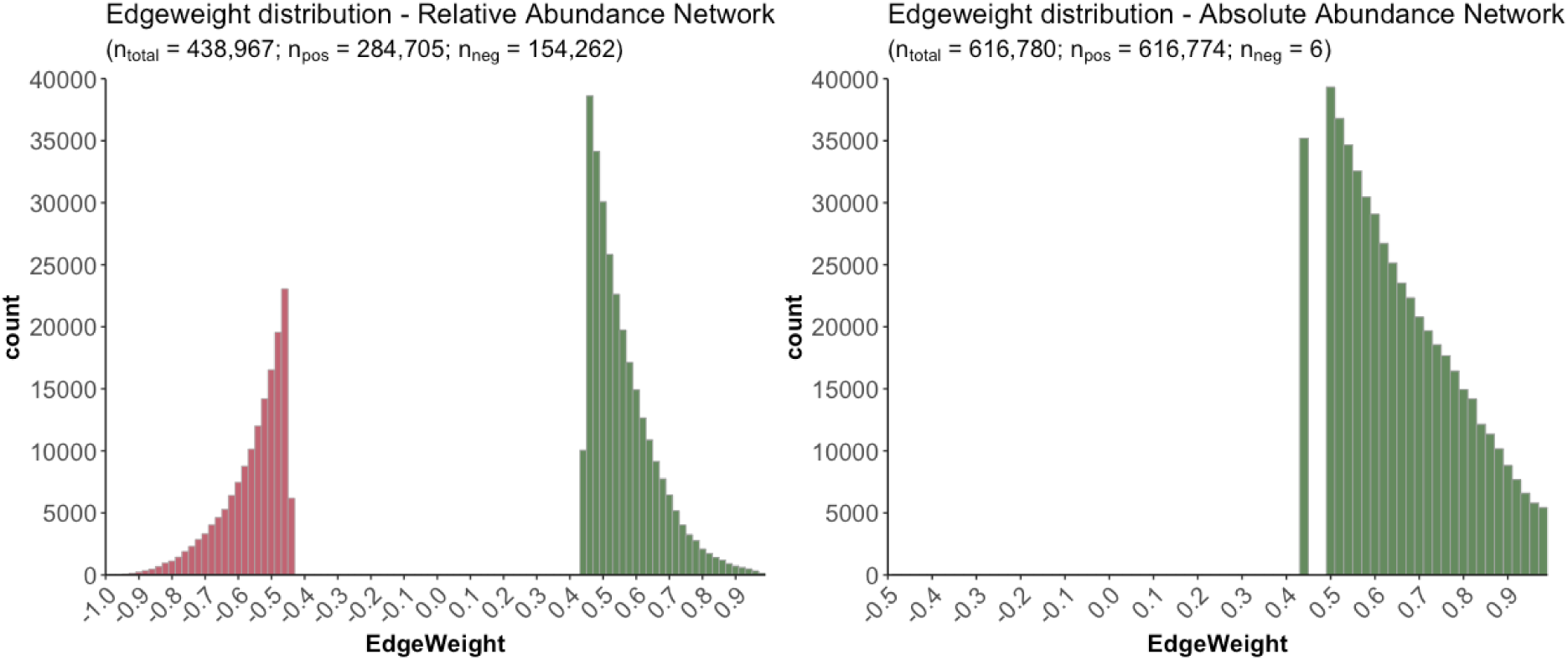
Histogram showing edge weight distributions of the relative (left) and absolute (right) abundance networks.

**Fig. S7.**
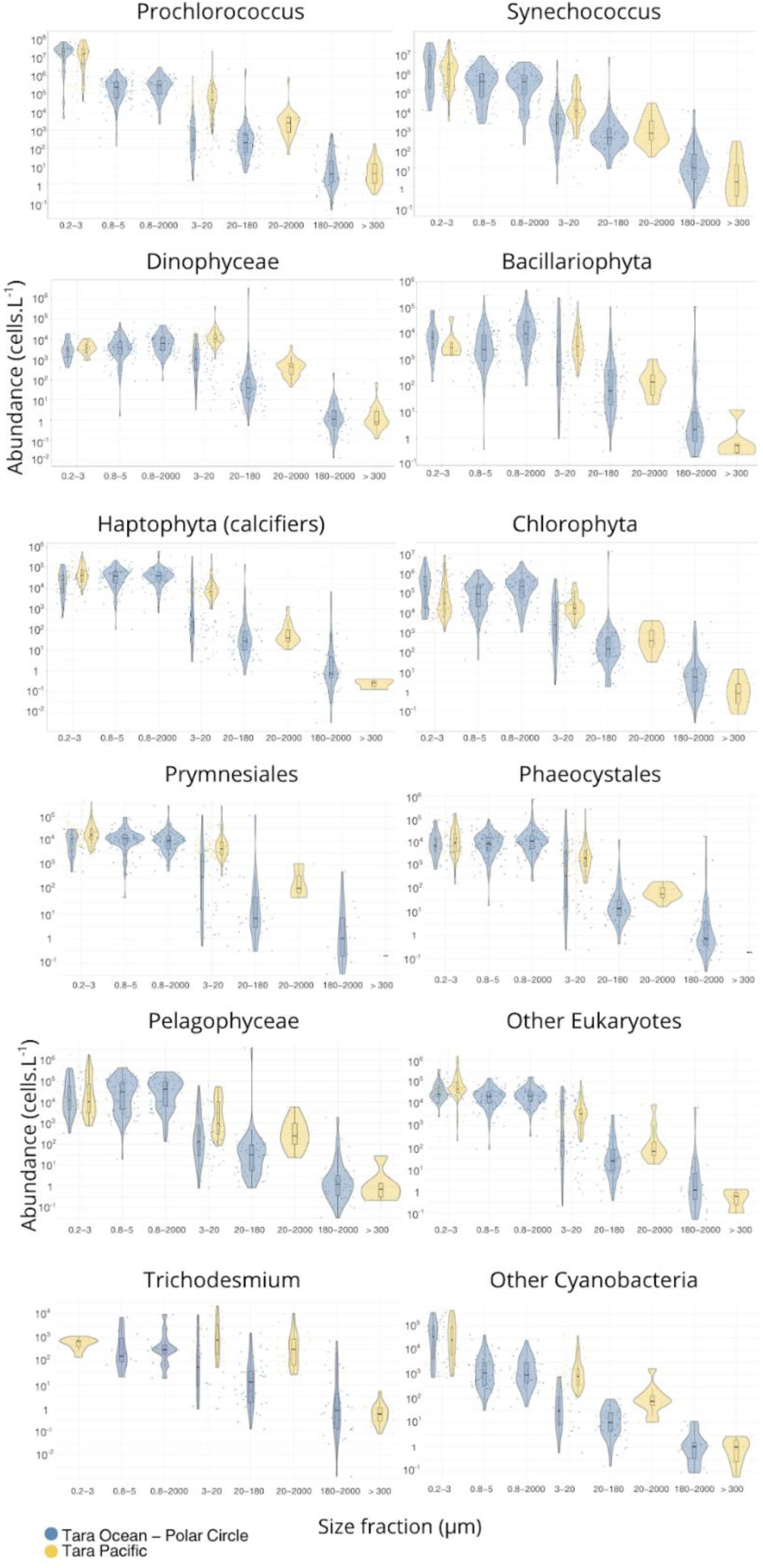
Distribution of cell abundances of photosynthetic lineages defined by the *psbO* marker gene across size fractions. Each point represents a surface sample collected during the Tara Oceans, Polar Circle (blue) and Tara Pacific (yellow) expeditions. Boxplots and violin plots show the distributions of cell abundance values (cells.L⁻¹) for each size fraction, along with the median values and interquartile ranges. Abundance values (y-axis) are displayed on a logarithmic scale.

**Fig. S8.**
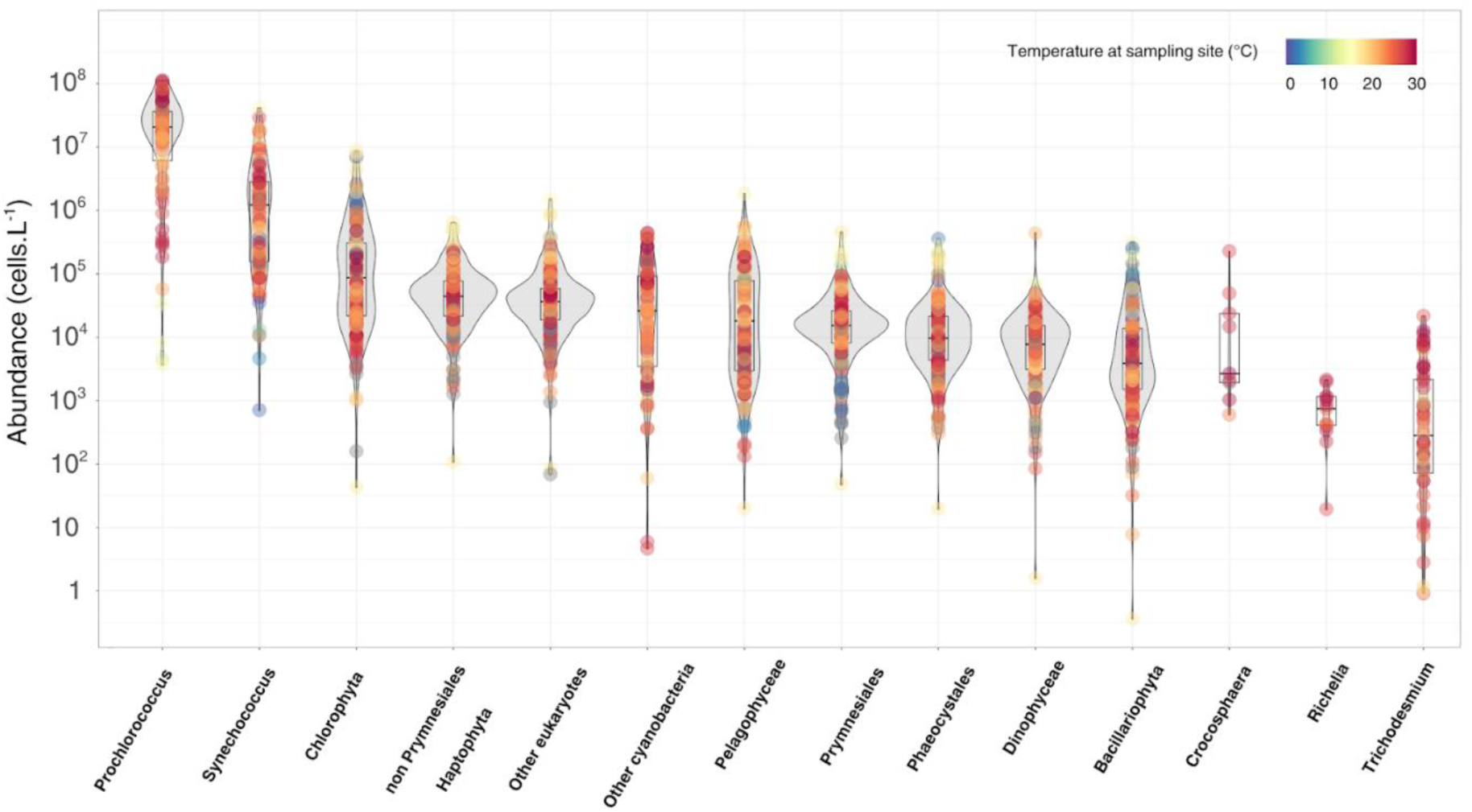
Estimates of total cell abundance, across the 0.2 to 20 μm size range, for eukaryotic and prokaryotic phytoplankton lineages defined by the *psbO* marker gene across *Tara* Oceans, Polar Circle and *Tara* Pacific surface stations. For each station, color indicates the sea surface temperature at the time of sampling. Significant correlations (padj < 0.05) between cell abundance and water temperature are indicated by Pearson correlation coefficients. Boxplots show the median and interquartile range of cell abundance values for each lineage. Lineages are ordered by decreasing median abundance.

**Fig. S9.**
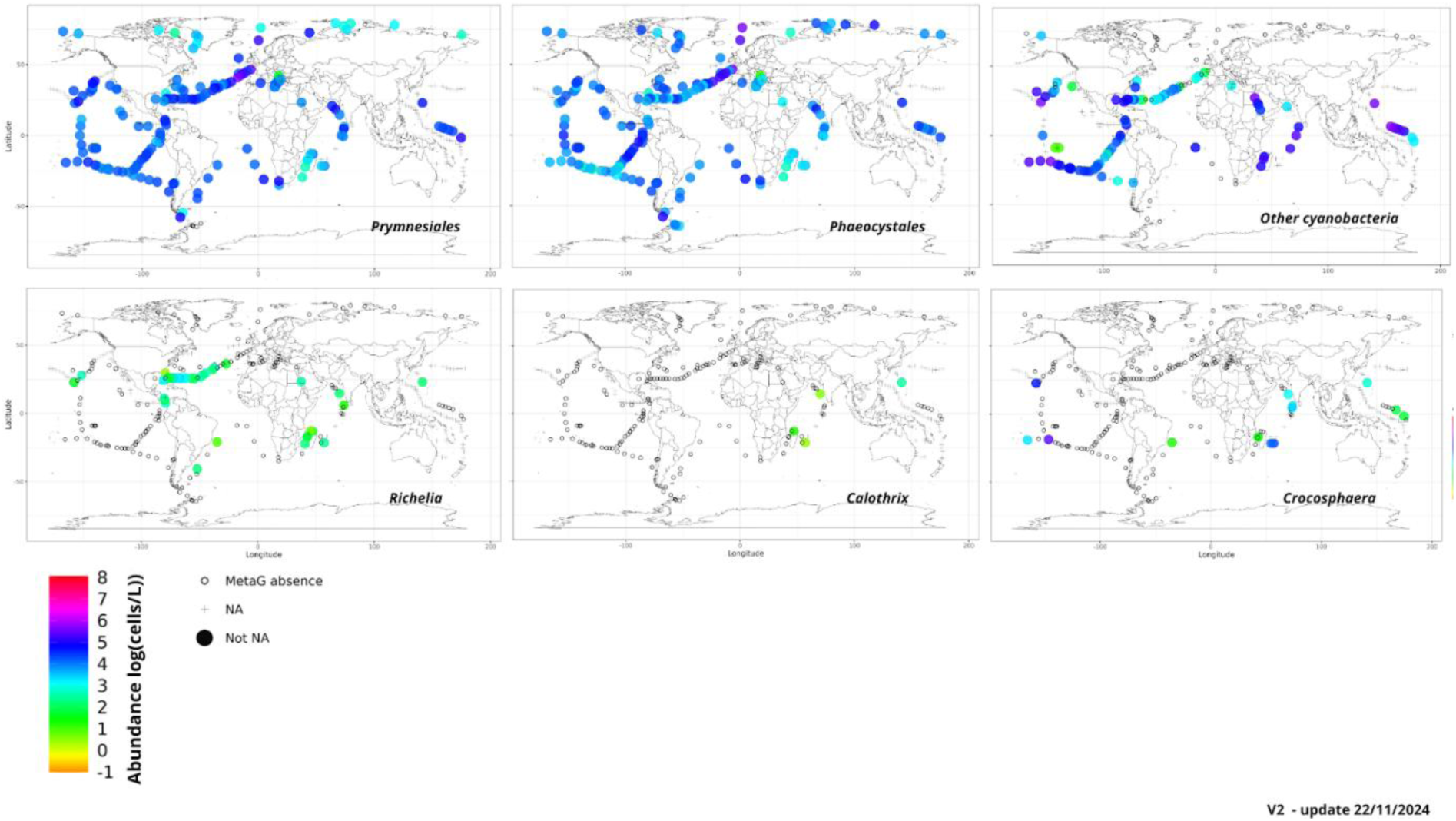
Global distribution of estimated total cell concentrations for phytoplankton lineages defined by the *psbO* marker gene across surface stations of the *Tara* Oceans, Polar Circle and *Tara* Pacific expeditions. Cell abundance values (log10(cells.L^-1^)) represent cumulative total cell concentrations across multiple organism size fractions ranging from 0.2 to 2,000 µm. Crosses (+) indicate stations with missing data for the corresponding lineage. Black circles (○) indicate stations where the lineage was not detected in the metagenomes of the available size fractions.

**Fig. S10.**
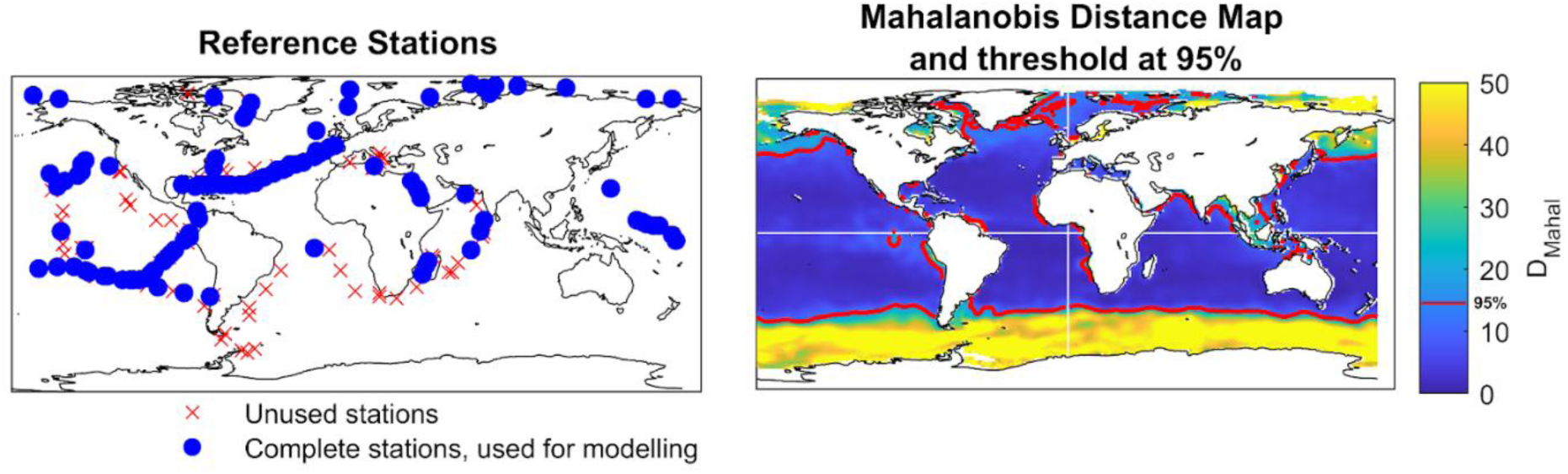
Training dataset and extrapolation domain. Geographic distribution of stations used in the training dataset (left) in blue, and unused stations marked by ×. Mahalanobis distance map (right) showing the environmental similarity of global ocean grid cells to the training dataset, with a 95% threshold overlay (red line). Regions beyond the threshold represent extrapolations into environmental conditions not well covered by the training dataset.

**Fig. S11.**
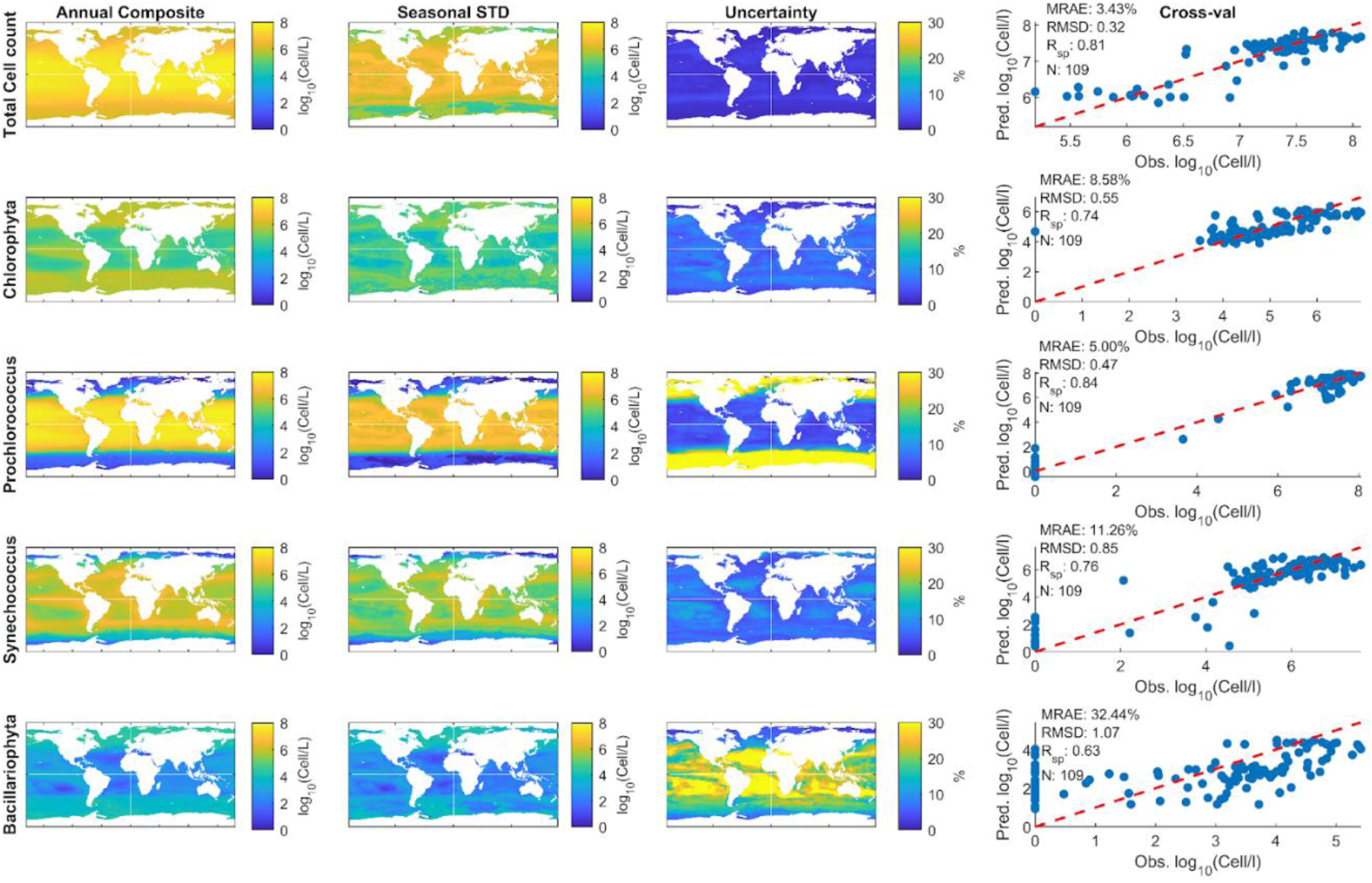
Global maps of phytoplankton cell abundance and model performance. Annual composite distributions, seasonal variability (standard deviation), model prediction uncertainty, and cross-validation of predicted versus observed log₁₀(cell L⁻¹) abundances are shown for total phytoplankton cell count, Chlorophyta, Synechococcus, Prochlorococcus, Bacillariophyta (diatoms), and Haptophyta. Cross-validation scatterplots show model predictive skill (blue points), with red dashed lines indicating 1:1 agreement for each group. Performance metrics include mean relative absolute error (MRAE), root mean square deviation (RMSD), and Spearman’s rank correlation (Rᵣₛₚ).

**Fig. S12.**
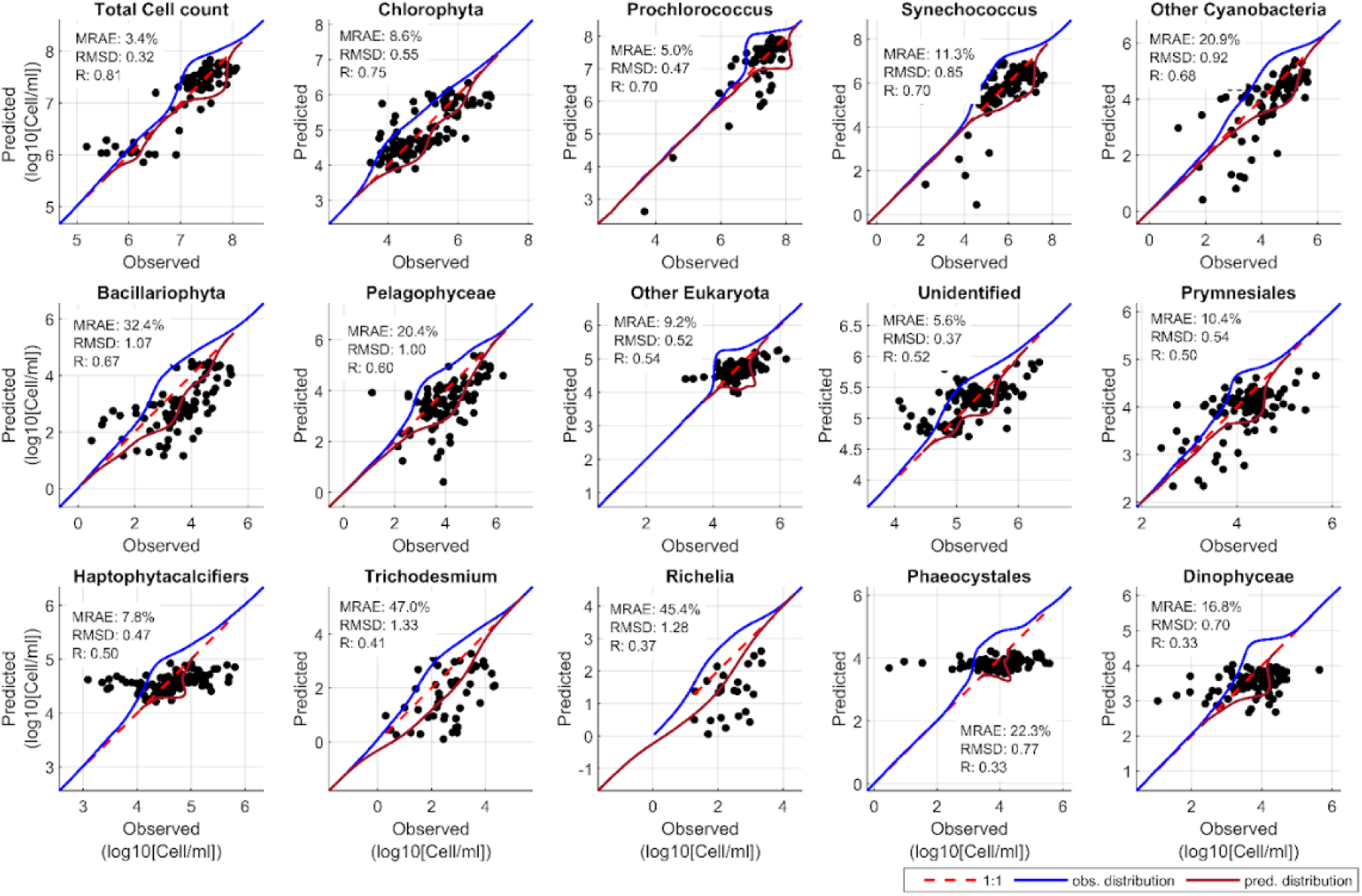
Cross-validation results. Observed versus predicted log₁₀ cell concentrations for total phytoplankton and 14 taxonomic groups. Each scatterplot shows model performance metrics (MRAE, RMSD, R). The 1:1 line (dashed red) and kernel density contours (blue = observed, red = predicted) illustrate the agreement and distribution overlap between modeled and observed values.

**Fig. S13.**
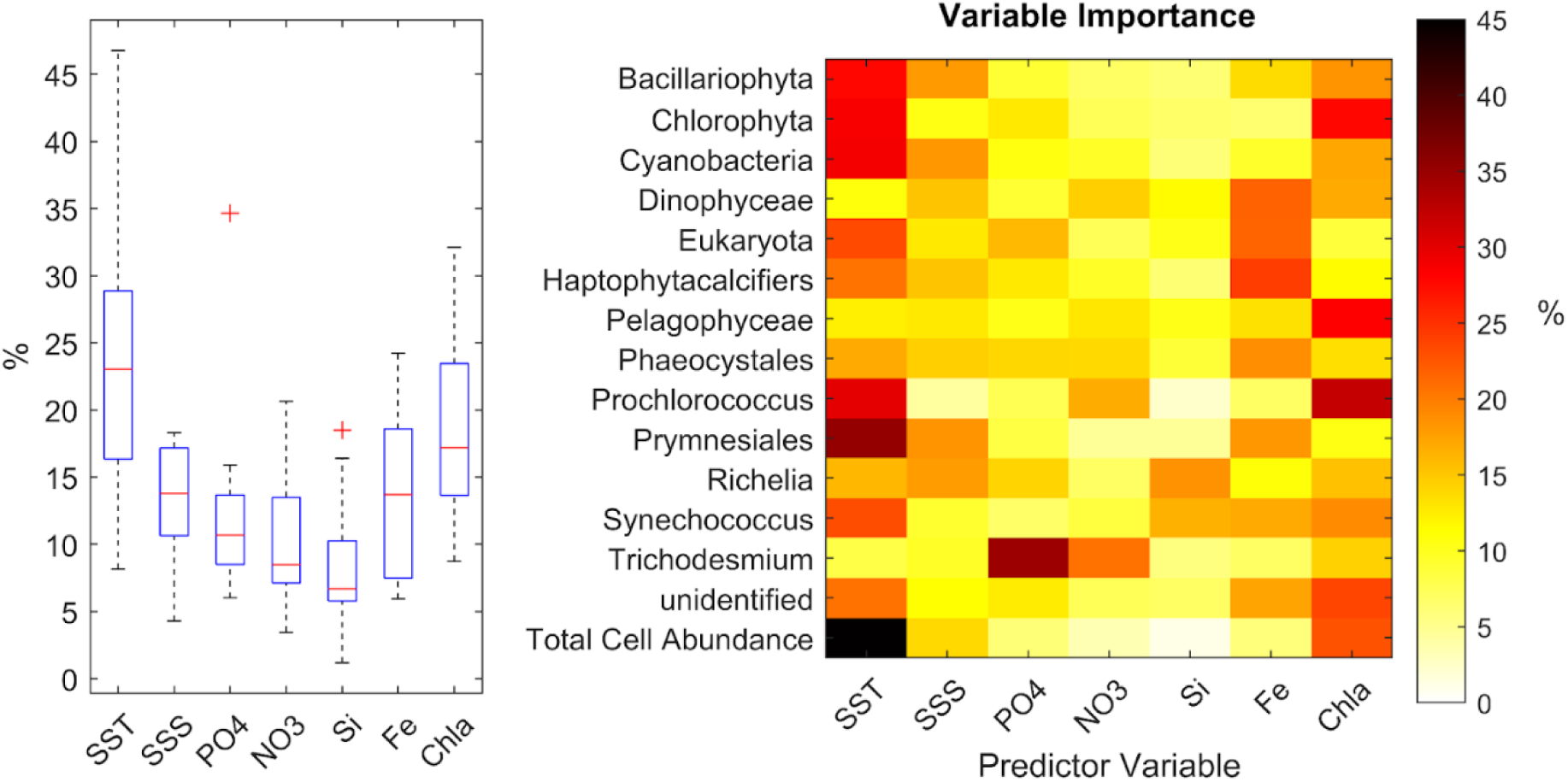
Relative importance of predictor variables used to model total cell abundance and 14 phytoplankton groups. Left: distribution of predictor importance across all groups, expressed as percentage contribution. Right: heatmap of variable importance by group, showing both consistent and group-specific patterns in predictor influence. SST = sea surface temperature, SSS = sea surface salinity, PO₄ = phosphate, NO₃ = nitrate, Si = silicate, Fe = iron, Chl-a = chlorophyll-a.

**Fig. S14.**
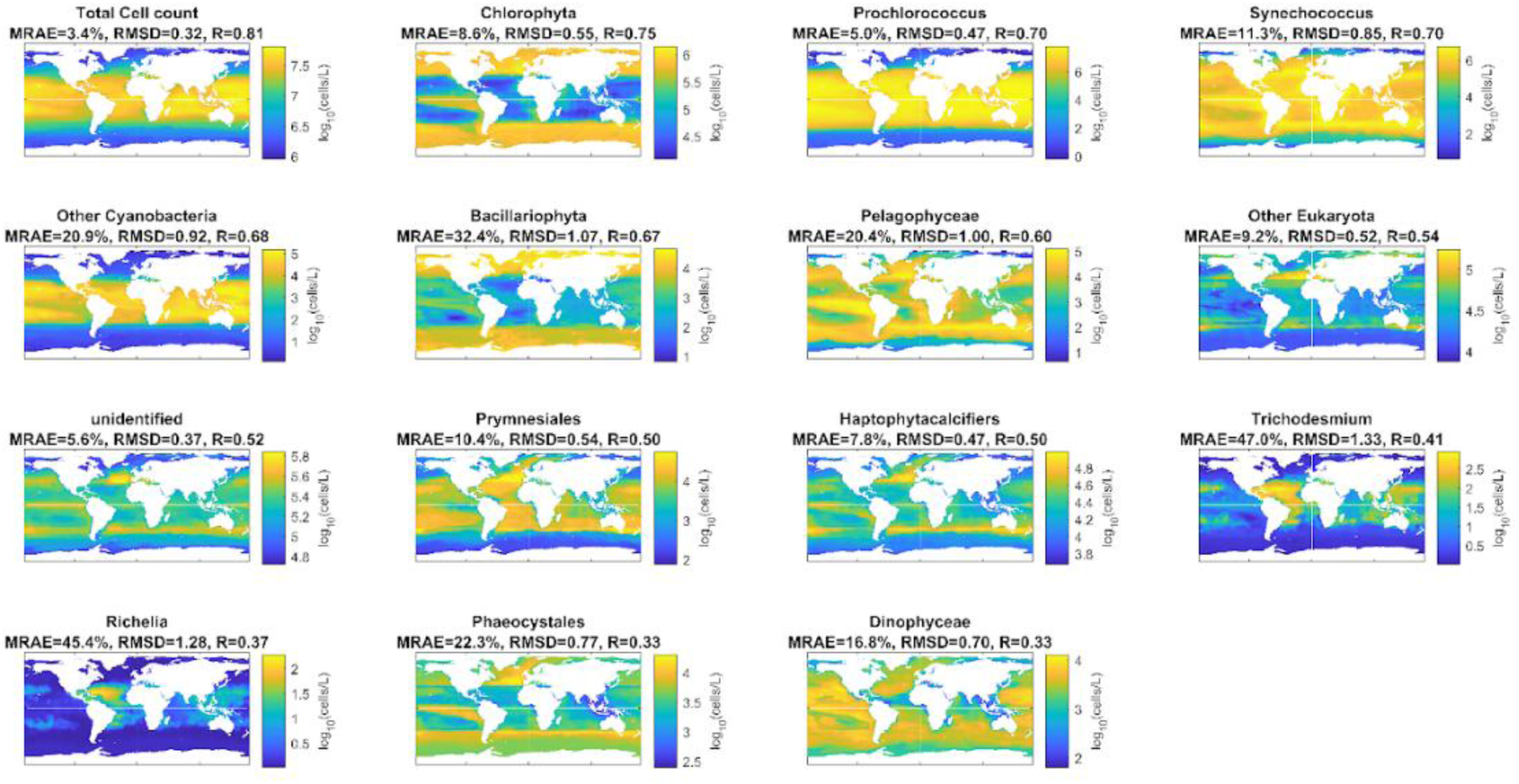
Mapping the annual composite distribution of each phytoplankton group. Modeled global distributions of total phytoplankton cell counts and 14 taxonomic groups expressed as log₁₀ (cells.L⁻¹). Results represent annual mean predictions across the global ocean. Model performance metrics are shown for each group (MRAE = mean relative absolute error; RMSD = root mean square deviation; R = spearman correlation).

**Fig. S15.**
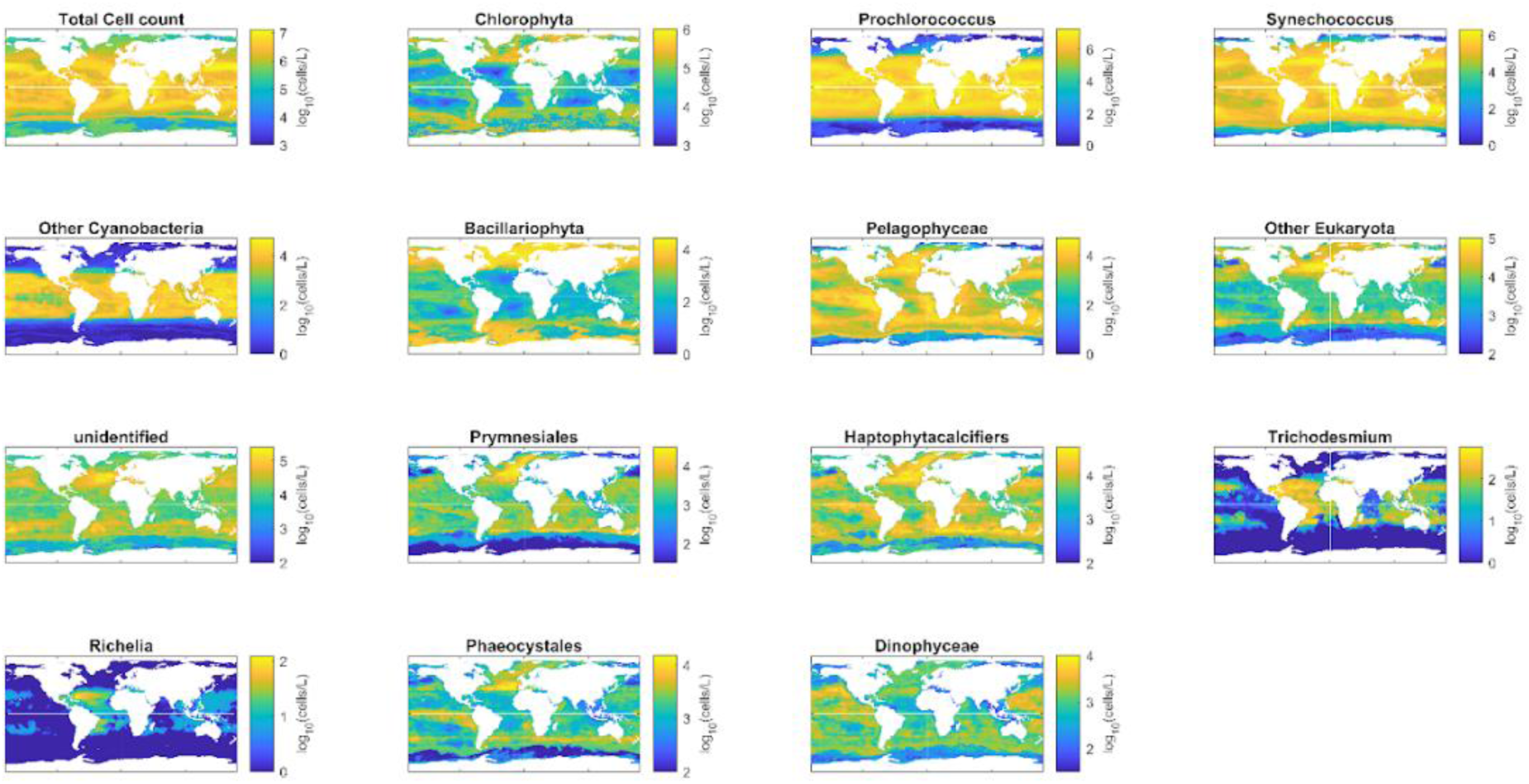
Global modeled seasonal variability (standard deviation). Modeled seasonal standard deviation of total phytoplankton cell counts and 14 taxonomic groups expressed as log₁₀ (cells.L⁻¹). The maps highlight regions of elevated temporal variability in phytoplankton abundance across the global ocean.

**Fig. S16.**
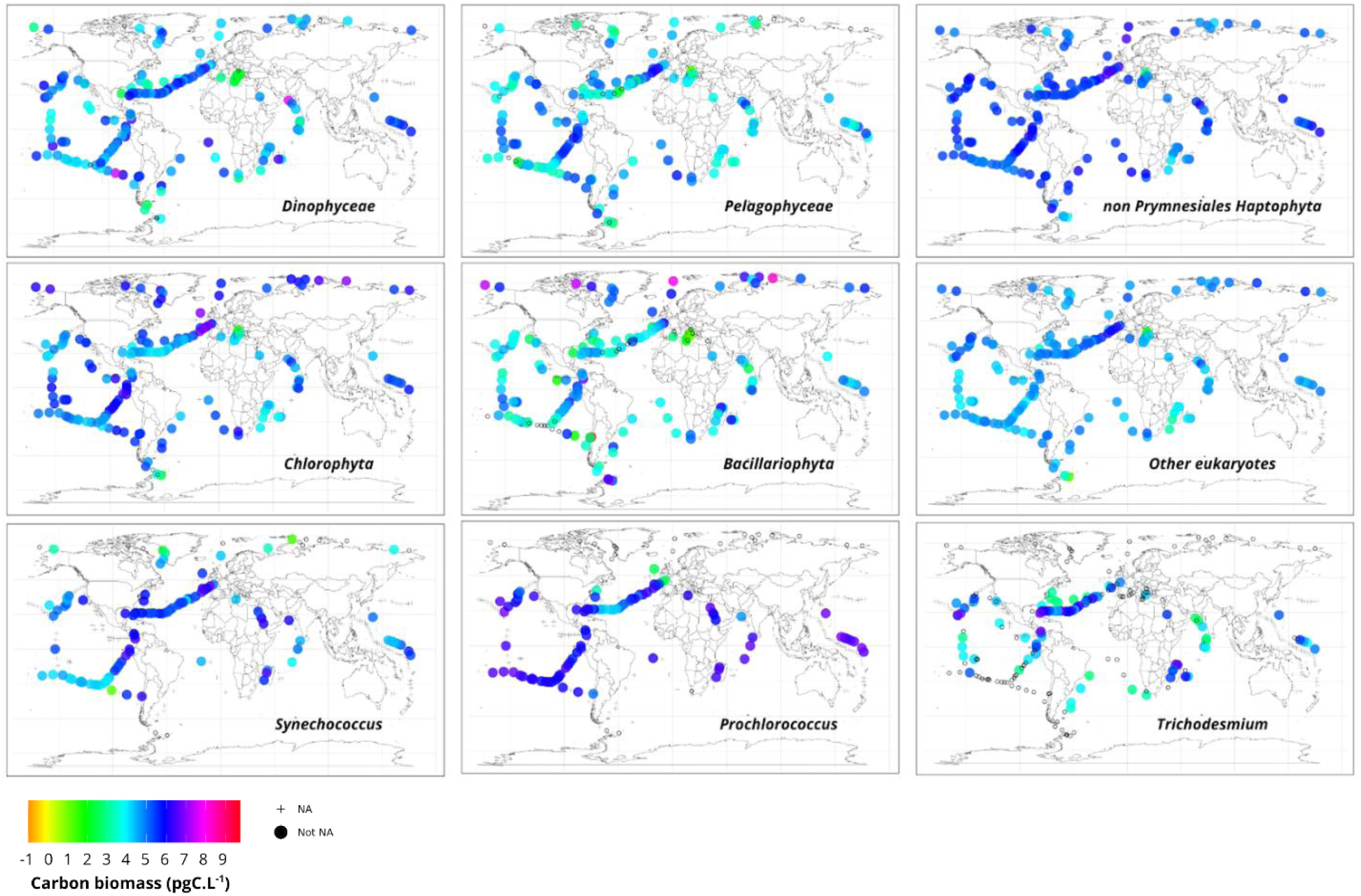
Global distribution of estimated total carbon biomass for phytoplankton lineages defined by the *psbO* marker gene across surface stations from the *Tara* Oceans, Polar Circle and *Tara* Pacific expeditions. Values (log10(pgC.L^-1^)) represent cumulative total carbon biomass across the 0.2 to 2,000 µm size range. Crosses (+) indicate stations with missing data for the corresponding lineage. Black circles (○) indicate stations where the lineage was not detected in the metagenomes of the available size fractions.

**Fig. S17.**
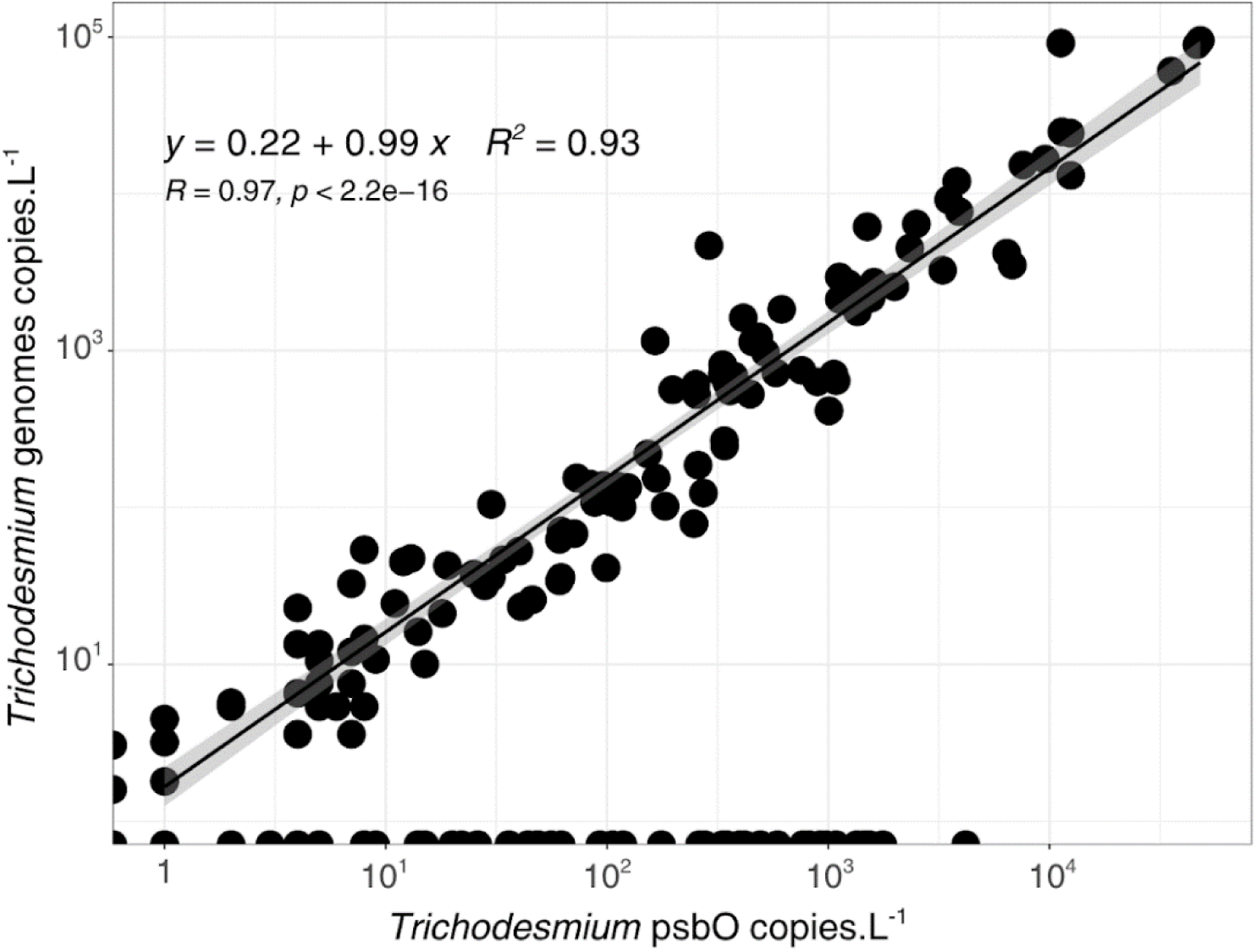
Comparison of *Trichodesmium* genome equivalents per liter estimated from environmental genome quantification versus *psbO* marker gene quantification. Each point represents a sample from one of the size fractions collected during the Tara Oceans expedition.

**Fig. S18.**
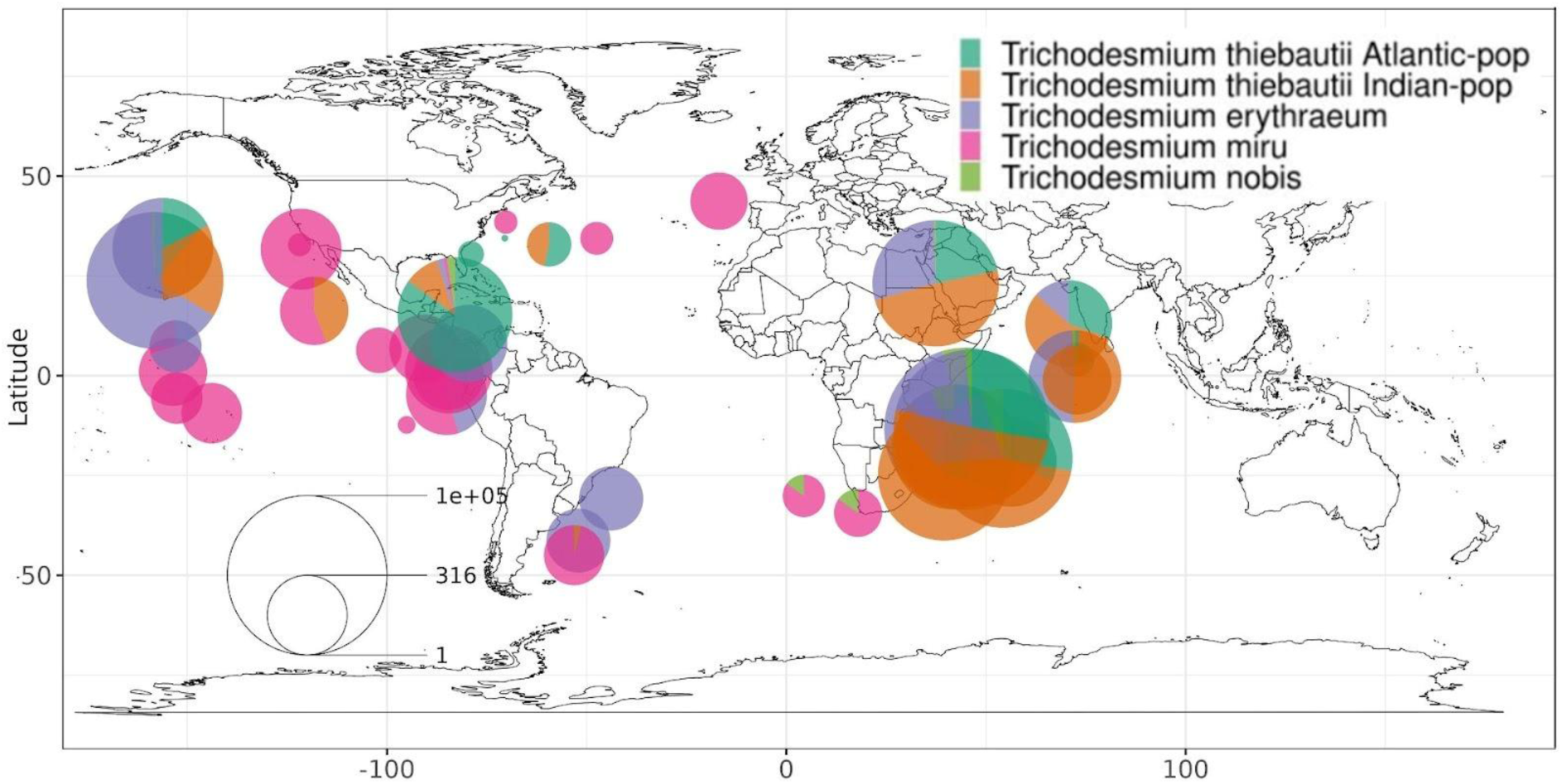
Abundance (log_10_(genome copies.L^-1^)) of the 5 MAGs of the genus Trichodesmium across Tara Oceans − Polar Circle surface stations. The cumulative abundance of each MAG is calculated across all size fractions from 0.8 to 2000 μm. The size of the pie charts represents the total abundance (log_10_(genome copies.L^−1^)) of all Trichodesmium MAGs at each sampling site.

**Table S1.** Network topology of genome-based co-occurrence graphs by size fraction under relative versus absolute abundance inputs. Mean weight: mean of weights assigned to edges, capturing the overall strength or “stability” of the associations / predicted interactions. Mean degree: mean number of connections or edges a given node has to other nodes, it is closely related to the density of a network. Mean strength (or weighted node degree): mean sum of edge weights of the adjacent edges for each node. Edge density: ratio between the number of edges and the number of possible edges. Transitivity (or clustering coefficient): measure of the degree to which nodes in a graph tend to cluster together (community clustering). Natural connectivity: measure of the redundancy of alternative paths in a network based on evaluating the weighted number of closed walks, it quantifies the robustness of a network.

| Size Fraction (μm) | Quantification Type | Mean Degree | Edge Density | Mean Strength | Natural Connectivity |
| --- | --- | --- | --- | --- | --- |
| 0.8 to 5/2000 | Relative Abundance | 2.59 | 0.01 | 1.29 | 0.01 |
|  | Absolute Abundance | 7.13 | 0.00 | 4.07 | 0.02 |
| 20 to 180 | Relative Abundance | 4.43 | 0.01 | 2.25 | 0.02 |
|  | Absolute Abundance | 7.13 | 0.00 | 4.07 | 0.02 |
| 180 to 2000 | Relative Abundance | 3.56 | 0.11 | 1.83 | 0.09 |
|  | Absolute Abundance | 4.32 | 0.01 | 2.87 | 0.03 |

## References

1. P. G. Falkowski, R. T. Barber, V. Smetacek, Biogeochemical Controls and Feedbacks on Ocean Primary Production. Science 281, 200–206 (1998).

2. J. F. Tjiputra, D. Couespel, R. Sanders, Marine ecosystem role in setting up preindustrial and future climate. Nat Commun 16, 2206 (2025).

3. A. R. Longhurst, Ecological Geography of the Sea (Academic Press, Amsterdam ; Boston, MA, 2nd ed., 2007).

4. E. T. Buitenhuis, W. K. W. Li, D. Vaulot, M. W. Lomas, M. R. Landry, F. Partensky, D. M. Karl, O. Ulloa, L. Campbell, S. Jacquet, F. Lantoine, F. Chavez, D. Macias, M. Gosselin, G. B. McManus, Picophytoplankton biomass distribution in the global ocean. Earth System Science Data 4, 37–46 (2012).

5. M. Winder, J. E. Cloern, The annual cycles of phytoplankton biomass. Phil. Trans. R. Soc. B 365, 3215– 3226 (2010).

6. L. A. Arteaga, E. Boss, M. J. Behrenfeld, T. K. Westberry, J. L. Sarmiento, Seasonal modulation of phytoplankton biomass in the Southern Ocean. Nat Commun 11, 5364 (2020).

7. R. El Hourany, J. Pierella Karlusich, L. Zinger, H. Loisel, M. Levy, C. Bowler, Linking satellites to genes with machine learning to estimate phytoplankton community structure from space. Ocean Sci. 20, 217–239 (2024).

8. F. M. Ibarbalz, N. Henry, F. Mahé, M. Ardyna, A. Zingone, E. Scalco, C. Lovejoy, F. Lombard, O. Jaillon, D. Iudicone, S. Malviya, Tara Oceans Coordinators, M. B. Sullivan, S. Chaffron, E. Karsenti, M. Babin, E. Boss, P. Wincker, L. Zinger, C. De Vargas, C. Bowler, L. Karp-Boss, Pan-Arctic plankton community structure and its global connectivity. Elementa: Science of the Anthropocene 11, 00060 (2023).

9. J. J. Pierella Karlusich, E. Pelletier, L. Zinger, F. Lombard, A. Zingone, S. Colin, J. M. Gasol, R. G. Dorrell, N. Henry, E. Scalco, S. G. Acinas, P. Wincker, C. De Vargas, C. Bowler, A robust approach to estimate relative phytoplankton cell abundances from metagenomes. Molecular Ecology Resources 23, 16–40 (2023).

10. Y. M. Bar-On, R. Phillips, R. Milo, The biomass distribution on Earth. Proc. Natl. Acad. Sci. U.S.A. 115, 6506–6511 (2018).

11. Y. Huang, D. Nicholson, B. Huang, N. Cassar, Global Estimates of Marine Gross Primary Production Based on Machine Learning Upscaling of Field Observations. Global Biogeochemical Cycles 35, e2020GB006718 (2021).

12. L. Paoli, H.-J. Ruscheweyh, C. C. Forneris, F. Hubrich, S. Kautsar, A. Bhushan, A. Lotti, Q. Clayssen, G. Salazar, A. Milanese, C. I. Carlström, C. Papadopoulou, D. Gehrig, M. Karasikov, H. Mustafa, M. Larralde, L. M. Carroll, P. Sánchez, A. A. Zayed, D. R. Cronin, S. G. Acinas, P. Bork, C. Bowler, T. O. Delmont, J. M. Gasol, A. D. Gossert, A. Kahles, M. B. Sullivan, P. Wincker, G. Zeller, S. L. Robinson, J. Piel, S. Sunagawa, Biosynthetic potential of the global ocean microbiome. Nature 607, 111–118 (2022).

13. M. A. Moran, E. B. Kujawinski, W. F. Schroer, S. A. Amin, N. R. Bates, E. M. Bertrand, R. Braakman, C. T. Brown, M. W. Covert, S. C. Doney, S. T. Dyhrman, A. S. Edison, A. M. Eren, N. M. Levine, L. Li, C. Ross, M. A. Saito, A. E. Santoro, D. Segrè, A. Shade, M. B. Sullivan, A. Vardi, Microbial metabolites in the marine carbon cycle. Nat Microbiol 7, 508–523 (2022).

14. D. Lovell, W. Muller, J. Taylor, A. Zwart, C. Helliwell, Caution! Compositions! Can constraints on omics data lead analyses astray? (2010).

15. G. B. Gloor, J. M. Macklaim, V. Pawlowsky-Glahn, J. J. Egozcue, Microbiome Datasets Are Compositional: And This Is Not Optional. Front. Microbiol. 8, 2224 (2017).

16. M. Greenacre, Compositional Data Analysis. Annu. Rev. Stat. Appl. 8, 271–299 (2021).

17. T. P. Quinn, I. Erb, G. Gloor, C. Notredame, M. F. Richardson, T. M. Crowley, A field guide for the compositional analysis of any-omics data. GigaScience 8, giz107 (2019).

18. J. Aitchison, The Statistical Analysis of Compositional Data. (1986).

19. D. Lovell, V. Pawlowsky-Glahn, J. J. Egozcue, S. Marguerat, J. Bähler, Proportionality: A Valid Alternative to Correlation for Relative Data. PLoS Comput Biol 11, e1004075 (2015).

20. T. O. Delmont, Discovery of nondiazotrophic *Trichodesmium* species abundant and widespread in the open ocean. Proc. Natl. Acad. Sci. U.S.A. 118, e2112355118 (2021).

21. Z. Li, W. J. Riley, G. L. Marschmann, U. Karaoz, I. A. Shirley, Q. Wu, N. J. Bouskill, K.-Y. Chang, P. M. Crill, R. F. Grant, E. King, S. R. Saleska, M. B. Sullivan, J. Tang, R. K. Varner, B. J. Woodcroft, K. C. Wrighton, the EMERGE Biology Integration Institute Coordinators, E. L. Brodie, A framework for integrating genomics, microbial traits, and ecosystem biogeochemistry. Nat Commun 16 (2025).

22. T. Mock, S. J. Daines, R. Geider, S. Collins, M. Metodiev, A. J. Millar, V. Moulton, T. M. Lenton, Bridging the gap between omics and earth system science to better understand how environmental change impacts marine microbes. Global Change Biology 22, 61–75 (2016).

23. C. L. Quéré, S. P. Harrison, I. Colin Prentice, E. T. Buitenhuis, O. Aumont, L. Bopp, H. Claustre, L. Cotrim Da Cunha, R. Geider, X. Giraud, C. Klaas, K. E. Kohfeld, L. Legendre, M. Manizza, T. Platt, R. B. Rivkin, S. Sathyendranath, J. Uitz, A. J. Watson, D. Wolf-Gladrow, Ecosystem dynamics based on plankton functional types for global ocean biogeochemistry models. Global Change Biology 11, 2016– 2040 (2005).

24. O. Aumont, C. Ethé, A. Tagliabue, L. Bopp, M. Gehlen, PISCES-v2: an ocean biogeochemical model for carbon and ecosystem studies. Geosci. Model Dev. 8, 2465–2513 (2015).

25. A. Régimbeau, O. Aumont, C. Bowler, L. Guidi, G. A. Jackson, E. Karsenti, L. Memery, A. Tagliabue, D. Eveillard, Unveiling the link between phytoplankton molecular physiology and biogeochemical cycling via genome-scale modeling. Sci. Adv. 11 (2025).

26. N. Giordano, M. Gaudin, C. Trottier, E. Delage, C. Nef, C. Bowler, S. Chaffron, Genome-scale community modelling reveals conserved metabolic cross-feedings in epipelagic bacterioplankton communities. Nature Communications 15, 2721 (2024).

27. D. Vandeputte, G. Kathagen, K. D’hoe, S. Vieira-Silva, M. Valles-Colomer, J. Sabino, J. Wang, R. Y. Tito, L. De Commer, Y. Darzi, S. Vermeire, G. Falony, J. Raes, Quantitative microbiome profiling links gut community variation to microbial load. Nature 551, 507–511 (2017).

28. Y. Lin, S. Gifford, H. Ducklow, O. Schofield, N. Cassar, Towards Quantitative Microbiome Community Profiling Using Internal Standards. Appl Environ Microbiol 85, e02634–18 (2019).

29. E. J. Contijoch, G. J. Britton, C. Yang, I. Mogno, Z. Li, R. Ng, S. R. Llewellyn, S. Hira, C. Johnson, K. M. Rabinowitz, R. Barkan, I. Dotan, R. P. Hirten, S.-C. Fu, Y. Luo, N. Yang, T. Luong, P. R. Labrias, S. Lira, I. Peter, A. Grinspan, J. C. Clemente, R. Kosoy, S. Kim-Schulze, X. Qin, A. Castillo, A. Hurley, A. Atreja, J. Rogers, F. Fasihuddin, M. Saliaj, A. Nolan, P. Reyes-Mercedes, C. Rodriguez, S. Aly, K. Santa-Cruz, L. Peters, M. Suárez-Fariñas, R. Huang, K. Hao, J. Zhu, B. Zhang, B. Losic, H. Irizar, W.M. Song, A. Di Narzo, W. Wang, B. L. Cohen, C. DiMaio, D. Greenwald, S. Itzkowitz, A. Lucas, J. Marion, E. Maser, R. Ungaro, S. Naymagon, J. Novak, B. Shah, T. Ullman, P. Rubin, J. George, P. Legnani, S. E. Telesco, J. R. Friedman, C. Brodmerkel, S. Plevy, J. H. Cho, J.-F. Colombel, E. E. Schadt, Argmann, M. Dubinsky, A. Kasarskis, B. Sands, J. J. Faith, Gut microbiota density influences host physiology and is shaped by host and microbial factors. eLife 8, e40553 (2019).

30. J. T. Barlow, S. R. Bogatyrev, R. F. Ismagilov, A quantitative sequencing framework for absolute abundance measurements of mucosal and lumenal microbial communities. Nat Commun 11, 2590 (2020).

31. J. G. Harrison, W. John Calder, B. Shuman, C. Alex Buerkle, The quest for absolute abundance: The use of internal standards for DNA-based community ecology. Molecular Ecology Resources 21, 30–43 (2021).

32. S. M. Gifford, S. Sharma, J. M. Rinta-Kanto, M. A. Moran, Quantitative analysis of a deeply sequenced marine microbial metatranscriptome. The ISME Journal 5, 461–472 (2011).

33. B. M. Satinsky, S. M. Gifford, B. C. Crump, M. A. Moran, “Use of Internal Standards for Quantitative Metatranscriptome and Metagenome Analysis” in Methods in Enzymology (Elsevier, 2013; https://linkinghub.elsevier.com/retrieve/pii/B9780124078635000125)vol. 531, pp. 237–250.

34. Q. Bei, N. L. R. Williams, L. E. Furtado, D. D. Blasi, J. Williams, V. Brotas, G. Tarran, A. P. Rees, C. Bowler, J. A. Fuhrman, Quantitative metagenomics for marine prokaryotes and photosynthetic eukaryotes. ISME Communications 5, ycaf131 (2025).

35. C. Jian, P. Luukkonen, H. Yki-Järvinen, A. Salonen, K. Korpela, Quantitative PCR provides a simple and accessible method for quantitative microbiota profiling. PLoS ONE 15, e0227285 (2020).

36. J. J. Pierella Karlusich, E. Pelletier, F. Lombard, M. Carsique, E. Dvorak, S. Colin, M. Picheral, F. M. Cornejo-Castillo, S. G. Acinas, R. Pepperkok, E. Karsenti, C. De Vargas, P. Wincker, C. Bowler, R. A. Foster, Global distribution patterns of marine nitrogen-fixers by imaging and molecular methods. Nat Commun 12, 4160 (2021).

37. M. Royo-Llonch, P. Sánchez, C. Ruiz-González, G. Salazar, C. Pedrós-Alió, M. Sebastián, K. Labadie, L. Paoli, F. M. Ibarbalz, L. Zinger, B. Churcheward, Tara Oceans Coordinators, M. Babin, P. Bork, E. Boss, G. Cochrane, C. De Vargas, G. Gorsky, N. Grimsley, L. Guidi, P. Hingamp, D. Iudicone, O. Jaillon, S. Kandels, F. Not, H. Ogata, S. Pesant, N. Poulton, J. Raes, C. Sardet, S. Speich, L. Setmmann, M. B. Sullivan, S. Chaffron, D. Eveillard, E. Karsenti, S. Sunagawa, P. Wincker, L. Karp-Boss, C. Bowler, S. G. Acinas, Compendium of 530 metagenome-assembled bacterial and archaeal genomes from the polar Arctic Ocean. Nat Microbiol 6, 1561–1574 (2021).

38. T. O. Delmont, M. Gaia, D. D. Hinsinger, P. Frémont, C. Vanni, A. Fernandez-Guerra, A. M. Eren, A. Kourlaiev, L. d’Agata, Q. Clayssen, E. Villar, K. Labadie, C. Cruaud, J. Poulain, C. Da Silva, M. Wessner, B. Noel, J.-M. Aury, C. De Vargas, C. Bowler, E. Karsenti, E. Pelletier, P. Wincker, O. Jaillon, S. Sunagawa, S. G. Acinas, P. Bork, E. Karsenti, C. Bowler, C. Sardet, L. Stemmann, C. De Vargas, P. Wincker, M. Lescot, M. Babin, G. Gorsky, N. Grimsley, L. Guidi, P. Hingamp, O. Jaillon, S. Kandels, Iudicone, H. Ogata, S. Pesant, M. B. Sullivan, F. Not, K.-B. Lee, E. Boss, G. Cochrane, M. Follows, N. Poulton, J. Raes, M. Sieracki, S. Speich, Functional repertoire convergence of distantly related eukaryotic plankton lineages abundant in the sunlit ocean. Cell Genomics 2, 100123 (2022).

39. E. Karsenti, S. G. Acinas, P. Bork, C. Bowler, C. De Vargas, J. Raes, M. Sullivan, D. Arendt, F. Benzoni, J.-M. Claverie, M. Follows, G. Gorsky, P. Hingamp, D. Iudicone, O. Jaillon, S. Kandels-Lewis, U. Krzic, F. Not, H. Ogata, S. Pesant, E. G. Reynaud, C. Sardet, M. E. Sieracki, S. Speich, D. Velayoudon, J. Weissenbach, P. Wincker, the Tara Oceans Consortium, A Holistic Approach to Marine Eco-Systems Biology. PLoS Biol 9, e1001177 (2011).

40. S. Pesant, F. Not, M. Picheral, S. Kandels-Lewis, N. Le Bescot, G. Gorsky, D. Iudicone, E. Karsenti, S. Speich, R. Troublé, C. Dimier, S. Searson, Tara Oceans Consortium Coordinators, S. G. Acinas, P. Bork, Boss, C. Bowler, C. De Vargas, M. Follows, G. Gorsky, N. Grimsley, P. Hingamp, D. Iudicone, O. Jaillon, S. Kandels-Lewis, L. Karp-Boss, E. Karsenti, U. Krzic, F. Not, H. Ogata, S. Pesant, J. Raes, E. G. Reynaud, C. Sardet, M. Sieracki, S. Speich, L. Stemmann, M. B. Sullivan, S. Sunagawa, D. Velayoudon, J. Weissenbach, P. Wincker, Open science resources for the discovery and analysis of Tara Oceans data. Sci Data 2, 150023 (2015).

41. S. Planes, D. Allemand, S. Agostini, B. Banaigs, E. Boissin, E. Boss, G. Bourdin, C. Bowler, E. Douville, J. M. Flores, D. Forcioli, P. Furla, P. E. Galand, J.-F. Ghiglione, E. Gilson, F. Lombard, C. Moulin, S. Pesant, J. Poulain, S. Reynaud, S. Romac, M. B. Sullivan, S. Sunagawa, O. P. Thomas, R. Troublé, C. De Vargas, R. Vega Thurber, C. R. Voolstra, P. Wincker, D. Zoccola, the Tara Pacific Consortium, The Tara Pacific expedition—A pan-ecosystemic approach of the “-omics” complexity of coral reef holobionts across the Pacific Ocean. PLoS Biol 17, e3000483 (2019).

42. G. Gorsky, G. Bourdin, F. Lombard, M. L. Pedrotti, S. Audrain, N. Bin, E. Boss, C. Bowler, N. Cassar, L. Caudan, G. Chabot, N. R. Cohen, D. Cron, C. De Vargas, J. R. Dolan, E. Douville, A. Elineau, J. M. Flores, J. F. Ghiglione, N. Haëntjens, M. Hertau, S. G. John, R. L. Kelly, I. Koren, Y. Lin, D. Marie, C. Moulin, Y. Moucherie, S. Pesant, M. Picheral, J. Poulain, M. Pujo-Pay, G. Reverdin, S. Romac, M. B. Sullivan, M. Trainic, M. Tressol, R. Troublé, A. Vardi, C. R. Voolstra, P. Wincker, S. Agostini, B. Banaigs, E. Boissin, D. Forcioli, P. Furla, P. E. Galand, E. Gilson, S. Reynaud, S. Sunagawa, O. P. Thomas, R. L. V. Thurber, D. Zoccola, S. Planes, D. Allemand, E. Karsenti, Expanding Tara Oceans Protocols for Underway, Ecosystemic Sampling of the Ocean-Atmosphere Interface During Tara Pacific Expedition (2016–2018). Front. Mar. Sci. 6, 750 (2019).

43. F. Lombard, G. Bourdin, S. Pesant, S. Agostini, A. Baudena, E. Boissin, N. Cassar, M. Clampitt, P. Conan, O. Da Silva, C. Dimier, E. Douville, A. Elineau, J. Fin, J. M. Flores, J.-F. Ghiglione, B. C. C. Hume, L. Jalabert, S. G. John, R. L. Kelly, I. Koren, Y. Lin, D. Marie, R. McMinds, Z. Mériguet, N. Metzl, D. A. Paz-García, M. L. Pedrotti, J. Poulain, M. Pujo-Pay, J. Ras, G. Reverdin, S. Romac, A. Rouan, E. Röttinger, A. Vardi, C. R. Voolstra, C. Moulin, G. Iwankow, B. Banaigs, C. Bowler, C. De Vargas, D. Forcioli, P. Furla, P. E. Galand, E. Gilson, S. Reynaud, S. Sunagawa, M. B. Sullivan, O. P. Thomas, R. Troublé, R. V. Thurber, P. Wincker, D. Zoccola, D. Allemand, S. Planes, E. Boss, G. Gorsky, Open science resources from the Tara Pacific expedition across coral reef and surface ocean ecosystems. Sci Data 10, 324 (2023).

44. A. Alberti, J. Poulain, S. Engelen, K. Labadie, S. Romac, I. Ferrera, G. Albini, J.-M. Aury, C. Belser, A. Bertrand, C. Cruaud, C. Da Silva, C. Dossat, F. Gavory, S. Gas, J. Guy, M. Haquelle, E. Jacoby, O. Jaillon, A. Lemainque, E. Pelletier, G. Samson, M. Wessner, Genoscope Technical Team, P. Bazire, O. Beluche, L. Bertrand, M. Besnard-Gonnet, I. Bordelais, M. Boutard, M. Dubois, C. Dumont, E. Ettedgui, P. Fernandez, E. Garcia, N. G. Aiach, T. Guerin, C. Hamon, E. Brun, S. Lebled, P. Lenoble, C. Louesse, E. Mahieu, B. Mairey, N. Martins, C. Megret, C. Milani, J. Muanga, C. Orvain, E. Payen, P. Perroud, E. Petit, D. Robert, M. Ronsin, B. Vacherie, S. G. Acinas, M. Royo-Llonch, F. M. Cornejo-Castillo, R. Logares, B. Fernández-Gómez, C. Bowler, G. Cochrane, C. Amid, P. T. Hoopen, C. De Vargas, N. Grimsley, E. Desgranges, S. Kandels-Lewis, H. Ogata, N. Poulton, M. E. Sieracki, R. Stepanauskas, M. B. Sullivan, J. R. Brum, M. B. Duhaime, B. T. Poulos, B. L. Hurwitz, Tara Oceans Consortium Coordinators, S. G. Acinas, P. Bork, E. Boss, C. Bowler, C. De Vargas, M. Follows, G. Gorsky, N. Grimsley, P. Hingamp, D. Iudicone, O. Jaillon, S. Kandels-Lewis, L. Karp-Boss, E. Karsenti, F. Not, H. Ogata, S. Pesant, J. Raes, C. Sardet, M. E. Sieracki, S. Speich, L. Stemmann, M. B. Sullivan, S. Sunagawa, P. Wincker, S. Pesant, E. Karsenti, P. Wincker, Viral to metazoan marine plankton nucleotide sequences from the Tara Oceans expedition. Sci Data 4, 170093 (2017).

45. C. Belser, J. Poulain, K. Labadie, F. Gavory, A. Alberti, J. Guy, Q. Carradec, C. Cruaud, C. Da Silva, S. Engelen, P. Mielle, A. Perdereau, G. Samson, S. Gas, Genoscope Technical Team, J. Batisse, O. Beluche, L. Bertrand, C. Bohers, I. Bordelais, E. Brun, M. Dubois, C. Dumont, E. H. Zineb, B. Estrada, E. Ettedgui, P. Fernandez, S. Garidi, T. Guérin, K. Gorrichon, C. Hamon, L. Kientzel, S. Lebled, C. Legrain, P. Lenoble, M. Lepretre, C. Louesse, G. Magdelenat, E. Mahieu, N. Martins, C. Milani, C. Orvain, S. Oztas, E. Payen, E. Petit, G. Rio, D. Robert, M. Ronsin, B. Vacherie, C. R. Voolstra, P. E. Galand, J. M. Flores, B. C. C. Hume, G. Perna, M. Ziegler, H.-J. Ruscheweyh, E. Boissin, S. Romac, G. Bourdin, G. Iwankow, C. Moulin, D. A. Paz García, S. Agostini, B. Banaigs, E. Boss, C. Bowler, C. De Vargas, E. Douville, D. Forcioli, P. Furla, E. Gilson, F. Lombard, S. Pesant, S. Reynaud, S. Sunagawa, O. P. Thomas, R. Troublé, R. V. Thurber, D. Zoccola, C. Scarpelli, E. K. Jacoby, P. H. Oliveira, J.-M. Aury, D. Allemand, S. Planes, P. Wincker, Integrative omics framework for characterization of coral reef ecosystems from the Tara Pacific expedition. Sci Data 10, 326 (2023).

46. Y. Ji, T. Huotari, T. Roslin, N. M. Schmidt, J. Wang, D. W. Yu, O. Ovaskainen, SPIKEPIPE: A metagenomic pipeline for the accurate quantification of eukaryotic species occurrences and intraspecific abundance change using DNA barcodes or mitogenomes. Molecular Ecology Resources 20, 256–267 (2020).

47. B. Li, X. Li, T. Yan, A Quantitative Metagenomic Sequencing Approach for High-Throughput Gene Quantification and Demonstration with Antibiotic Resistance Genes. Appl Environ Microbiol 87, e00871–21 (2021).

48. S. Nayfach, K. S. Pollard, Toward Accurate and Quantitative Comparative Metagenomics. Cell 166, 1103–1116 (2016).

49. S. Sathyendranath, J. Aiken, S. Alvain, R. Barlow, H. Bouman, A. Bracher, R. Brewin, A. Bricaud, C. W. Brown, A. M. Ciotti, L. A. Clementson, S. E. Craig, E. Devred, N. Hardman-Mountford, T. Hirata, C. Hu, T. S. Kostadinov, S. Lavender, H. Loisel, T. S. Moore, J. Morales, C. B. Mouw, A. Nair, D. Raitsos, C. Roesler, J. D. Shutler, H. M. Sosik, I. Soto, V. Stuart, A. Subramaniam, J. Uitz, Phytoplankton functional types from Space., S. Sathyendranath, V. Stuart, Eds., (Reports of the International Ocean-Colour Coordinating Group (IOCCG) ; 15) (2014)pp. 1–156.

50. Lombard, G. Lionel, M. C. Brandão, C. L. Pedro, C. Sébastien, D. J. Richard, E. Amanda, J. M. Gasol, G. P. Luc, H. Nicolas, F. M. Ibarbalz, J. Laëtitia, L. Michel, M. Séverinne, M. Zoé, P. Marc, J. J. Pierella Karlusich, R. Pepperkok, R. Jean-Baptiste, Z. Lucie, Tara Oceans Coordinators, S. Lars, S. G. Acinas, K.-B. Lee, B. Emmanuel, M. B. Sullivan, C. De Vargas, B. Chris, K. Eric, G. Gabriel, Ubiquity of inverted ’gelatinous’ ecosystem pyramids in the global ocean. [Preprint] (2024). 10.1101/2024.02.09.579612.

51. Z. Mériguet, G. Bourdin, N. Kristan, L. Jalabert, O. Bun, M. Picheral, L. Caray-Counil, J. Maury, M.-L. Pedrotti, A. Elineau, D. A. Paz-Garcia, L. Karp-Boss, G. Gorsky, F. Lombard, the Tara Pacific Consortium Coordinators team, Quantitative imaging datasets of surface micro-to mesoplankton communities and microplastic across the Pacific and North Atlantic oceans from the Tara Pacific expedition. Earth Syst. Sci. Data 17, 2761–2792 (2025).

52. D. Marie, S. Romac, Flow cytometry data of phytoplankton, bacteria and viruses obtained for samples collected during the Tara Pacific Expedition 2016-2018, PANGAEA (2022); 10.1594/PANGAEA.944490.

53. P. Hingamp, N. Grimsley, S. G. Acinas, C. Clerissi, L. Subirana, J. Poulain, I. Ferrera, H. Sarmento, E. Villar, G. Lima-Mendez, K. Faust, S. Sunagawa, J.-M. Claverie, H. Moreau, Y. Desdevises, P. Bork, J. Raes, C. De Vargas, E. Karsenti, S. Kandels-Lewis, O. Jaillon, F. Not, S. Pesant, P. Wincker, H. Ogata, Exploring nucleo-cytoplasmic large DNA viruses in Tara Oceans microbial metagenomes. The ISME Journal 7, 1678–1695 (2013).

54. M. Thyssen, G. Grégori, V. Créach, S. Lahbib, M. Dugenne, H. M. Aardema, L.-F. Artigas, B. Huang, A. Barani, L. Beaugeard, A. Bellaaj-Zouari, A. Beran, R. Casotti, Y. Del Amo, M. Denis, G. B. J. Dubelaar, S. Endres, L. Haraguchi, B. Karlson, C. Lambert, A. Louchart, D. Marie, G. Moncoiffé, D. Pecqueur, F. Ribalet, M. Rijkeboer, T. Silovic, R. Silva, S. Marro, H. M. Sosik, M. Sourisseau, G. Tarran, N. Van Oostende, L. Zhao, S. Zheng, Interoperable vocabulary for marine microbial flow cytometry. Front. Mar. Sci. 9, 975877 (2022).

55. S. Sunagawa, L. P. Coelho, S. Chaffron, J. R. Kultima, K. Labadie, G. Salazar, B. Djahanschiri, G. Zeller, D. R. Mende, A. Alberti, F. M. Cornejo-Castillo, P. I. Costea, C. Cruaud, F. d’Ovidio, S. Engelen, I. Ferrera, J. M. Gasol, L. Guidi, F. Hildebrand, F. Kokoszka, C. Lepoivre, G. Lima-Mendez, J. Poulain, B. T. Poulos, M. Royo-Llonch, H. Sarmento, S. Vieira-Silva, C. Dimier, M. Picheral, S. Searson, S. Kandels-Lewis, Tara Oceans coordinators, C. Bowler, C. De Vargas, G. Gorsky, N. Grimsley, P. Hingamp, D. Iudicone, O. Jaillon, F. Not, H. Ogata, S. Pesant, S. Speich, L. Stemmann, M. B. Sullivan, J. Weissenbach, P. Wincker, E. Karsenti, J. Raes, S. G. Acinas, P. Bork, E. Boss, C. Bowler, M. Follows, L. Karp-Boss, U. Krzic, E. G. Reynaud, C. Sardet, M. Sieracki, D. Velayoudon, Structure and function of the global ocean microbiome. Science 348, 1261359 (2015).

56. F. M. Ibarbalz, N. Henry, M. C. Brandão, S. Martini, G. Busseni, H. Byrne, L. P. Coelho, H. Endo, J. M. Gasol, A. C. Gregory, F. Mahé, J. Rigonato, M. Royo-Llonch, G. Salazar, I. Sanz-Sáez, E. Scalco, D. Soviadan, A. A. Zayed, A. Zingone, K. Labadie, J. Ferland, C. Marec, S. Kandels, M. Picheral, C. Dimier, J. Poulain, S. Pisarev, M. Carmichael, S. Pesant, M. Babin, E. Boss, D. Iudicone, O. Jaillon, S. G. Acinas, H. Ogata, E. Pelletier, L. Stemmann, M. B. Sullivan, S. Sunagawa, L. Bopp, C. De Vargas, L. Karp-Boss, P. Wincker, F. Lombard, C. Bowler, L. Zinger, S. G. Acinas, M. Babin, P. Bork, E. Boss, C. Bowler, G. Cochrane, C. De Vargas, M. Follows, G. Gorsky, N. Grimsley, L. Guidi, P. Hingamp, D. Iudicone, O. Jaillon, S. Kandels, L. Karp-Boss, E. Karsenti, F. Not, H. Ogata, S. Pesant, N. Poulton, J. Raes, C. Sardet, S. Speich, L. Stemmann, M. B. Sullivan, S. Sunagawa, P. Wincker, Global Trends in Marine Plankton Diversity across Kingdoms of Life. Cell 179, 1084–1097.e21 (2019).

57. G. Lima-Mendez, K. Faust, N. Henry, J. Decelle, S. Colin, F. Carcillo, S. Chaffron, J. C. Ignacio-Espinosa, S. Roux, F. Vincent, L. Bittner, Y. Darzi, J. Wang, S. Audic, L. Berline, G. Bontempi, A. M. Cabello, L. Coppola, F. M. Cornejo-Castillo, F. d’Ovidio, L. De Meester, I. Ferrera, M.-J. Garet-Delmas, L. Guidi, E. Lara, S. Pesant, M. Royo-Llonch, G. Salazar, P. Sánchez, M. Sebastian, C. Souffreau, C. Dimier, M. Picheral, S. Searson, S. Kandels-Lewis, Tara Oceans coordinators, G. Gorsky, F. Not, H. Ogata, S. Speich, L. Stemmann, J. Weissenbach, P. Wincker, S. G. Acinas, S. Sunagawa, P. Bork, M. B. Sullivan, E. Karsenti, C. Bowler, C. De Vargas, J. Raes, Determinants of community structure in the global plankton interactome. Science 348, 1262073 (2015).

58. A. Ott, M. Quintela-Baluja, A. M. Zealand, G. O’Donnell, M. R. M. Haniffah, D. W. Graham, Improved quantitative microbiome profiling for environmental antibiotic resistance surveillance. Environmental Microbiome 16, 21 (2021).

59. J. P. Pettersen, M. S. Gundersen, E. Almaas, Robust bacterial co-occurence community structures are independent of r-and K-selection history. Sci Rep 11, 23497 (2021).

60. J. H. Poelen, J. D. Simons, C. J. Mungall, Global biotic interactions: An open infrastructure to share and analyze species-interaction datasets. Ecological Informatics 24, 148–159 (2014).

61. T. Nakayama, M. Nomura, Y. Takano, G. Tanifuji, K. Shiba, K. Inaba, Y. Inagaki, M. Kawata, Single-cell genomics unveiled a cryptic cyanobacterial lineage with a worldwide distribution hidden by a dinoflagellate host. Proc. Natl. Acad. Sci. U.S.A. 116, 15973–15978 (2019).

62. F. Partensky, W. R. Hess, D. Vaulot, *Prochlorococcus*, a Marine Photosynthetic Prokaryote of Global Significance. Microbiol Mol Biol Rev 63, 106–127 (1999).

63. P. Flombaum, J. L. Gallegos, R. A. Gordillo, J. Rincón, L. L. Zabala, N. Jiao, D. M. Karl, W. K. W. Li, M. W. Lomas, D. Veneziano, C. S. Vera, J. A. Vrugt, A. C. Martiny, Present and future global distributions of the marine Cyanobacteria *Prochlorococcus* and *Synechococcus*. Proc. Natl. Acad. Sci. U.S.A. 110, 9824–9829 (2013).

64. K. Leblanc, J. Arístegui, L. Armand, P. Assmy, B. Beker, A. Bode, E. Breton, V. Cornet, J. Gibson, M.P. Gosselin, E. Kopczynska, H. Marshall, J. Peloquin, S. Piontkovski, A. J. Poulton, B. Quéguiner, R. Schiebel, R. Shipe, J. Stefels, M. A. Van Leeuwe, M. Varela, C. Widdicombe, M. Yallop, A global diatom database – abundance, biovolume and biomass in the world ocean. *Earth Syst*. Sci. Data 4, 149– 165 (2012).

65. E. Villarino, J. R. Watson, B. Jönsson, J. M. Gasol, G. Salazar, S. G. Acinas, M. Estrada, R. Massana, R. Logares, C. R. Giner, M. C. Pernice, M. P. Olivar, L. Citores, J. Corell, N. Rodríguez-Ezpeleta, J. L. Acuña, A. Molina-Ramírez, J. I. González-Gordillo, A. Cózar, E. Martí, J. A. Cuesta, S. Agustí, E. Fraile-Nuez, C. M. Duarte, X. Irigoien, G. Chust, Large-scale ocean connectivity and planktonic body size. Nat Commun 9, 142 (2018).

66. R. Massana, Eukaryotic Picoplankton in Surface Oceans. Annu. Rev. Microbiol. 65, 91–110 (2011).

67. R. Massana, R. Logares, Eukaryotic versus prokaryotic marine picoplankton ecology. Environmental Microbiology 15, 1254–1261 (2013).

68. A. J. Irwin, A. M. Nelles, Z. V. Finkel, Phytoplankton niches estimated from field data. Limnology & Oceanography 57, 787–797 (2012).

69. P. Frémont, M. Gehlen, M. Vrac, J. Leconte, T. O. Delmont, P. Wincker, D. Iudicone, O. Jaillon, Restructuring of plankton genomic biogeography in the surface ocean under climate change. Nat. Clim. Chang. 12, 393–401 (2022).

70. M. J. Follows, S. Dutkiewicz, Modeling Diverse Communities of Marine Microbes. Annu. Rev. Mar. Sci. 3, 427–451 (2011).

71. D. Righetti, M. Vogt, N. Gruber, A. Psomas, N. E. Zimmermann, Global pattern of phytoplankton diversity driven by temperature and environmental variability. Sci. Adv. 5, eaau6253 (2019).

72. T. Soulié, F. Vidussi, S. Mas, B. Mostajir, Functional Stability of a Coastal Mediterranean Plankton Community During an Experimental Marine Heatwave. Front. Mar. Sci. 9, 831496 (2022).

73. H. A. Bouman, O. Ulloa, D. J. Scanlan, K. Zwirglmaier, W. K. W. Li, T. Platt, V. Stuart, R. Barlow, O. Leth, L. Clementson, V. Lutz, M. Fukasawa, S. Watanabe, S. Sathyendranath, Oceanographic Basis of the Global Surface Distribution of *Prochlorococcus* Ecotypes. Science 312, 918–921 (2006).

74. F. Not, M. Latasa, R. Scharek, M. Viprey, P. Karleskind, V. Balagué, I. Ontoria-Oviedo, A. Cumino, E. Goetze, D. Vaulot, R. Massana, Protistan assemblages across the Indian Ocean, with a specific emphasis on the picoeukaryotes. Deep Sea Research Part I: Oceanographic Research Papers 55, 1456–1473 (2008).

75. E. T. Buitenhuis, M. Vogt, R. Moriarty, N. Bednaršek, S. C. Doney, K. Leblanc, C. Le Quéré, Y.-W. Luo, C. O’Brien, T. O’Brien, J. Peloquin, R. Schiebel, C. Swan, MAREDAT: towards a world atlas of MARine Ecosystem DATa. *Earth Syst*. Sci. Data 5, 227–239 (2013).

76. Y.-W. Luo, S. C. Doney, L. A. Anderson, M. Benavides, I. Berman-Frank, A. Bode, S. Bonnet, K. H. Boström, D. Böttjer, D. G. Capone, E. J. Carpenter, Y. L. Chen, M. J. Church, J. E. Dore, L. I. Falcón, A. Fernández, R. A. Foster, K. Furuya, F. Gómez, K. Gundersen, A. M. Hynes, D. M. Karl, S. Kitajima, R. J. Langlois, J. LaRoche, R. M. Letelier, E. Marañón, D. J. McGillicuddy, P. H. Moisander, C. M. Moore, B. Mouriño-Carballido, M. R. Mulholland, J. A. Needoba, K. M. Orcutt, A. J. Poulton, E. Rahav, P. Raimbault, A. P. Rees, L. Riemann, T. Shiozaki, A. Subramaniam, T. Tyrrell, K. A. Turk-Kubo, M. Varela, T. A. Villareal, E. A. Webb, A. E. White, J. Wu, J. P. Zehr, Database of diazotrophs in global ocean: abundance, biomass and nitrogen fixation rates. *Earth Syst*. Sci. Data 4, 47–73 (2012).

77. D. G. Capone, J. P. Zehr, H. W. Paerl, B. Bergman, E. J. Carpenter, *Trichodesmium*, a Globally Significant Marine Cyanobacterium. Science 276, 1221–1229 (1997).

78. D. Karl, R. Letelier, Nitrogen fixation-enhanced carbon sequestration in low nitrate, low chlorophyll seascapes. Mar. Ecol. Prog. Ser. 364, 257–268 (2008).

79. R. L. Mather, S. E. Reynolds, G. A. Wolff, R. G. Williams, S. Torres-Valdes, E. M. S. Woodward, A. Landolfi, X. Pan, R. Sanders, E. P. Achterberg, Phosphorus cycling in the North and South Atlantic Ocean subtropical gyres. Nature Geosci 1, 439–443 (2008).

80. C. M. Moore, M. M. Mills, K. R. Arrigo, I. Berman-Frank, L. Bopp, P. W. Boyd, E. D. Galbraith, R. J. Geider, C. Guieu, S. L. Jaccard, T. D. Jickells, J. La Roche, T. M. Lenton, N. M. Mahowald, E. Marañón, I. Marinov, J. K. Moore, T. Nakatsuka, A. Oschlies, M. A. Saito, T. F. Thingstad, A. Tsuda, O. Ulloa, Processes and patterns of oceanic nutrient limitation. Nature Geosci 6, 701–710 (2013).

81. B. Rost, U. Riebesell, “Coccolithophores and the biological pump: responses to environmental changes” in *Coccolithophores*, H. R. Thierstein, J. R. Young, Eds. (Springer Berlin Heidelberg, Berlin, Heidelberg, 2004; http://link.springer.com/10.1007/978-3-662-06278-4_5), pp. 99–125.

82. H. Liu, I. Probert, J. Uitz, H. Claustre, S. Aris-Brosou, M. Frada, F. Not, C. De Vargas, Extreme diversity in noncalcifying haptophytes explains a major pigment paradox in open oceans. Proc. Natl. Acad. Sci. U.S.A. 106, 12803–12808 (2009).

83. S. Dutkiewicz, P. Cermeno, O. Jahn, M. J. Follows, A. E. Hickman, D. A. A. Taniguchi, B. A. Ward, Dimensions of marine phytoplankton diversity. Biogeosciences 17, 609–634 (2020).

84. R. M. Wright, C. Le Quéré, E. Buitenhuis, S. Pitois, M. J. Gibbons, Role of jellyfish in the plankton ecosystem revealed using a global ocean biogeochemical model. Biogeosciences 18, 1291–1320 (2021).

85. S. M. Vallina, M. J. Follows, S. Dutkiewicz, J. M. Montoya, P. Cermeno, M. Loreau, Global relationship between phytoplankton diversity and productivity in the ocean. Nat Commun 5, 4299 (2014).

86. L. Bopp, O. Aumont, P. Cadule, S. Alvain, M. Gehlen, Response of diatoms distribution to global warming and potential implications: A global model study. Geophysical Research Letters 32, 2005GL023653 (2005).

87. T. O. Delmont, J. J. Pierella Karlusich, I. Veseli, J. Fuessel, A. M. Eren, R. A. Foster, C. Bowler, P. Wincker, E. Pelletier, Heterotrophic bacterial diazotrophs are more abundant than their cyanobacterial counterparts in metagenomes covering most of the sunlit ocean. The ISME Journal 16, 927–936 (2022).

88. M. R. McLaren, A. D. Willis, B. J. Callahan, Consistent and correctable bias in metagenomic sequencing experiments. eLife 8, e46923 (2019).

89. M. R. McLaren, J. T. Nearing, A. D. Willis, K. G. Lloyd, B. J. Callahan, Implications of taxonomic bias for microbial differential-abundance analysis. [Preprint] (2022). 10.1101/2022.08.19.504330.

90. S. Colin, L. P. Coelho, S. Sunagawa, C. Bowler, E. Karsenti, P. Bork, R. Pepperkok, C. De Vargas, Quantitative 3D-imaging for cell biology and ecology of environmental microbial eukaryotes. eLife 6, e26066 (2017).

91. J. J. P. Karlusich, F. M. Ibarbalz, C. Bowler, Phytoplankton in the *Tara* Ocean. Annu. Rev. Mar. Sci. 12, 233–265 (2020).

92. G. Salazar, L. Paoli, A. Alberti, J. Huerta-Cepas, H.-J. Ruscheweyh, M. Cuenca, C. M. Field, L. P. Coelho, C. Cruaud, S. Engelen, A. C. Gregory, K. Labadie, C. Marec, E. Pelletier, M. Royo-Llonch, S. Roux, P. Sánchez, H. Uehara, A. A. Zayed, G. Zeller, M. Carmichael, C. Dimier, J. Ferland, S. Kandels, M. Picheral, S. Pisarev, J. Poulain, S. G. Acinas, M. Babin, P. Bork, C. Bowler, C. De Vargas, L. Guidi, P. Hingamp, D. Iudicone, L. Karp-Boss, E. Karsenti, H. Ogata, S. Pesant, S. Speich, M. B. Sullivan, P. Wincker, S. Sunagawa, S. G. Acinas, M. Babin, P. Bork, E. Boss, C. Bowler, G. Cochrane, C. De Vargas, M. Follows, G. Gorsky, N. Grimsley, L. Guidi, P. Hingamp, D. Iudicone, O. Jaillon, S. Kandels-Lewis, L. Karp-Boss, E. Karsenti, F. Not, H. Ogata, S. Pesant, N. Poulton, J. Raes, C. Sardet, S. Speich, L. Stemmann, M. B. Sullivan, S. Sunagawa, P. Wincker, Gene Expression Changes and Community Turnover Differentially Shape the Global Ocean Metatranscriptome. Cell 179, 1068–1083.e21 (2019).

93. T. Cavalier-Smith, “The Evolution of genome size” (1985; https://api.semanticscholar.org/CorpusID:86524579).

94. T. R. Gregory, A BIRD’S-EYE VIEW OF THE C-VALUE ENIGMA: GENOME SIZE, CELL SIZE, AND METABOLIC RATE IN THE CLASS AVES. Evolution 56, 121–130 (2002).

95. T. Cavalier-Smith, Economy, Speed and Size Matter: Evolutionary Forces Driving Nuclear Genome Miniaturization and Expansion. Annals of Botany 95, 147–175 (2005).

96. J. Beardall, D. Allen, J. Bragg, Z. V. Finkel, K. J. Flynn, A. Quigg, T. A. V. Rees, A. Richardson, J. A. Raven, Allometry and stoichiometry of unicellular, colonial and multicellular phytoplankton. New Phytologist 181, 295–309 (2009).

97. L. Gonzalez-de-Salceda, F. Garcia-Pichel, The allometry of cellular DNA and ribosomal gene content among microbes and its use for the assessment of microbiome community structure. Microbiome 9, 173 (2021).

98. M. Luo, Y. Ji, D. Warton, D. W. Yu, Extracting abundance information from DNA -based data. Molecular Ecology Resources 23, 174–189 (2023).

99. A. Houdan, “CYCLE BIOLOGIQUE ET STRATEGIES DE DEVELOPPEMENT CHEZ LES COCCOLITHOPHORES (PRYMNESIOPHYCEAE, HAPTOPHYTA). IMPLICATIONS ECOLOGIQUES,” (2003).

100. H. Li, R. Durbin, Fast and accurate short read alignment with Burrows–Wheeler transform. Bioinformatics 25, 1754–1760 (2009).

101. J. Weissenbach, A. Aguilera, L. Bas Conn, J. Pinhassi, C. Legrand, H. Farnelid, Ploidy levels in diverse picocyanobacteria from the Baltic Sea. Environ Microbiol Rep 16 (2024).

102. M. Griese, C. Lange, J. Soppa, Ploidy in cyanobacteria. FEMS Microbiol Lett 323, 124–131 (2011).

103. J. Tackmann, J. F. Matias Rodrigues, C. Von Mering, Rapid Inference of Direct Interactions in Large-Scale Ecological Networks from Heterogeneous Microbial Sequencing Data. Cell Systems 9, 286–296.e8 (2019).

104. S. Chaffron, E. Delage, M. Budinich, D. Vintache, N. Henry, C. Nef, M. Ardyna, A. A. Zayed, P. C. Junger, P. E. Galand, C. Lovejoy, A. E. Murray, H. Sarmento, Tara Oceans coordinators, S. G. Acinas, M. Babin, D. Iudicone, O. Jaillon, E. Karsenti, P. Wincker, L. Karp-Boss, M. B. Sullivan, C. Bowler, C. De Vargas, D. Eveillard, Environmental vulnerability of the global ocean epipelagic plankton community interactome. Sci. Adv. 7, eabg1921 (2021).

105. G. Gorsky, M. D. Ohman, M. Picheral, S. Gasparini, L. Stemmann, J.-B. Romagnan, A. Cawood, S. Pesant, C. Garcia-Comas, F. Prejger, Digital zooplankton image analysis using the ZooScan integrated system. Journal of Plankton Research 32, 285–303 (2010).

106. M. Picheral, L. Guidi, L. Stemmann, D. M. Karl, G. Iddaoud, G. Gorsky, The Underwater Vision Profiler 5: An advanced instrument for high spatial resolution studies of particle size spectra and zooplankton. Limnology and Oceanography: Methods 8, 462–473 (2010).

107. C. Sieracki, M. Sieracki, C. Yentsch, An imaging-in-flow system for automated analysis of marine microplankton. Mar. Ecol. Prog. Ser. 168, 285–296 (1998).

108. Z. Mériguet, G. Bourdin, L. Jalabert, O. Bun, L. Caray--Counil, A. Elineau, G. Gorsky, F. Lombard, Global scale surface micro-plankton dataset collected with Deck Net and imaged with FlowCam during the Tara Pacific Expedition, SEANOE (2024); 10.17882/102697.

109. Z. Mériguet, G. Bourdin, L. Jalabert, O. Bun, L. Caray--Counil, A. Elineau, G. Gorsky, F. Lombard, Global scale surface meso-plankton dataset collected with High-Speed Net and imaged with ZooScan during the Tara Pacific Expedition, SEANOE (2024); 10.17882/102336.

110. S. Menden-Deuer, E. J. Lessard, Carbon to volume relationships for dinoflagellates, diatoms, and other protist plankton. Limnology and Oceanography 45, 569–579 (2000).

111. P. Legendre, L. Legendre, P. Legendre, Numerical Ecology (Elsevier, Amsterdam Boston, 3rd English ed (Online-Ausg.)., 2012)*Developments in environmental modelling*.

112. H. E. Garcia, T. P. Boyer, O. K. Baranova, World Ocean Atlas 2018: Product Documentation. doi: 10.25923/TZYW-RP36 (2019).

113. S. Sathyendranath, R. Brewin, C. Brockmann, V. Brotas, B. Calton, A. Chuprin, P. Cipollini, A. Couto, J. Dingle, R. Doerffer, C. Donlon, M. Dowell, A. Farman, M. Grant, S. Groom, A. Horseman, T. Jackson, H. Krasemann, S. Lavender, V. Martinez-Vicente, C. Mazeran, F. Mélin, T. Moore, D. Müller, P. Regner, S. Roy, C. Steele, F. Steinmetz, J. Swinton, M. Taberner, A. Thompson, A. Valente, M. Zühlke, V. Brando, H. Feng, G. Feldman, B. Franz, R. Frouin, R. Gould, S. Hooker, M. Kahru, S. Kratzer, B. Mitchell, F. Muller-Karger, H. Sosik, K. Voss, J. Werdell, T. Platt, An Ocean-Colour Time Series for Use in Climate Studies: The Experience of the Ocean-Colour Climate Change Initiative (OC-CCI). Sensors 19, 4285 (2019).

114. L. Breiman, Bagging predictors. Mach Learn 24, 123–140 (1996).

115. J. Snoek, H. Larochelle, R. P. Adams, Practical Bayesian Optimization of Machine Learning Algorithms. arXiv [Preprint] (2012). 10.48550/ARXIV.1206.2944.

116. D. R. Cutler, T. C. Edwards, K. H. Beard, A. Cutler, K. T. Hess, J. Gibson, J. J. Lawler, RANDOM FORESTS FOR CLASSIFICATION IN ECOLOGY. Ecology 88, 2783–2792 (2007).

117. P. C. Mahalanobis, On the generalized distance in statistics. Proceedings of the National Institute of Sciences (Calcutta) 2, 49–55 (1936).

118. D. S. Wilks, Statistical Methods in the Atmospheric Sciences (Elsevier, Amsterdam Paris, 2nd ed., 2006)International geophysics series.

119. B. Efron, R. J. Tibshirani, *An Introduction to the Bootstrap* (Chapman and Hall/CRC, ed. 0, 1994; https://www.taylorfrancis.com/books/9781000064988).

120. B. Bergman, G. Sandh, S. Lin, J. Larsson, E. J. Carpenter, *Trichodesmium* – a widespread marine cyanobacterium with unusual nitrogen fixation properties. FEMS Microbiol Rev 37, 286–302 (2013).

121. C. Cesar-Ribeiro, C. S. Barbosa, V. Terra, N. D. C. Ghisi, Prochlorococcus and Synechococcus marine cyanobacteria: a scientometrics review. Lat. Am. J. Aquat. Res. 51, 556–569 (2023).

122. C. M. Brown, J. E. Lawrence, D. A. Campbell, Are phytoplankton population density maxima predictable through analysis of host and viral genomic DNA content? J. Mar. Biol. Ass. 86, 491–498 (2006).

123. I. S. Hamza, I. Biegala, A. B. Zouari, F. Akrout, F. A. Keskes, A. Hamza, M. B. Hassen, Diversity and abundance of diazotrophic cyanobacteria in the central coastal area of the Gulf of Gabès (South-eastern Tunisia). Regional Studies in Marine Science 42, 101653 (2021).

124. M. Vogt, C. O’Brien, J. Peloquin, V. Schoemann, E. Breton, M. Estrada, J. Gibson, D. Karentz, M. A. Van Leeuwe, J. Stefels, C. Widdicombe, L. Peperzak, Global marine plankton functional type biomass distributions: *Phaeocystis* spp. *Earth Syst*. Sci. Data 4, 107–120 (2012).

125. Roy, S. Sathyendranath, T. Platt, Retrieval of phytoplankton size from bio-optical measurements: theory and applications. J. R. Soc. Interface. 8, 650–660 (2011).

